# Single-cell-level response to drought in *Sorghum bicolor* reveals novel targets for improving water use efficiency

**DOI:** 10.1101/2025.08.28.671794

**Authors:** Matt Stata, Sharon Greenblum, Purva Karia, Maxim Koriabine, Yuko Yoshinaga, Keykhosrow Keymanesh, Cheng Zhao, Ronan C. O’Malley, Seung Y. Rhee

## Abstract

Increasing drought threatens global agriculture, especially in water-limited regions. *Sorghum bicolor*, a drought-tolerant C_4_ grass, is a promising bioenergy crop for cultivation on marginal lands, yet its molecular drought responses remain poorly understood. To uncover single-cell-level transcriptional responses to drought, we performed single-nucleus RNA sequencing on mature sorghum leaves under well-watered and drought conditions. We identified major cell types and analyzed differential gene expression across mesophyll, bundle sheath, epidermal, vascular, and stomatal cells. Surprisingly, drought effects on transcriptomes exceeded differences due to cell identity, revealing a shared response across cell types. We leveraged this convergence to identify candidate regulators of drought-responsive gene expression. These findings advance our understanding of sorghum drought adaptation and offer new targets for engineering enhanced water use efficiency in bioenergy crops.

## Introduction

*Sorghum bicolor* is a drought-resilient C_4_ cereal crop critical for food and fodder production. As the world’s fifth most important cereal, it feeds over 500 million people in Africa and Asia (Paterson *et al.*, 2009; Mace *et al.*, 2013; Ameen *et al.*, 2024). Its success on marginal lands is linked to traits such as a deep root system, waxy leaf cuticle, and efficient stomatal regulation (Lamb *et al.*, 2021; Al-Salman *et al.*, 2024). These features not only support food security but also position sorghum as a compelling candidate for sustainable bioenergy production (Lamb *et al.*, 2021).

Sorghum uses the C_4_ photosynthetic pathway, which enhances carbon assimilation under high-temperature and water-limited conditions, contributing to its superior water use efficiency (WUE) relative to other economically important crops (Ghannoum *et al.*, 2011; Schittenhelm & Schroetter, 2014; Vadez *et al.*, 2021). Coupled with high biomass accumulation and publicly available genomic resources, sorghum presents a promising system for studying and improving drought tolerance in cereal crops (Paterson *et al.*, 2009; McCormick *et al.*, 2018). Despite its natural resilience, molecular breeding to further enhance sorghum’s drought performance has lagged, in part due to an incomplete understanding of the genes and regulatory networks underlying its stress adaptation.

Although several physiological traits linked to drought tolerance have been characterized -such as the stay-green phenotype, osmotic adjustment, and tight stomatal control - much of the underlying genetic variation remains unexploited in modern breeding (Harris *et al.*, 2007; Kassahun *et al.*, 2010; Thomas & Ougham, 2014; Johnson *et al.*, 2015; Goche *et al.*, 2020; Zhang *et al.*, 2024). Bulk-tissue transcriptomic studies identified candidate genes and stress-induced pathways, including abscisic acid (ABA) signaling, reactive oxygen species (ROS) detoxification, and delayed senescence (Varoquaux *et al.*, 2019). However, bulk transcriptomic approaches obscure responses at the individual cell and tissue levels, limiting the ability to pinpoint spatially regulated drought-adaptive mechanisms (Berrío *et al.,* 2025; Guo *et al.,* 2025; Liu *et al.,* 2022).

Emerging single-cell and spatial transcriptomic methods offer an opportunity to resolve these limitations by capturing gene expression at the level of individual cells. In C_4_ species like sorghum, where photosynthesis requires coordination between distinct cell types in the leaf, this resolution is especially valuable. Here, we apply single-nucleus RNA sequencing (snRNA-seq) to generate a high-resolution atlas of drought-responsive gene expression across leaf cell types in sorghum. This approach enables the identification of shared and cell-specific responses, revealing candidate regulators of drought adaptation that can inform crop improvement strategies.

## Results

### A sorghum leaf cell atlas

To understand how *Sorghum bicolor* leaves respond to drought at the cellular level, we conducted single-nucleus RNAseq on the most recent fully-expanded mature leaves of plants that were either well-watered or water-limited (< 40% soil water content) at two timepoints, six and ten days into drought treatment **(Fig. S1)**. To ensure robust cell-type identification across experimental conditions, we clustered control and drought cells separately and cross-validated their identities before uniform manifold approximation and projection (UMAP) visualization. This approach revealed largely congruent clusters corresponding to the major leaf cell types - mesophyll, bundle sheath, pavement, phloem parenchyma and other vascular parenchyma **(Figs. 1A, S2-4)**. The remaining two clusters represent mixed cell-type groups: one containing companion cells with sieve elements and the other a mix of subsidiary and guard cells, neither of which could be cleanly separated by marker genes or more granular clustering. The final UMAP of 25,794 control and 29,234 drought-treated, filtered cells comprises five discrete cell-type clusters alongside these two mixed-cell clusters, demonstrating robust cell-type identification across conditions.

**Fig. 1.**
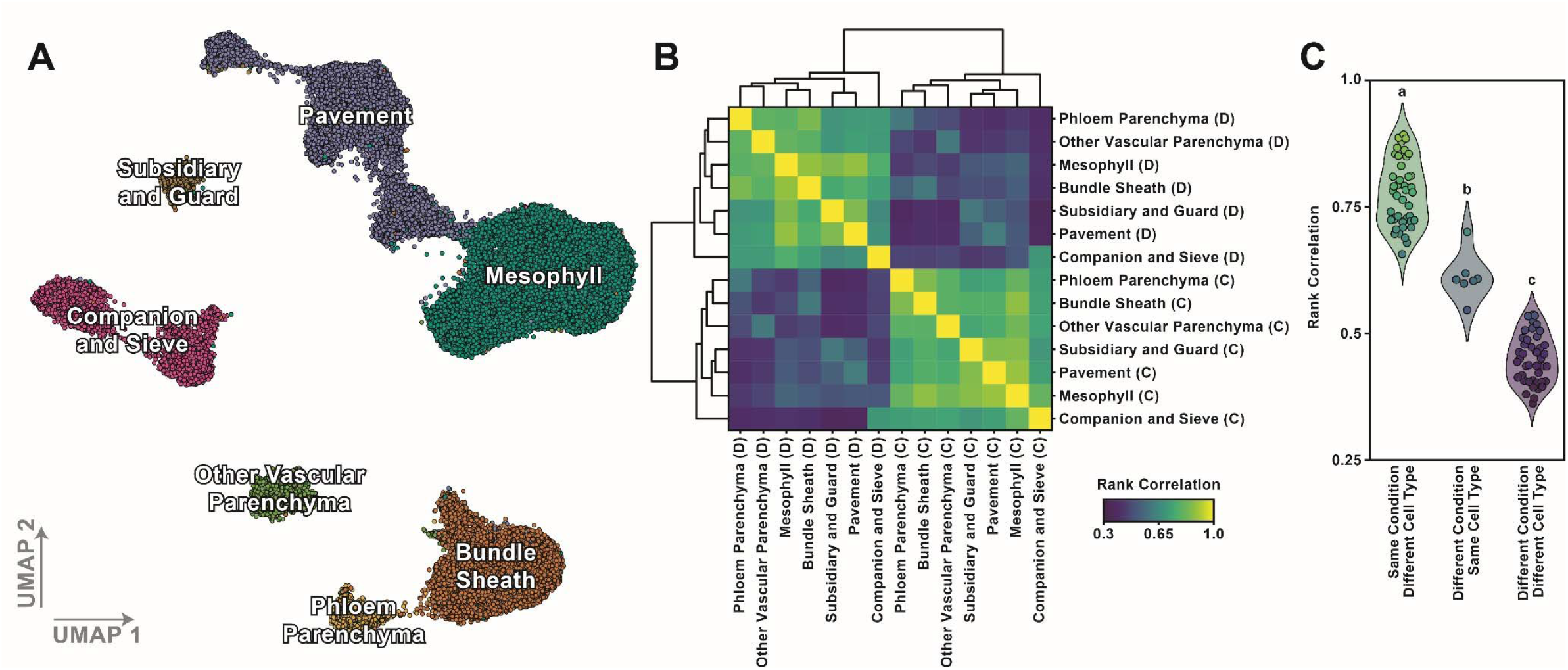
Cell identities and response to drought. A, UMAP of cell identities from eight total snRNA-seq libraries. Two replicates each for control and drought treated sorghum leaves from six and ten days into drought treatment were integrated. Sorghum orthologs of cell type marker genes from the literature were used to determine cell identities as outlined in the Supplementary Information. B, Clustered correlation heatmap plot showing Spearman rank correlations among control and drought cell types based genes with expression over 10 TPM in at least one cell type. Control and drought treatments are denoted with letters C and D, respectively. C, Violin plot showing three categories of pairwise rank correlation comparisons from (B) based on samples from different cell types (n = 42), experimental conditions (n = 7), or both (n = 42). Letters denote statistical groups determined by one-way ANOVA and Tukey HSD post-hoc tests, and marker and violin colors correspond to the color scale in b, using the mean for violin colors.

### Drought response supersedes intrinsic cell identity

We next asked how cell-type specific expression profiles differed across cell types and water availability. Correlation analysis of pseudobulk transcripts-per-million (TPM) expression profiles under control and drought conditions showed that water availability, not cell identity, is the dominant source of transcriptomic variation, with cells clustering by condition rather than by cell type (**Fig. 1B**). Different cell types under the same water availability had the highest correlation (0.78 ± 0.07), followed by cells of the same type across treatments (0.61 ± 0.05), which were still more correlated than cells differing in both condition and cell type (0.45 ± 0.05); **Fig. 1C**).

Consistently, principal component analysis (PCA) showed that transcriptome variation associated with drought response was primarily captured by the first principal component (PC1, 27.2% of the variation), while differences associated with cell identity were represented along secondary components, PCs 2-4 (**Fig. 2A**). The largest separation within PC2 was between photosynthetic (mesophyll, bundle sheath) and non-photosynthetic cell types. In addition to PC2 (12.9% of the variation), PCs 3 and 4 (11.2% and 8.9% of the variation) recovered further intrinsic cell-type distinctions regardless of treatment (**Fig. 2B**). Using the pseudobulk expression data, we next defined two sets of genes, those whose abundances were increased or decreased by drought treatment in at least one cell type, respectively. When performing PCA using only the 1,679 drought-induced differentially expressed genes (DEGs) (p_adj_ < 0.05), cell types responded similarly, with differences between cell types persisting or even increasing (**Fig. 2C**). In contrast, PCA on the 2,233 drought-reduced DEGs converged all cell types toward a common state (**Fig. 2D**). Drought-induced DEGs therefore preserve cell identities while drought-downregulated DEGs have a homogenizing effect on cell-type specific gene expression, indicative of down-regulation of cell-specific functions. Important functions enriched among drought-induced genes include vesicle-mediated transport, intracellular protein transport, and small GTPase-mediated signal transduction, while drought-reduced genes are primarily related to photosynthesis and growth **(Fig. 2E)**.

**Fig. 2.**
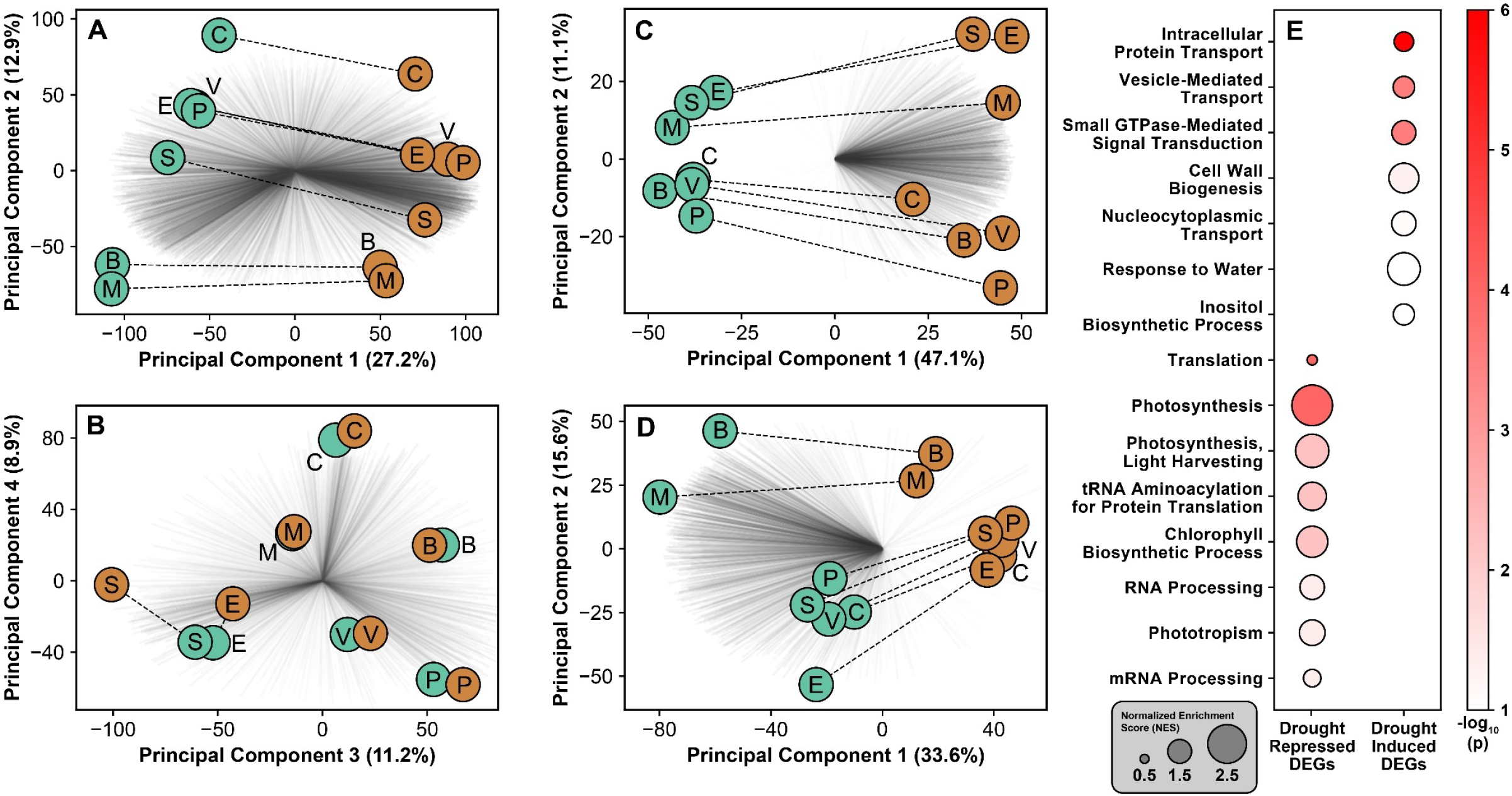
Drivers of cell-type transcriptome patterns. A-B, principal component analysis showing that component 1 (A) primarily reflects differences in response to drought while components 2-4 (A, B) primarily reflect differences in cell identity regardless of drought. Gene vectors are shaded by combined PC loadings and scaled to match the plot area for visibility. C-D, principal component analysis on drought-induced differentially expressed genes (DEGs) (C) and drought-reduced DEGs (D), with scaled gene vectors shaded by combined weighted loadings for PC1 and PC2. (E), gene-set enrichment analysis (GSEA) bubble plot for drought-induced and drought-reduced DEGs. Bubble size and color denote normalized enrichment score (NES) and -log_10_(p) values, respectively. The Kolmogorov-Smirnov statistic and “elim” algorithm in TopGO were used with genes ranked by the best DESeq2 Wald test statistic across cell types. GO enrichment for the drivers of PC1 and PC2 are presented in Figure S5. Letters denote cell type: B, bundle sheath cells; C, companion cells and sieve elements; E, epidermal pavement cells; M, mesophyll cells; P, phloem parenchyma cells; S, subsidiary and guard cells; V, other vascular parenchyma cells.

Given the observed prominence of drought response in driving the transcriptome variation across cell types, we next asked what functions were enriched among the genes driving PC1 in the PCA based on all expressed genes. PC1-driver genes that were positively associated with droughted samples were enriched in translation, vesicle-mediated transport, intracellular protein transport, and small GTPase-mediated signal transduction, consistent with drought-induced genes (**Figs. 2E, S5**). PC1-driver genes that were associated with control samples were enriched in similar GO processes as drought-repressed DEGs such as photosynthesis, light harvesting, chlorophyll biosynthesis, and phototropism (**Figs. 2E, S5**). These results further confirm that drought response is the primary driver of variability in the transcriptome across cell types. GO terms enriched among the drivers of PC2 primarily separate photosynthetic from non-photosynthetic cell types, with photosynthesis and light harvesting enriched in genes driving the separation of mesophyll and bundle sheath cells from non-photosynthetic cells, and vesicle mediated transport, TCA cycle, phosphorelay signal transduction, and intracellular protein transport driving the separation in the opposite direction (**Fig. S5**). Overall, our PCA and GO enrichment results indicate a surprisingly similar transcriptomic response to drought across cell types coupled with an expected drought-induced shutdown of processes related to photosynthesis and growth.

### Cell-type specific responses to drought

We next asked whether cell-type specific changes in gene expression in response to drought could be identified despite the largely uniform transcriptomic reprogramming. We compared the extended tau cell-specificity metric (Yanai et al., 2005; Lüleci & Yılmaz, 2022) between control and drought conditions, to ask whether any genes gained or lost cell-type specificity between treatments (**Fig. 3A-B**). This analysis uncovered 299 genes whose cell-specific transcript abundance is reduced by drought, and 355 genes with cell-specific increase in transcript abundance (**Fig. 3A-B**). For example, pavement cells induced several cellulose synthases and a synaptotagmin-3 ortholog (Sobic.010G236100), which is an endoplasmic reticulum-plasma membrane tether that maintains membrane integrity and lipid homeostasis under abiotic stress (Ruiz-Lopez *et al.*, 2021; García-Hernández *et al.*, 2025A; Garcia-Hernandez *et al.*, 2025B) (**Fig. 3C**). Companion and sieve elements induced a hexokinase 1 (HXK1) ortholog (Sobic.003G291800) which functions as a glucose sensor to integrate nutrient, light and hormone signals (Mu *et al.*, 2024; Rawat *et al.*, 2024), while non-phloem vascular parenchyma specifically increased a 2-oxoglutarate/Fe(II)-dependent oxygenase (Sobic.001G314300), whose Arabidopsis orthologs have salicylic acid (SA) hydroxylase activity that inactivates SA (Falcone Ferreyra *et al.*, 2015; Zhang *et al.*, 2017) (**Fig. 3D-E**). Bundle sheath cells selectively reduced a fatty acid desaturase 2 (FAD2) ortholog (Sobic.004G260800), which provides polyunsaturated fatty acids to ER-derived membranes (Launhardt *et al.*, 2024; **Fig. 3F**). These examples of cell-type-specific shifts in expression spanning functions such as cell-wall remodeling, lipid and sugar signaling may reflect fine-tuned adaptation in different cell types under water deficit even while most drought response is conserved across cell types.

**Fig. 3.**
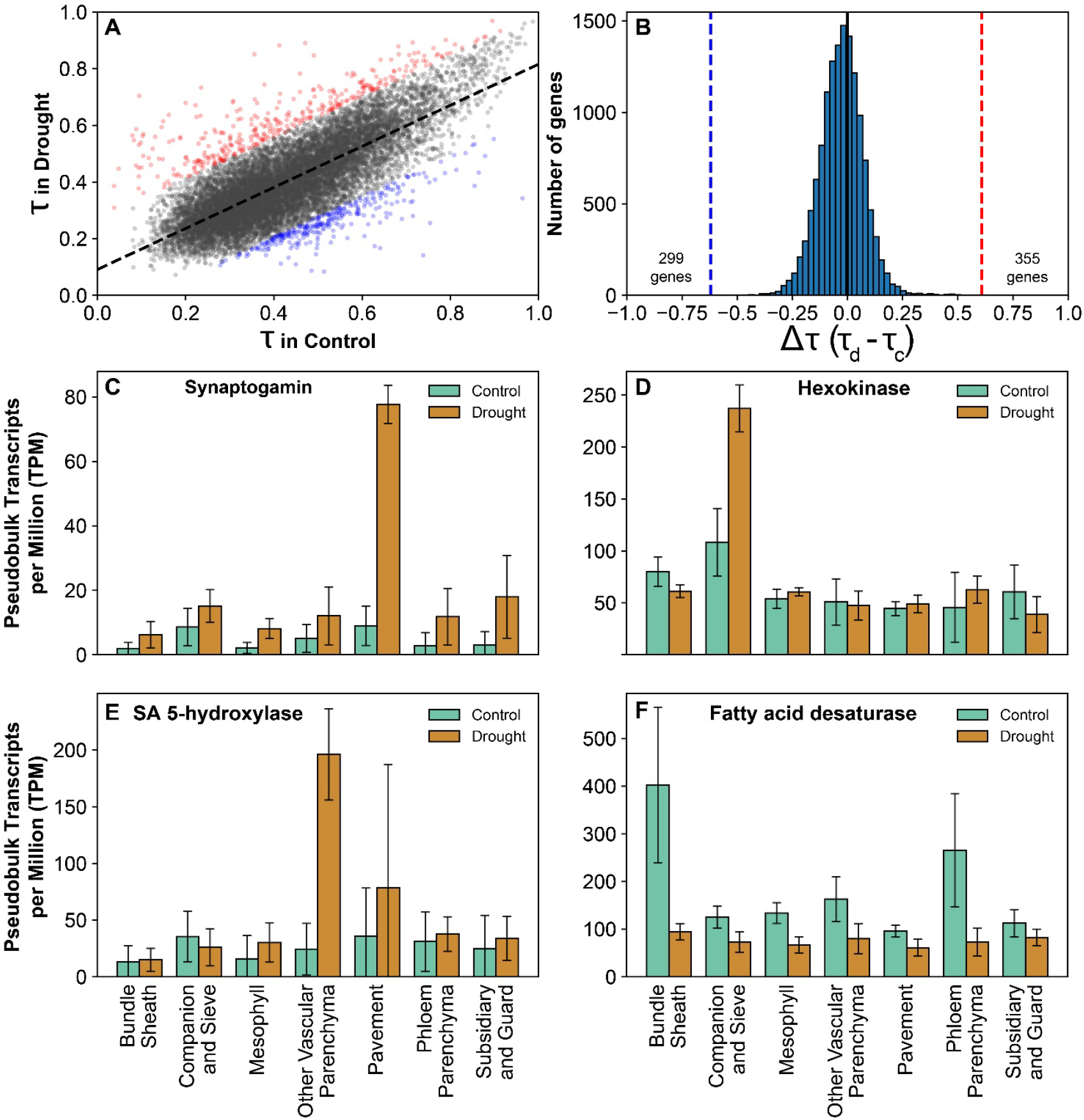
Cell-type-specific responses to drought. A, a scatter plot of tau values in control versus drought. The dashed black line shows the best-fit linear regression, and genes outside of a 95% prediction interval are colored red if they are more specific under drought conditions and blue if they are more specific under control conditions. B, a histogram of Δ tau values with the high and low extremes from (A) denoted with vertical blue and red dashed lines. C-F, gene expression across cell types under control and drought conditions for C, synaptotagmin (Sobic.010G236100), D, hexokinase (Sobic.003G291800), E, salicylic acid 5-hydroxylase (Sobic.001G314300), and F, fatty acid desaturase (Sobic.004G260800). Error bars show standard deviations based with n = 4, based on two day 6 and two day 10 samples per condition, as cell types show high similarity within condition despite sampling day (**Fig. S6**).

### Transcription factors regulating drought response

Given the cell-type-dependent and independent expression patterns in response to drought, we next sought to identify master regulators of the sorghum drought transcriptional program. We first tested for enrichment of conserved transcription factor (TF) binding sites associated with genes whose expression was increased in response to drought in any cell type. To do so, we leveraged our recent work that assessed the binding of 74 *A. thaliana* TFs in 10 plant species, including a closely related *S. bicolor* strain RTx430 (Baumgart *et al.,* 2025). In this study, conserved binding sites (those identified in the promoters of orthologous target genes from multiple species) were more frequently located within accessible chromatin, harbored reduced intra-species nucleotide diversity, and marked gene sets associated with significantly more GO terms than species-specific binding sites - all hallmarks of functionally active binding sites (Baumgart *et al.,* 2025). Sorghum binding sites were filtered to retain only sites that were either conserved in both rice and sorghum, or - if absent in rice - conserved in at least four eudicot species. Over-representation analysis identified 10 TF families whose binding sites were significantly enriched in drought-induced DEGs (**Fig. 4A**). TFs from all 10 families have sorghum orthologs that are leaf-expressed, and four families (c-repeat binding factor, CBF; cooperatively regulated by ethylene and jasmonate, CEJ1; G-box binding factor, GBF2-3; and a family including several WRKY TFs) show strong drought induction (**Fig. 4B**). In addition, when we generated a network of binding site presence among sorghum orthologs of the drought-enriched TFs, we found putative co-regulation among three TFs: calmodulin-binding transcription activator (CAMTA), CBF, and CEJ (**Fig. 4A-B**). CAMTA orthologs are highly expressed in leaves but unresponsive to drought at the transcriptional level, whereas the highest-expressed orthologs of CBF (Sobic.010G016500) and CEJ1 (Sobic.010G052600) TFs are both strongly induced by drought (**Fig. 4B**). This observation supports the hypothesis that a constitutive CAMTA module may act as a first response, potentially via post-transcriptional regulation or activation, priming stress pathways that are further activated by other drought-induced TFs. GO enrichment analysis of target gene sets for these TFs highlight a role for trehalose metabolism and protein phosphorylation in drought response as these are enriched among the targets of multiple highly expressed or drought responsive TFs (**Fig. S7**). Gene expression data provide strong indication of which particular sorghum orthologs within each TF family are most important for regulating drought response in leaves, providing promising putative master regulators of drought response in sorghum that will be valuable for breeding and engineering efforts aiming to improve WUE in this important crop.

**Fig. 4.**
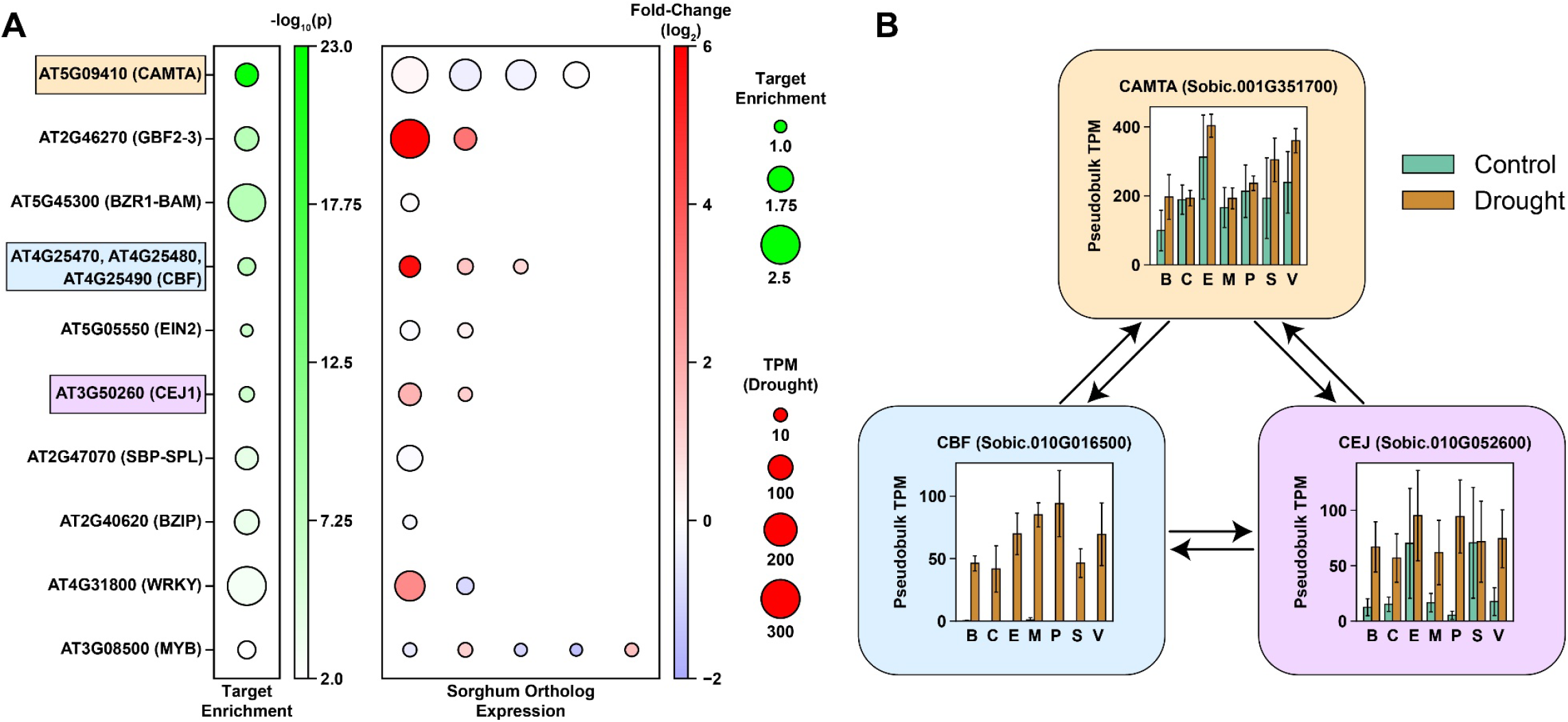
Key transcriptional regulators of drought response. A, over-representation analysis of binding sites for 76 conserved transcription factors (TFs) among drought-induced DEGs across cell types (first column) and expression of all sorghum orthologs over 10 TPM (all other columns). B, a co-regulatory module schematic with expression plots for the top sorghum orthologs of CAMTA, CBF, and CEJ TF families. Sorghum orthologs of all three TF families have binding sites for one another, implying a coordinated response to drought via these TFs. Cell type abbreviations are: B, bundle sheath cells; C, companion cells and sieve elements; E, epidermal pavement cells; M, mesophyll cells; P, phloem parenchyma cells; S, subsidiary and guard cells; V, other vascular parenchyma cells. Error bars show standard deviation with n = 4 samples, based on two day 6 and two day 10 samples per condition.

## Discussion

The mature leaf of plants is the source tissue providing energy to the rest of the plant for growth and development, as well as producing biomass and grains. Most single-cell transcriptomics studies to date focus on developing organs and single-cell transcriptome responses of the mature leaf to drought are not known. Our single-nucleus leaf transcriptomic analysis of *Sorghum bicolor* revealed that drought has a broadly-shared impact on gene expression across diverse cell types. Using Seurat-based clustering, we identified well-resolved leaf cell types under both well-watered and drought-stressed conditions. Most clusters mapped to a single cell type while two clusters - those corresponding to guard and subsidiary cells, and companion and sieve elements - could not be confidently distinguished based on known marker genes. Our final filtered dataset comprised over 55,000 high-confidence nuclei and enabled robust analysis of drought-induced transcriptional responses at the cell-type level.

Unexpectedly, we found that drought had a greater effect on transcriptomic variation than cell type identity. Across multiple analytic frameworks - including correlation matrices, principal component analysis (PCA), and gene set enrichment - cells grouped primarily by treatment rather than by identity. PC1 explained over a quarter of the variation and was strongly associated with drought response, capturing similar biological processes to those associated with drought-induced DEGs. These responses are consistent with known drought-adaptive mechanisms, including osmotic regulation, stress signaling, and suppression of photosynthesis. The tight clustering of cell types along the PC1 axis, and their largely parallel displacement under drought, points to a coordinated and conserved transcriptional response to water deficit across all mature leaf cell types.

This consistency in drought response across cell types is particularly striking given the cell-type specialization of C_4_ leaves, which results in significant biochemical, ultrastructural, and energetic differences between mesophyll and bundle sheath cells. Yet, our data show that drought-reduced genes contribute disproportionately to transcriptome-wide convergence, suggesting that repression of growth- and identity-associated programs under drought may act as a global cellular adaptation strategy. Indeed, PCA using only differentially expressed genes showed that drought-induced genes maintained or slightly enhanced cell-type distinctions, while drought-repressed genes drove convergence, underscoring their potential role in stress-induced reprogramming.

Although the global transcriptional response was largely consistent, we identified a subset of genes with strong cell-type-specific expression changes under drought. A synaptotagmin implicated in preserving plasma membrane integrity by removing diaglycerols produced by the phosphatidylinositol cycle under stress conditions (Ruiz-Lopez *et al.*, 2021b; Garcia-Hernandez *et al.*, 2025) was increased in transcript abundance specifically in pavement cells (**Fig. 3C**). This suggests either that epidermal cells are a key participant in the phosphatidylinositol cycle or that diaglycerols are particularly problematic for membranes in these cells. We also identified cell-specific responses in key metabolic and signaling components such as HXK1 in companion and sieve cells (**Fig. 3D**), which has been implicated in sugar sensing and tuning phloem loading and which is both drought-induced and has been implicated in drought responses in strawberry (Tong *et al.*, 2022; Wu *et al.*, 2023). Phloem-specific HXK1 induction under drought fits HXK1’s established role as a nuclear glucose sensor and signaling hub that couples carbon status to transcriptional programs rather than merely catalyzing glycolysis (Cho *et al.*, 2006; Sheen, 2014). HXK1-dependent sugar signaling intersects with abscisic acid (ABA) control of stomata: in *Arabidopsis*, guard-cell HXK1 overexpression drives sugar-induced, ABA-mediated stomatal closure and improves water-use efficiency and drought performance, and glucose-triggered closure relies on ABA signaling and HXK1 (Kelly *et al.*, 2013, 2019; Li *et al.*, 2018). Another example is a salicylic acid 5-hydroxlase, specifically induced in vascular parenchyma cells (**Fig. 3E**), which has mainly been examined for its role in biotic stress responses in other species (Zeilmaker *et al.*, 2015; Falcone Ferreyra *et al.*, 2015; Zhang *et al.*, 2017). Induction of SA hydroxylases in vascular parenchyma would locally reduce free SA and blunt SA-dependent signaling during drought, consistent with strong ABA-SA antagonism (Yasuda *et al.*, 2008; Ding *et al.*, 2016). Cell-type-specific downregulation of a fatty acid desaturase in bundle sheath cells (**Fig. 3F**) suggests that bundle sheath cells might remodel their membranes to reduce fluidity under drought since an *Arabidopsis* ortholog has been shown to provide polyunsaturated fatty acids to membranes (Launhardt *et al.*, 2024). Maintaining membrane integrity under water stress could be particularly important for bundle sheath cells given their high organellar content in C_4_ species. Such cell-type-restricted transcriptional shifts, while relatively few, point to distinct roles for different cell types in drought response and may be critical for modulating local stress responses or preserving essential functions under water limitation.

Leveraging global drought responses of the transcriptome across cell types and conservation of transcription factor (TF) binding sites across species (Baumgart *et al.*, 2025), we identified a set of TFs with conserved regulatory roles in drought response across angiosperms, whose binding sites were significantly enriched among drought-induced genes in sorghum. We identified the most critical sorghum orthologs within these TF families for drought response in sorghum leaves based on overall expression and drought induction. Four TF families (GBF2-3, CBF, CEJ1, and WRKY) have highly expressed, strongly drought-induced orthologs in sorghum. We also identified a co-regulated module among these, with sorghum orthologs of CAMTA, CBF, and CEJ1 all containing binding sites for one another. Within this module, CBF and CEJ1 orthologs were both highly expressed and strongly drought-induced in sorghum, while CAMTA orthologs were expressed but not drought responsive, suggesting differential regulatory deployment within the same transcriptional network. CAMTA TFs are known regulators of stress responses including drought and can act as activators or repressors of transcription, and have been shown to regulate other stress-responsive TFs including CBF TFs in Arabidopsis (Kim *et al.*, 2013; Pandey *et al.*, 2013). The *Arabidopsis* orthologs within the CBF and CEJ1 TF families are mainly implicated in response to cold stress in *Arabidopsis* (Tsutsui *et al.*, 2009; Chao *et al.*, 2022; Lee *et al.*, 2024; Gómez-Martínez *et al.*, 2024), indicating that the same TFs may regulate different stress responses in *Arabidopsis* and sorghum. Co-regulatory interactions and shared functional enrichment among the targets of these TFs highlight complex, possibly hierarchical control of drought-responsive gene modules, offering new avenues for mechanistic dissection and crop improvement.

Together, these results reveal that while *Sorghum bicolor* exhibits a coordinated, transcriptome-wide drought response that transcends cell identity, a minority of cell-specific responses may hold functional significance. The drought-responsive regulators and cell-enriched genes identified here provide promising targets for precision breeding and engineering aimed at improving water use efficiency and drought resilience in this important bioenergy and food crop.

## Methods Summary

Plants (*Sorghum bicolor* BTx623) were grown in a greenhouse under a 14 h photoperiod (30-32°C day/24-26 °C night) in Pro-Mix PGX with slow-release fertilizer. After two weeks of daily bottom watering to saturation, drought was imposed by adjusting soil water content to 40 % (measured gravimetrically) each morning for 10 days. On days 6 and 10, pooled samples (n = 3 plants per sample) were harvested for single-nucleus and bulk RNA-seq profiling. Nuclei were isolated via sorbitol-Triton density gradients, stained with SYBR Green, counted by flow cytometry, and ∼30,000 nuclei per sample were loaded on the 10X Genomics Chromium Next GEM Single Cell 3′ v3.1 (11 PCR cycles; 200–400 M reads/library on NovaSeq). snRNA-seq reads were filtered for rRNA (BBTools), aligned with CellRanger v7.0.1 to a nuclear and organelle genome reference, and processed through Velocyto and Seurat v5 (SCTransform, integration, clustering at resolution 0.1). Cell types were defined independently in control and drought samples, cross-validated via reciprocal module scoring to remove ambiguous assignments and supported by marker gene sets curated from multiple published studies (Berrio *et al.,* 2025; Delannoy *et al.,* 2023; Procko *et al.,* 2022; Sun *et al.,* 2022). Pseudobulk counts were analyzed with DESeq2 (design ∼ condition), GO GSEA, PCA on log-TPM values, and hypergeometric TF-binding site enrichment analysis. Full protocols and parameter settings are provided in Supplementary Information.

## Supporting information

Supplementary methods, results, & figures

Reciprocal control-drought cluster marker mapping

Cell type marker gene mapping, control

Cell type marker gene mapping, drought

## Acknowledgements

We thank members of the Rhee lab and the DOE Harnessing water to Optimize Crops (H2O-C) consortium for their helpful discussions. This work was funded in part by U.S. National Science Foundation grants (IOS-2312181, IOS-2406533, IOS-1546838, MCB-1916797, MCB-2052590, MCB-2420360, DBI-2213983, DBI-2419923, and OISE-2434687 to SYR) and U.S. Department of Energy, Office of Science, Office of Biological and Environmental Research, Genomic Science Program grants (DE-SC0018277, DE-SC0020366, DE-SC0023160, DE-SC0021286, and DE-SC0008769 to SYR). The work conducted by the US Department of Energy Joint Genome Institute (https://ror.org/04xm1d337), a DOE Office of Science User Facility, is supported by the Office of Science of the US Department of Energy operated under Contract No. DE-AC02-05CH11231. This work was done in part on the ancestral land of the Muwekma Ohlone Tribe which was and continues to be of great importance to the Ohlone people, and on the ancestral, traditional, and contemporary lands of the Anishinaabeg – Three Fires Confederacy of Ojibwe, Odawa, and Potawatomi peoples.

## Author contributions

Conceptualization: SYR, MS

Methodology: MS, SG, PK, RCO, YY, KK, MK, CZ

Investigation: MS, SG, RCO

Visualization: MS, SG Resources: SYR, RCO

Funding acquisition: SYR

Project administration: SYR

Supervision: SYR

Writing – original draft: MS, SYR

Writing – review & editing: MS, SYR, PK, SG

## Supplementary Information

- Methods

○ Detailed materials and methods.

- Results

○ Results of sequencing QC and cell filtering.

- Figures S1-S7

○ S1: The effect of experimental drought on biomass accumulation.
○ S2: Flow diagram of the cell type identification pipeline.
○ S3: Cell clusters under control and drought conditions.
○ S4: A UMAP of experimental conditions across merged cells.
○ S5: GO enrichment of drivers of PCA separation.
○ S6: A UMAP of all cell types by replicate.
○ S7: GO enrichment of transcription factor targets.

- Cell type identification appendix 1

○ Violin plots mapping marker genes from control cells to drought cells and vice versa to determine clusters representing the same cell types across conditions.

- Cell type identification appendix 2A and 2B

○ Violin plots mapping marker genes from the literature to control cell clusters (2A) and drought cell clusters (2B), used to determine cell type identities.

- Data sets

○ An Excel file containing cell type marker genes under both conditions, gene expression metrics (pseudobulk TPM values plus differential expression and extended tau statistics), and sorghum transcription factor target information. **Not included with preprint.**

