## Supplementary methods, results, & figures for "Single-cell-level response to drought in *Sorghum bicolor* reveals novel targets for improving water use efficiency"

### Supplementary Information

#### Materials and methods

##### Plant growth and experimental conditions

BTx623 sorghum plants were each grown in 125g of oven-dried PRO-MIX PGX soil (Premier Tech Horticulture, Quakertown, PA) supplemented with 3g each of Scotts slow-release Osmocote Pro 17-5-11 fertilizer (The Scotts Miracle-Gro Company, Marysville, OH). Prior to seeding, pots were saturated with water, allowed to sit until they ceased dripping, and weighed to obtain 100% soil water content (SWC) reference weight. Seeds were surface-sterilized with 90% ethanol for one minute then rinsed with DI water prior to planting. Three seeds per pot were started and thinned to one upon emergence. Plants were grown in a glass house on the Stanford campus in California with shade cloth blocking direct sunlight to reduce variability and supplemental LED lighting providing 800 µmol m⁻² s⁻¹ PPFD. The photoperiod was 14 hours with temperatures of 30-32°C during the photoperiod and 24-26°C at night. Plants were bottom-watered daily for two weeks, after which drought treatment began. Drought-treated plants were provided with limited water to bring the soil water content (SWC) back to 40% daily between 10am and 11am. On each of days 0, 2, 4, 6, 8, and 10 of the drought treatment, three drought treatment plants were sampled for above- and below-ground biomass measurements, and the average was used to correct for plant weight when calculating water to add to reach 40% SWC. The 40% SWC target for rewatering was based on the observation that SWC would decline as low as 10% prior to rewatering, resulting in a visible drought phenotype (wilted and paler green leaves), and that maintaining this reduction in water reduced growth by 10 days (**Fig. S1**). On days 6 and 10, samples for snRNA-seq and bulk RNA-seq were taken from the most recent fully emerged leaf with its ligule exposed. Two biological replicates for each time point and condition were used for this study. Each replicate comprised three individual plants in order to reduce variability. Samples were flash-frozen in liquid nitrogen and stored at -80°C prior to nuclei isolation for snRNA-seq or bulk RNA extraction.

##### Nuclei isolation, library preparation, and sequencing

Tissue samples were flash‐frozen and approximately 0.1 g of frozen sorghum leaf tissue was placed with one 2.4 mm and one 3.2 mm steel bead in a 2 mL screw‐cap tube. We added 1 mL of Gradient Solution A (0.275 M sorbitol, 10 mM MgCl₂) containing 0.15% Triton X-100 and 2 µL Antifoam B, then disrupted the tissue in a Bead Ruptor Elite at 4 m/s for 20 seconds (three cycles, 5 second rests). The homogenate was passed sequentially through 40 µm and 20 µm strainers into a 50 mL tube, layered over 4 mL of Gradient Solution B (0.4 M sorbitol, 10 mM MgCl₂, 0.1% Triton X-100), and centrifuged at 400×g, 4 °C for 5 min. This gradient spin process was repeated twice. The resulting pellet of nuclei was gently resuspended in Gradient Solution A, stained with SYBR Green, and counted on an Accuri flow cytometer. Approximately 30,000 nuclei were loaded onto the 10X Genomics Chromium platform (Next GEM Single Cell 3′ v3.1), with 11 cycles of cDNA PCR and a target of 200–400 million reads per library on the NovaSeq S4 or NovaSeq X platforms.

##### snRNA-seq quality control and data cleaning

Raw FASTQ files were first filtered with BBTools v38.96 (bbduk.sh) to remove read pairs whose second read contained 31-mers matching common ribosomal sequences. A nuclear reference for *Sorghum bicolor* BTx623 v5.1 was obtained from Phytozome and supplemented with the mitochondrial (NC_008360.1) and chloroplast (EF115542.1) genomes from NCBI. The rRNA‐filtered reads were processed with Cellranger count v7.0.1; resulting BAMs were then analyzed with Velocyto to generate spliced and unspliced count matrices. One library containing pooled sorghum and *Arabidopsis thaliana* nuclei was aligned to a combined reference, and only reads mapping to sorghum were retained. For each barcode, QC metrics - total UMIs, gene counts, proportion of organellar UMIs, spliced/unspliced ratio, and ambient RNA rate (via the R package diem) - were computed. Barcodes were retained if they had ≥ 400 nuclear UMIs, ≥ 250 nuclear genes, ≥ 90% nuclear UMIs, ≥ 15% unspliced UMIs, and a debris score ≤ 1. Filtered matrices (excluding organelle genes) were normalized per library using Seurat v5.0.0 SCTransform v2. Initial clustering (5,000 variable genes, 20 PCs, resolution 0.8) identified marker genes (logFC > 0.5, min.pct > 0.4, p_val_adj < 0.05); clusters lacking markers and with low UMI counts were removed as low quality. Finally, DoubletFinder was applied, and barcodes with a doublet score > 0.4 were excluded.

##### Cell clustering and cell type identification

SCT‐normalized objects were integrated by first selecting 3,000 shared features (SelectIntegrationFeatures), preparing them for integration (PrepSCTIntegration), identifying SCT‐based anchors across the first 30 dimensions (FindIntegrationAnchors, normalization.method="SCT", dims=1:30), and then merging all datasets with IntegrateData (normalization.method="SCT", dims=1:30). The pooled data were then separated into control and drought cell sets, and cell clustering was performed in each subset independently using the FindNeighbors and FindClusters functions, with a resolution parameter of 0.1. To identify clusters between conditions corresponding to the same cell types, marker genes were first identified for each cluster within the "Control" and "Drought" Seurat objects using FindAllMarkers, considering only positive markers (i.e., genes more highly expressed in that cluster relative to all others). A stringent filtering criterion was applied, retaining only markers with an adjusted p-value less than 0.001 and an average log_2_-fold change greater than 1.0. Subsequently, the filtered marker gene sets from each condition were used to calculate module scores in the other condition's clusters using AddModuleScore. These module scores were visualized across the clusters of the reciprocal condition, along with known marker genes for specific cell types, to identify analogous cell populations between conditions and assign cell type identities (Appendices S1 and S2).

##### Correlation analysis across cell types and conditions

A clustered heatmap was generated by computing Spearman rank correlations among paired-sample feature vectors (constructed from all genes with over 10 TPM expression in at least one cell type, and using both day 6 and day 10 expression data). The resulting correlation matrix was hierarchically clustered using Euclidean distance and average linkage. Pairwise Spearman rank correlations were grouped into three categories (same cell type across different conditions, different cell types within the same condition, and different cell types across different conditions). Distributions of correlations in each category were first tested for normality (Shapiro–Wilk) and homogeneity of variances (Levene’s test). Because all groups met assumptions of approximate normality and equal variances, we applied a one-way ANOVA to test for overall differences among categories, followed by Tukey’s HSD for pairwise post hoc comparisons.

##### Differential gene expression and GO enrichment

snRNA‐seq counts were aggregated to “pseudobulk” by summing gene‐level counts across all cells within each unique combination of sampling day, cell type, and condition using Seurat’s AggregateExpression function. The resulting pseudobulk count matrix and associated sample metadata were used to construct DESeq2 datasets via DESeqDataSetFromMatrix (design = ∼condition). For each day–cell type group we fit a negative‐binomial model in DESeq2 and recorded log₂ fold-changes, Wald statistics, and Benjamini–Hochberg–adjusted p-values for further analysis.

Gene set enrichment analysis (GSEA) was used to find gene ontology (GO) terms of the “biological process” category which were enriched among drought-induced DEGs (based on positive Wald statistics) and drought down-reduced DEGs (based on negative Wald statistics).

##### Principal component analysis

Pseudobulk TPM values across all cell types under control and drought were log-transformed and three separate PCAs were performed: one on all expressed genes (≥ 10 TPM), one on drought-reduced genes, and one on drought-induced genes. From the all-gene PCA, we generated two biplots (PC1 vs PC2 and PC3 vs PC4), and from each DEG PCA a single PC1 vs PC2 biplot was generated. In each biplot, pseudobulk samples are shown as points colored by condition, and gene loading vectors are shown as gray lines. Vector lengths are first set proportional to their raw PC loadings (i.e. to the 1st power) and then further amplified by raising those lengths to the 3rd power; this scaling makes genes with stronger contributions stand out visually. Each vector’s opacity is also tied to that same cubic scaling, so longer vectors appear darker.

##### Identification of cell-type-specific drought responses

Extended Tau values (τ_ext_) were calculated per gene for control and drought conditions on days 6 and 10. For each gene, the day 6 and day 10 values were averaged to obtain a single τ_ext_ estimate per condition. A Δτ_ext_ value was then defined as the difference between drought and control averages (τ_ext,drought_ − τ_ext,control_).

To identify genes with extreme differences in τ_ext_, we fit a linear ordinary least squares (OLS) regression of averaged τ_ext,drought_ against averaged τ_ext,control_ across all genes. A 95% prediction interval was calculated from the regression model, and genes with values outside the prediction interval were classified as outliers, and the corresponding Δτ_ext_ values were used to designate genes with extreme condition-specific differences in tissue specificity.

##### Transcription factor binding site enrichment analysis

Transcription factor (TF) binding site information based on DAP-seq results for conserved TFs (Baumgart *et al.*, 2025) was used to identify transcriptional regulators of drought response in sorghum. Over-representation of TF binding sites was carried out in Python using pandas and NumPy for data handling, SciPy’s hypergeom.sf for one-sided hypergeometric tests, and Statsmodels’ multipletests for Benjamini-Hochberg FDR correction. For each TF, the count of DAP-seq peaks overlapping promoters in the target gene set was compared to the genome-wide promoter background via a hypergeometric test. P-values were adjusted by the Benjamini–Hochberg procedure, and TFs with FDR < 0.1 and positive enrichment scores were retained for plotting and further analysis.

#### Results

##### Cell clustering, identification, and filtering

Cell type clustering using the FindClusters function in Seurat with a resolution parameter value of 0.1 identified 10 clusters in the pooled control cells and 9 clusters in the pooled drought cells. Reciprocal scoring of top marker genes for each cluster between conditions mostly indicated a one-to-one mapping of clusters between conditions, and mapping of marker genes for cell identities made most cell identities apparent, with a few exceptions: the same cluster in both conditions scored best for both guard cell and subsidiary cell marker genes; companion cells and sieve cells could not be distinguished from one another; and the smallest cluster in the control cells could not be assigned an identity based on any marker genes, with reciprocal cluster marker scoring indicated a partial match with both mesophyll and pavement cells in the drought data. Only cells which could be assigned identities and which had the same identity assigned directly and via label transfer between conditions were retained for further analysis; this resulted in removal of 8.2% of control cells and 12% of drought cells, leaving 25,794 and 29,234 cells with high confidence identity assignments per condition, respectively. Our final UMAP of filtered cell types identifies mesophyll cells, bundle sheath cells, pavement cells, phloem parenchyma cells, and other vascular parenchyma cells, along with two clusters comprised of multiple cell types, one including companion cells and sieve elements and the other subsidiary and guard cells. In the case of the former, there are two apparent sub-clusters but cell-type markers did not unambiguously indicate which was which; in the case of the latter, markers for both cell types strongly indicated the same cluster, and even rerunning the clustering process on just those cells in isolation with various resolution parameters was unable to further separate this group into sub-clusters (not shown).

#### Figures

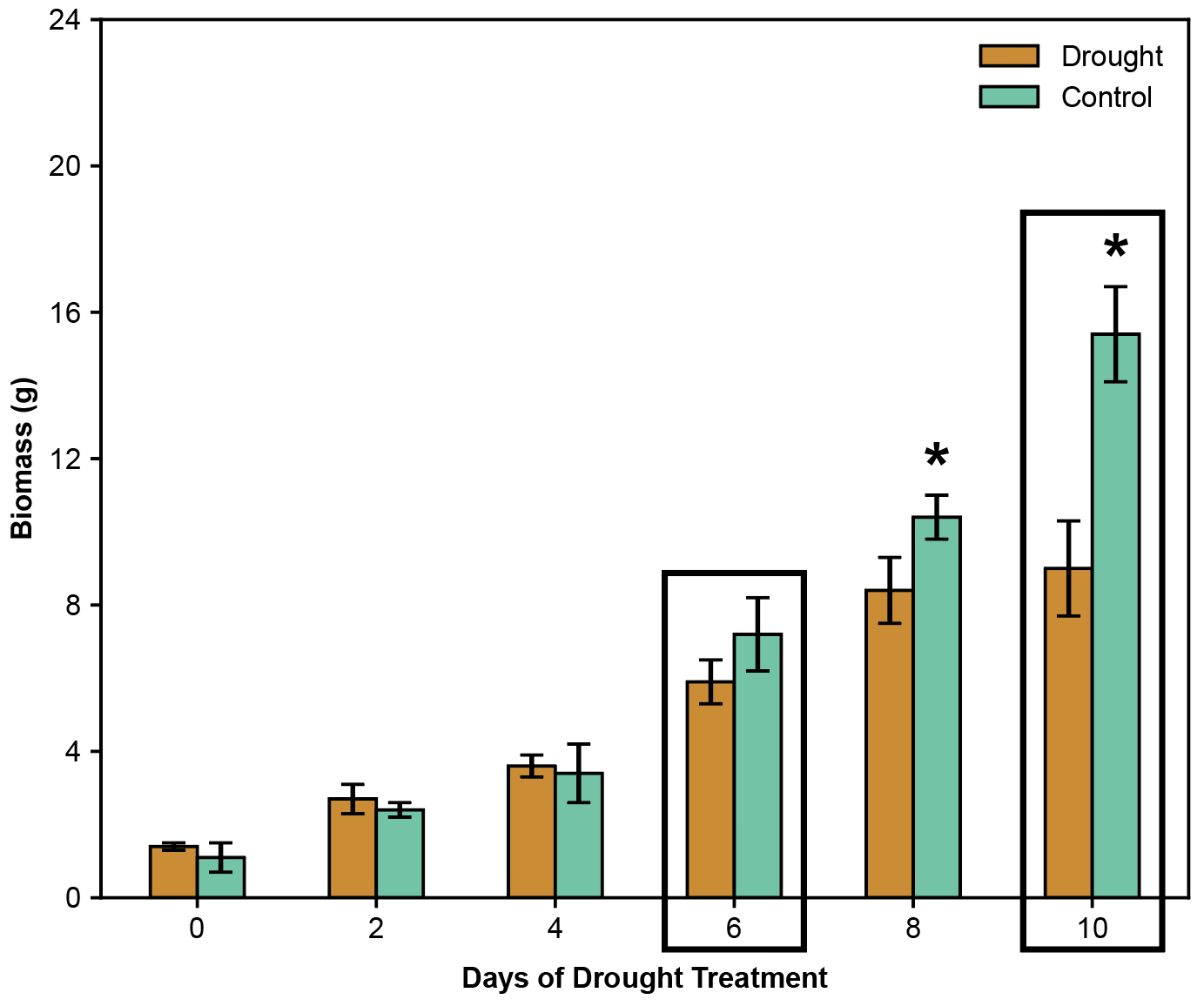

Supplemental Fig. S1: The effect of experimental drought on total (root and shoot) dry weight biomass accumulation, demonstrating that by 8 days into drought treatment there is a significant decline in growth. Black boxes denote sampling days for snRNA-seq. Asterisks denote significant difference (Welch’s two‐sample t-test with unequal variances, with raw p-values corrected for multiple comparisons using the Benjamini–Hochberg FDR procedure, α = 0.05). N = 6, three replicates from two repeated experiments.

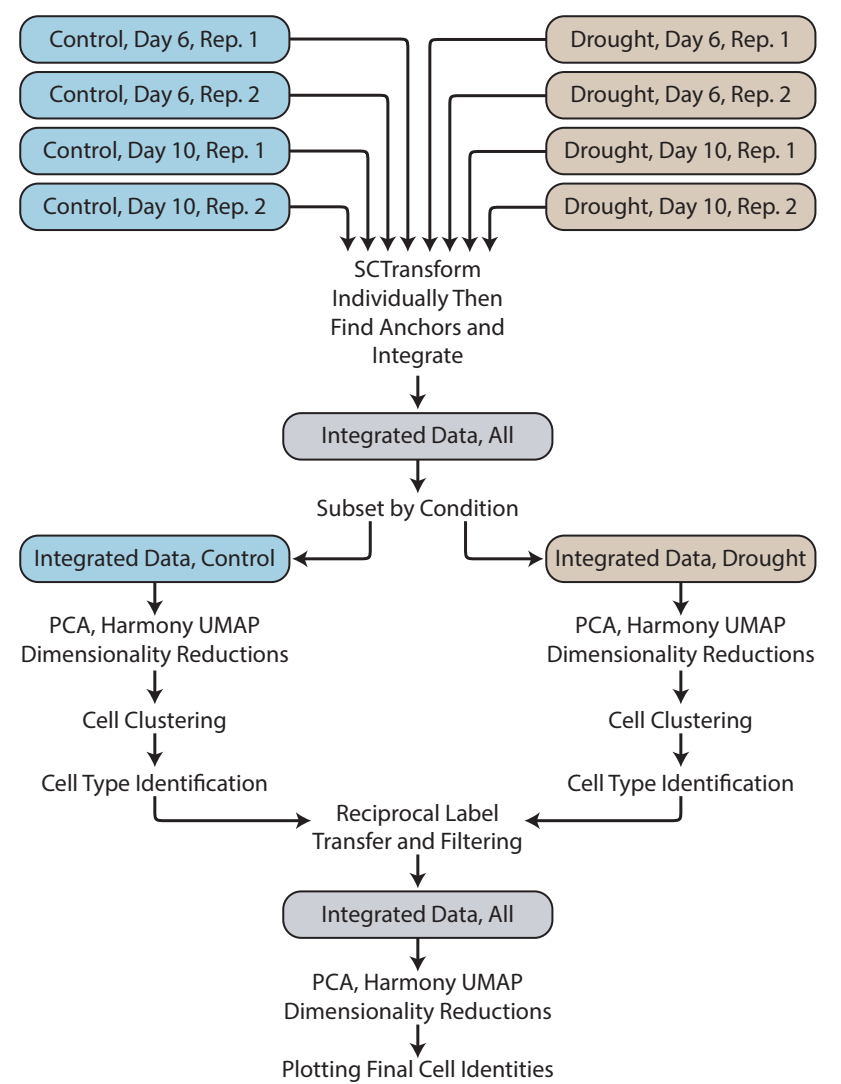

Supplemental Fig. S2: Flow diagram of the cell type identification pipeline. Individual libraries were normalized, integrated, then split by condition. For the control and drought data sets, cells were clustered, and cell types were identified based on identified based on four marker gene sets from the literature coupled with reciprocal marker gene mapping between control and drought cell clusters to identify equivalent cell types. We independently annotated each condition, performed reciprocal label transfer between conditions to map cells based on transcriptomic similarity and identify shared cell states, and excluded cells who’s direct and transferred labels conflicted - treating them as ambiguous - to obtain a high-confidence cell-type set.

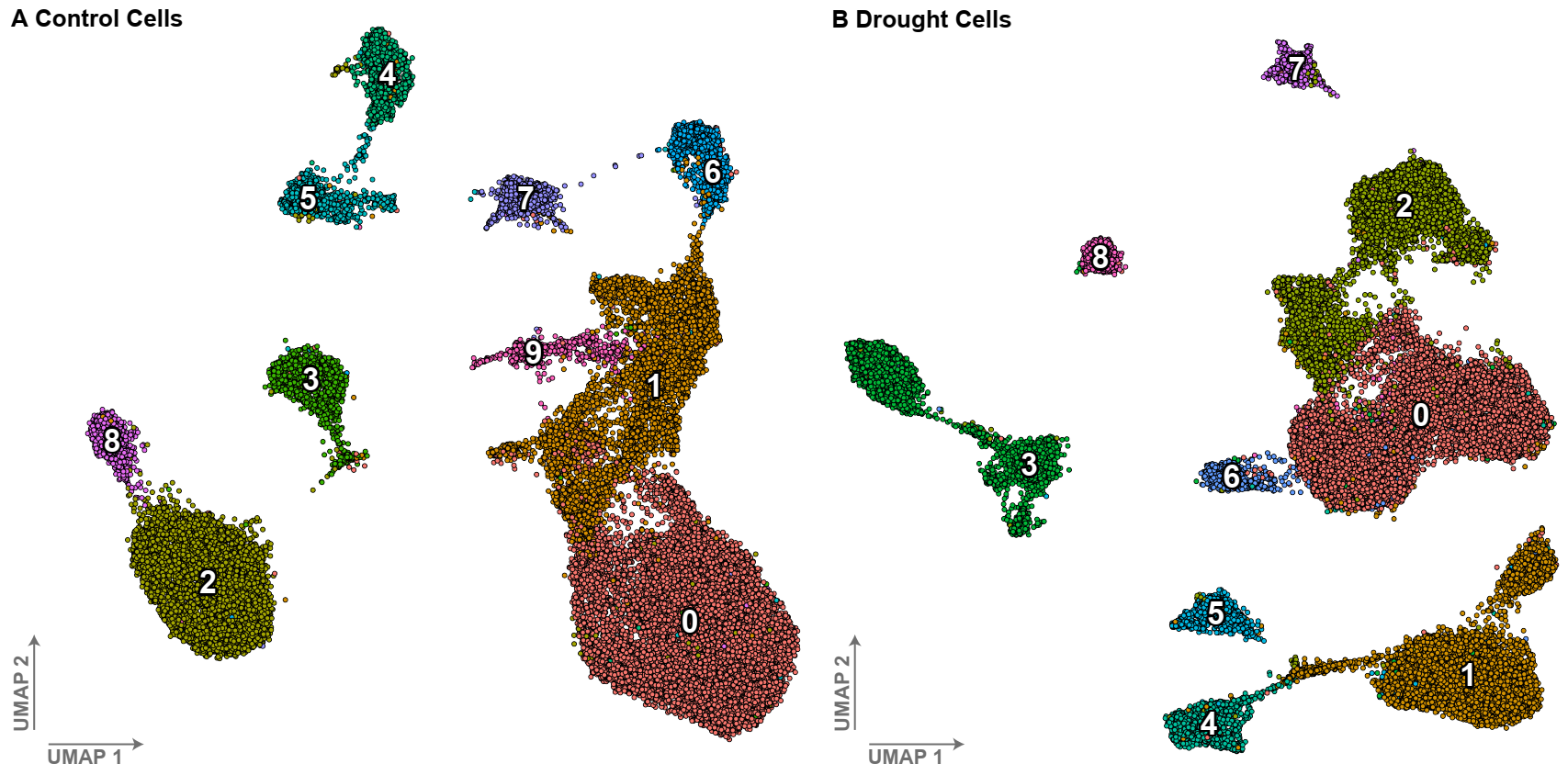

Supplemental Fig. S3: Cell clusters in control (A) and drought (B) single-nucleus datasets. Numbers and colors are assigned within each condition based on the number of cells in each cluster and do not necessarily represent the same cell types between conditions.

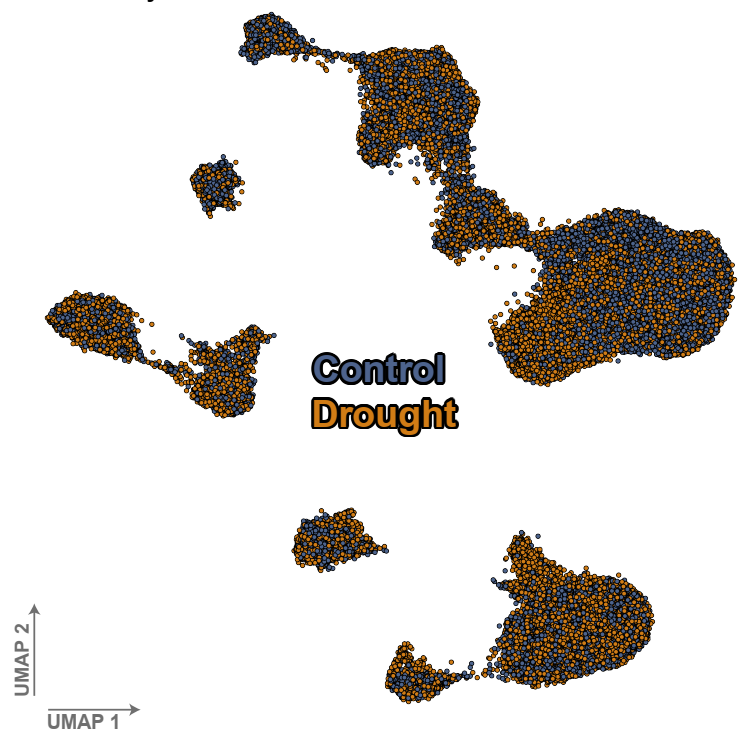

Supplemental Fig. S4: A UMAP of experimental conditions across merged cells. Strong overlap between conditions across all clusters indicates proper anchor-based integration of cell types between the two datasets.

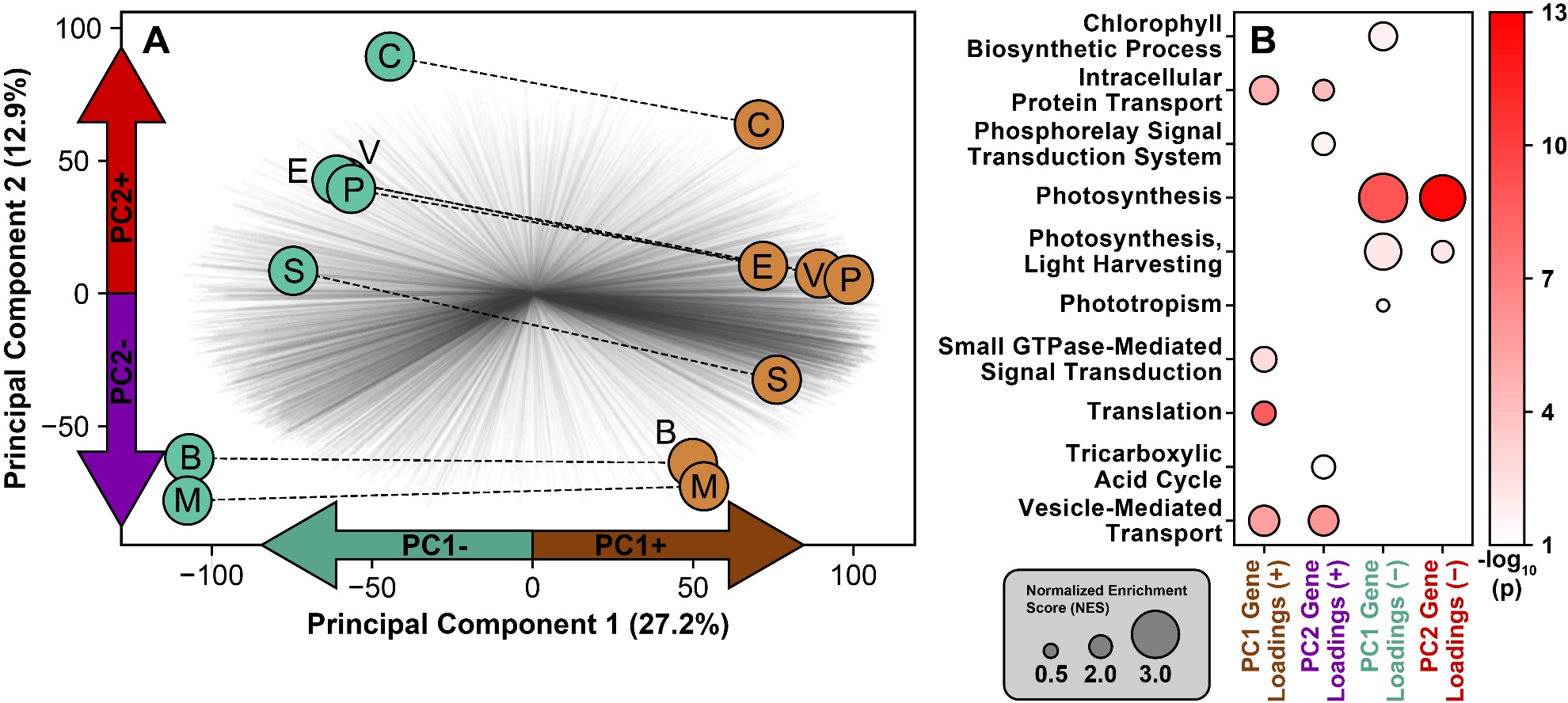

Supplemental Fig. S5: GO enrichment of drivers of PCA separation. A, the same principal component analysis shown in Fig. 2 based on all expressed genes, with gene loading vector directions indicated with colored arrows. B, a gene-set enrichment analysis (GSEA) bubble plot based on positive and negative drivers of PC1 and PC2 in (A). Bubble size and color denote normalized enrichment score (NES) and -log_10_(p) values, respectively. The Kolmogorov-Smirnov statistic and “elim” algorithm in TopGO were used with genes ranked by PC1 and PC2 loadings. Letters denote cell type: B, bundle sheath cells; C, companion cells and sieve elements; E, epidermal pavement cells; M, mesophyll cells; P, phloem parenchyma cells; S, subsidiary and guard cells.

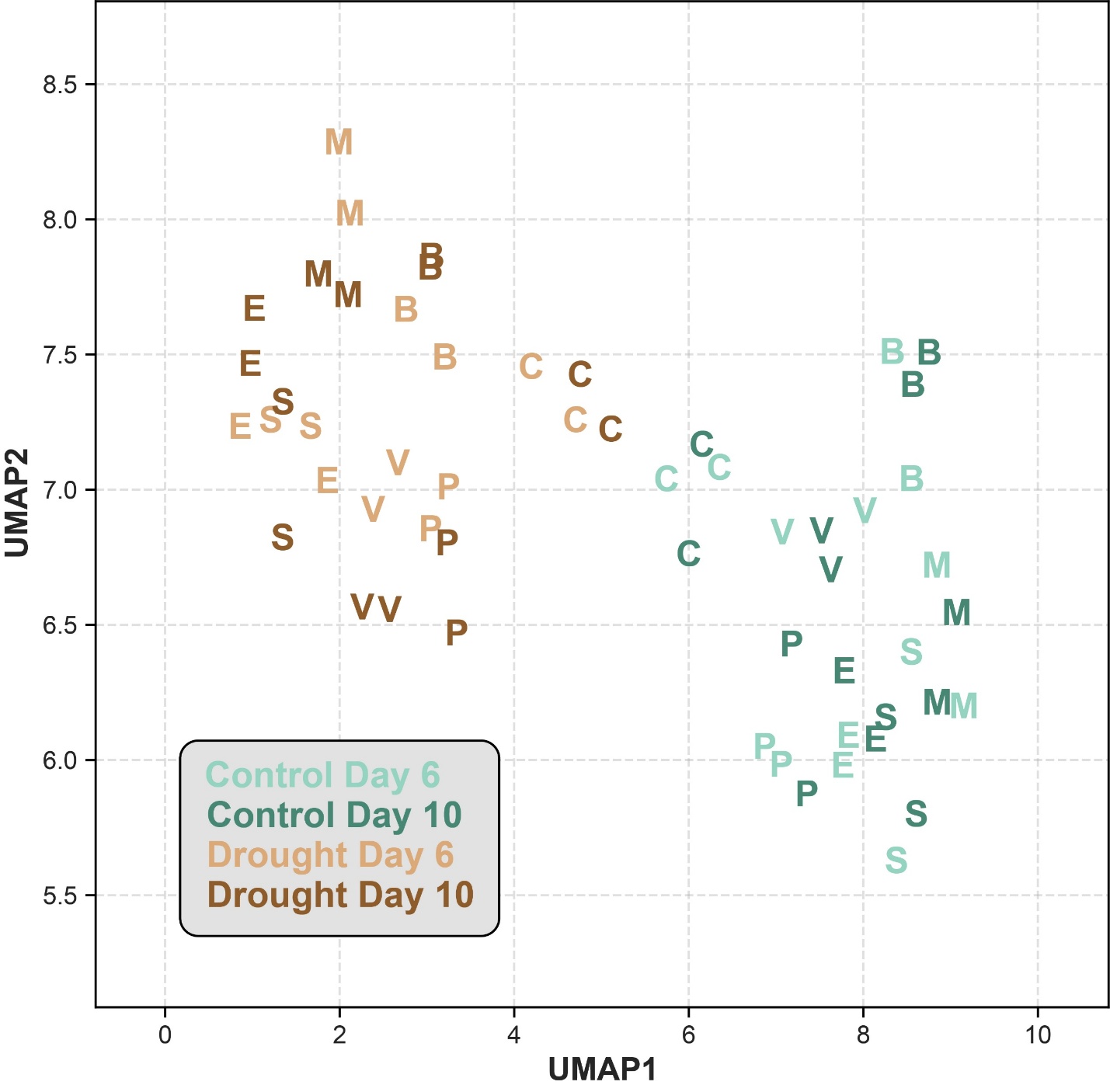

Supplemental Fig. S6: A UMAP of all cell types by replicate. Two replicates were sequenced for each combination of day and condition. Cell types cluster together within two larger clusters distinguished by condition, with little difference between the day 6 and day 10 samples within the same condition. Letters denote cell type: B, bundle sheath cells; C, companion cells and sieve elements; E, epidermal pavement cells; M, mesophyll cells; P, phloem parenchyma cells; S, subsidiary and guard cells; V, other vascular parenchyma cells.

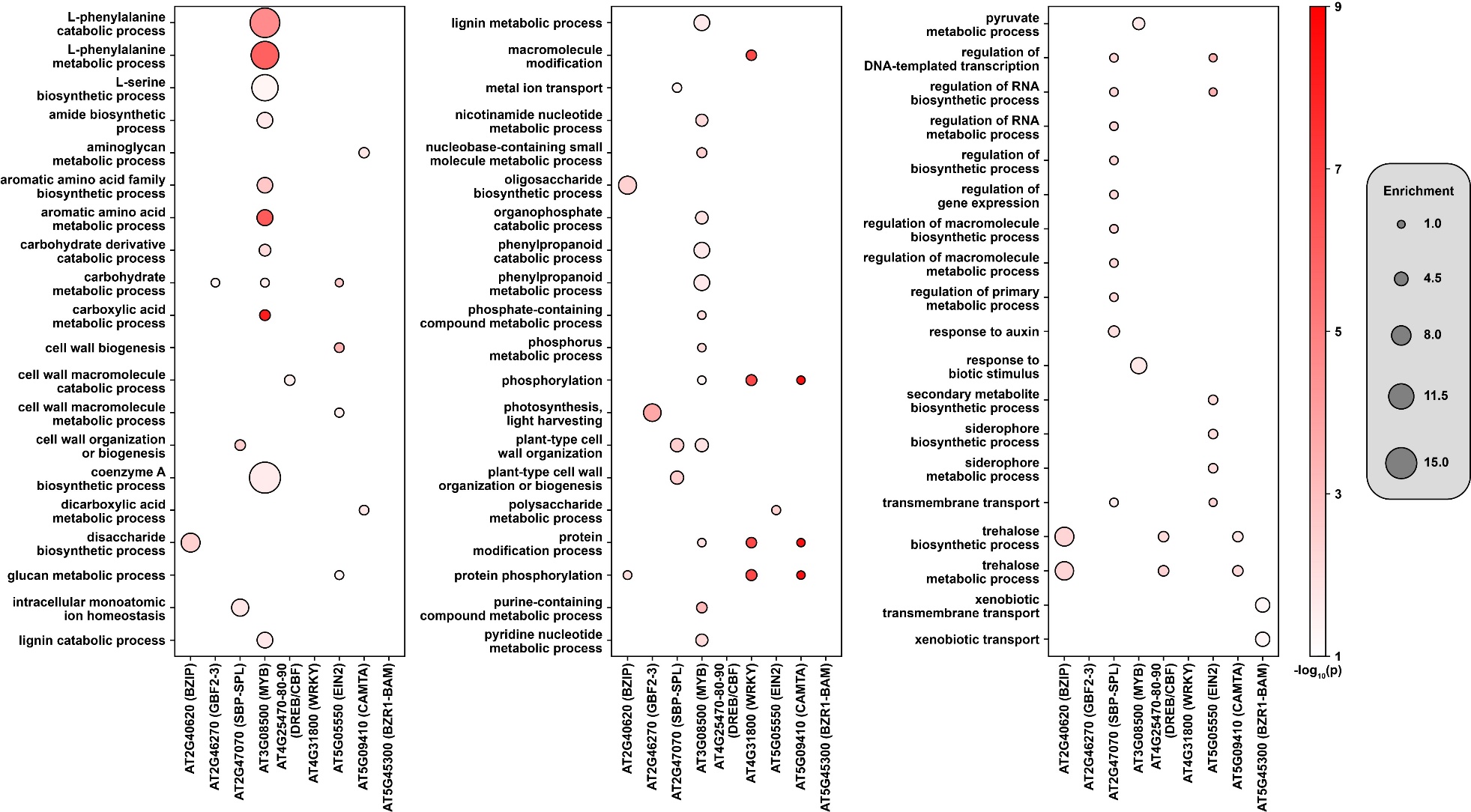

Supplemental Fig. S7: A bubble plot of enriched GO terms among the putative targets for the transcription factors in Fig. 4. Broader terms were pruned if they had a child term with a more significant adjusted p-value. No enriched GO terms were found among targets of CEJ (AT3G50260.
