## Supplementary material for "Single-cell-level response to drought in *Sorghum bicolor* reveals novel targets for improving water use efficiency": Reciprocal control-drought cluster marker mapping

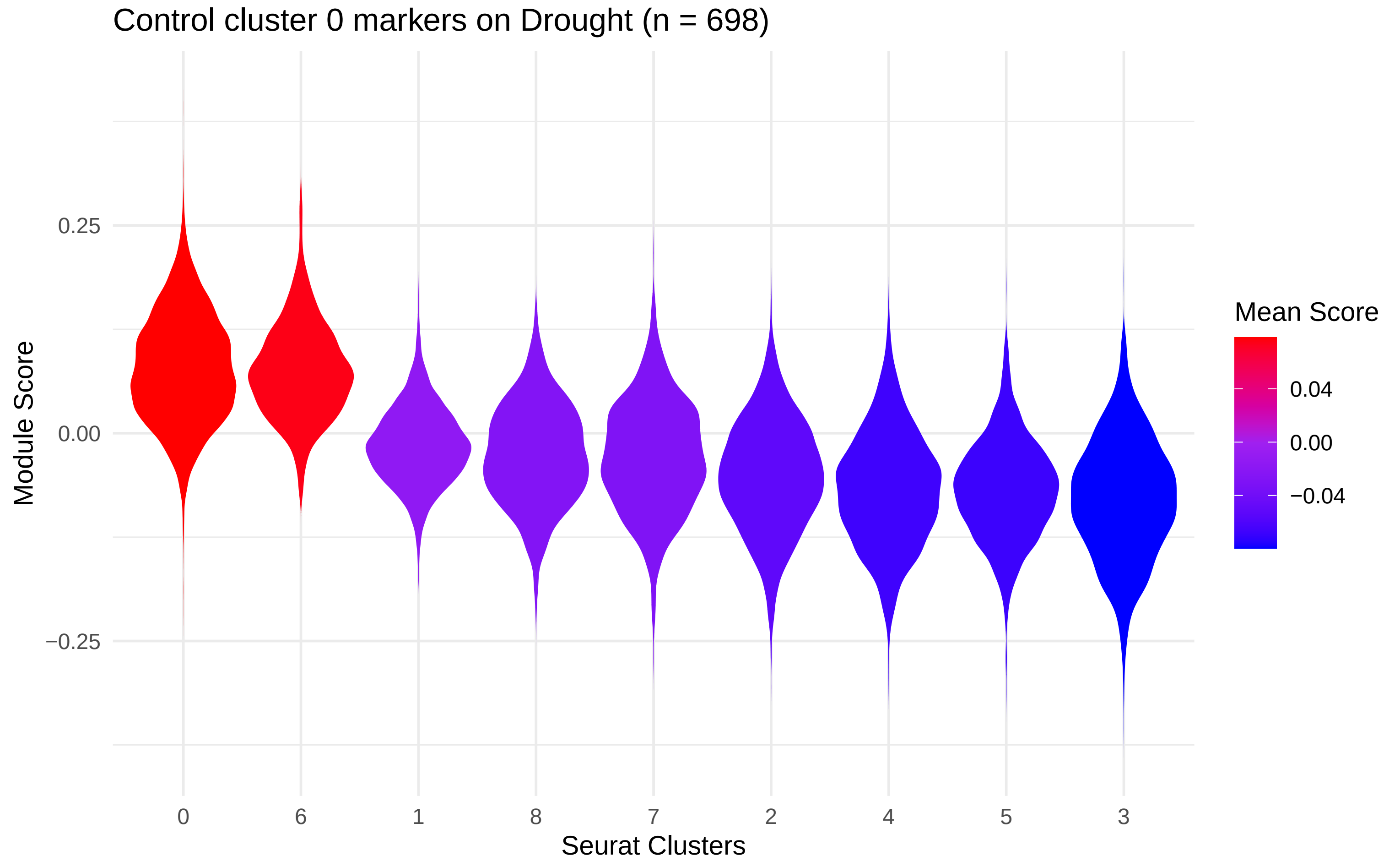

Control cluster 0 markers on Drought (n = 698)
Module Score
0.25
0.00
−0.25
0
6
1
8
7
2
4
5
3
Seurat Clusters
Mean Score
0.04
0.00
−0.04

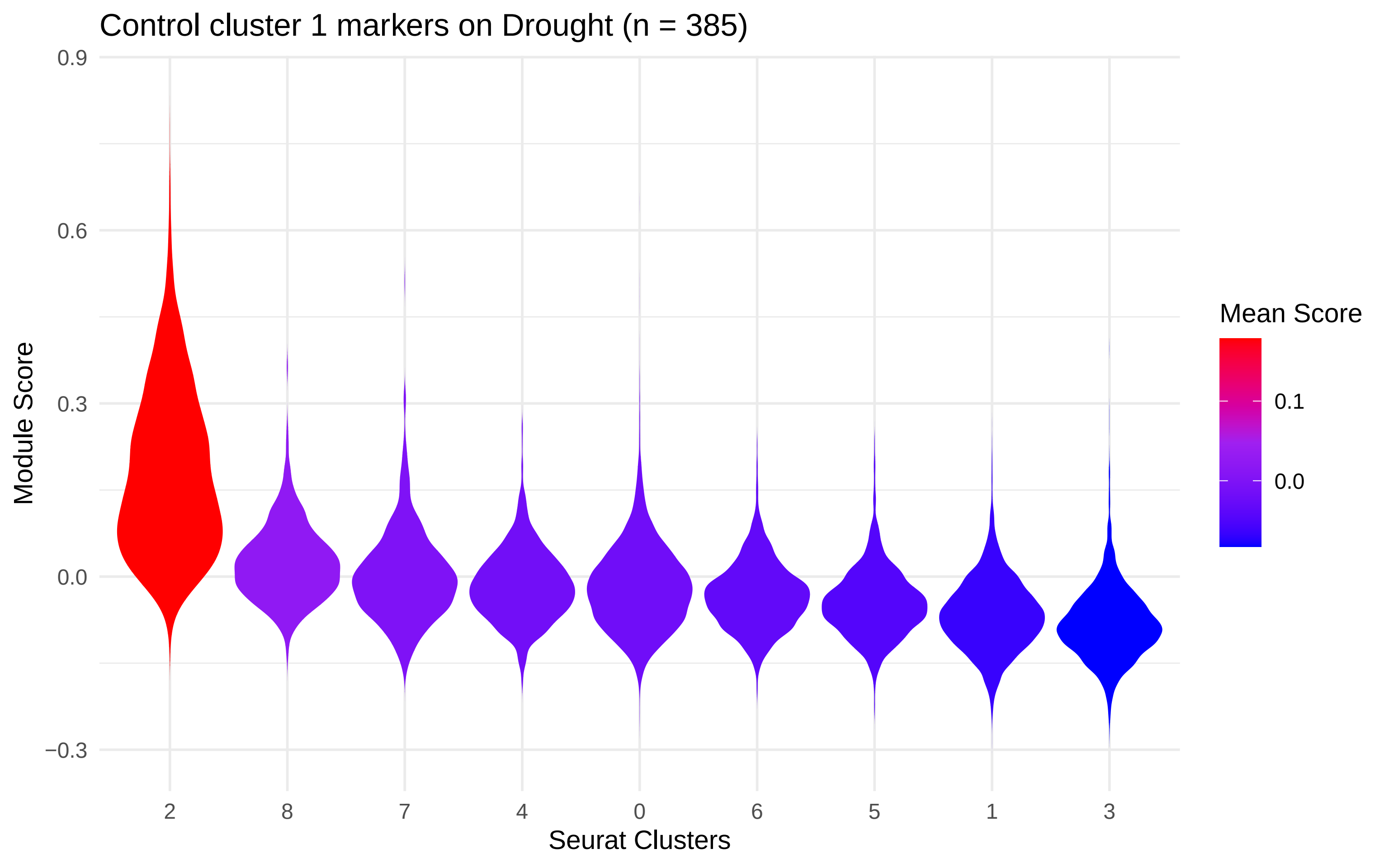

Control cluster 1 markers on Drought (n = 385)
0.9
0.6
0.3
0.0
−0.3
Module Score
2
8
7
4
0
6
5
1
3
Seurat Clusters
Mean Score
0.1
0.0

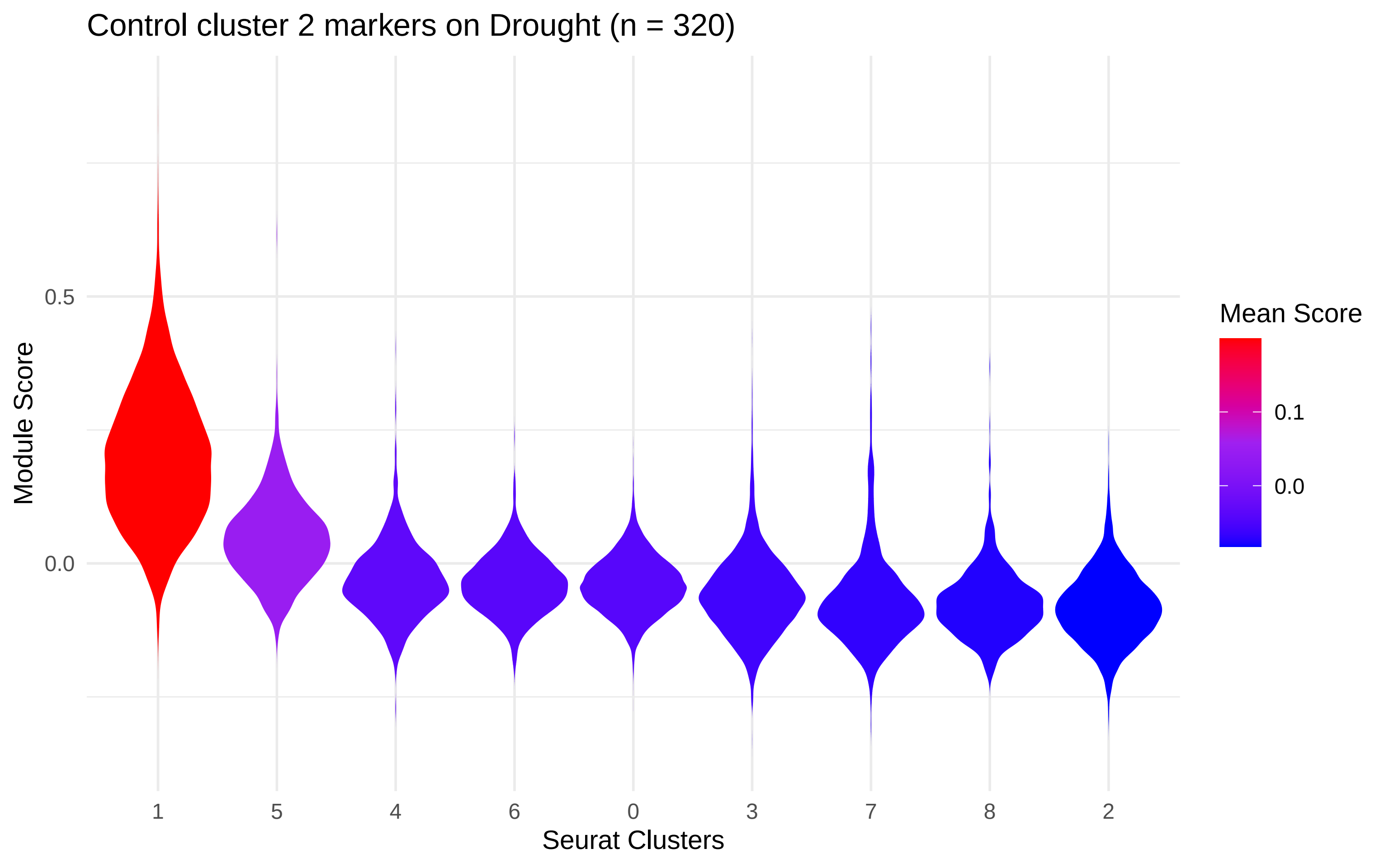
Control cluster 2 markers on Drought (n = 320)
Module Score
0.5
0.0
Seurat Clusters
1
5
4
6
0
3
7
8
2
Mean Score
0.1
0.0

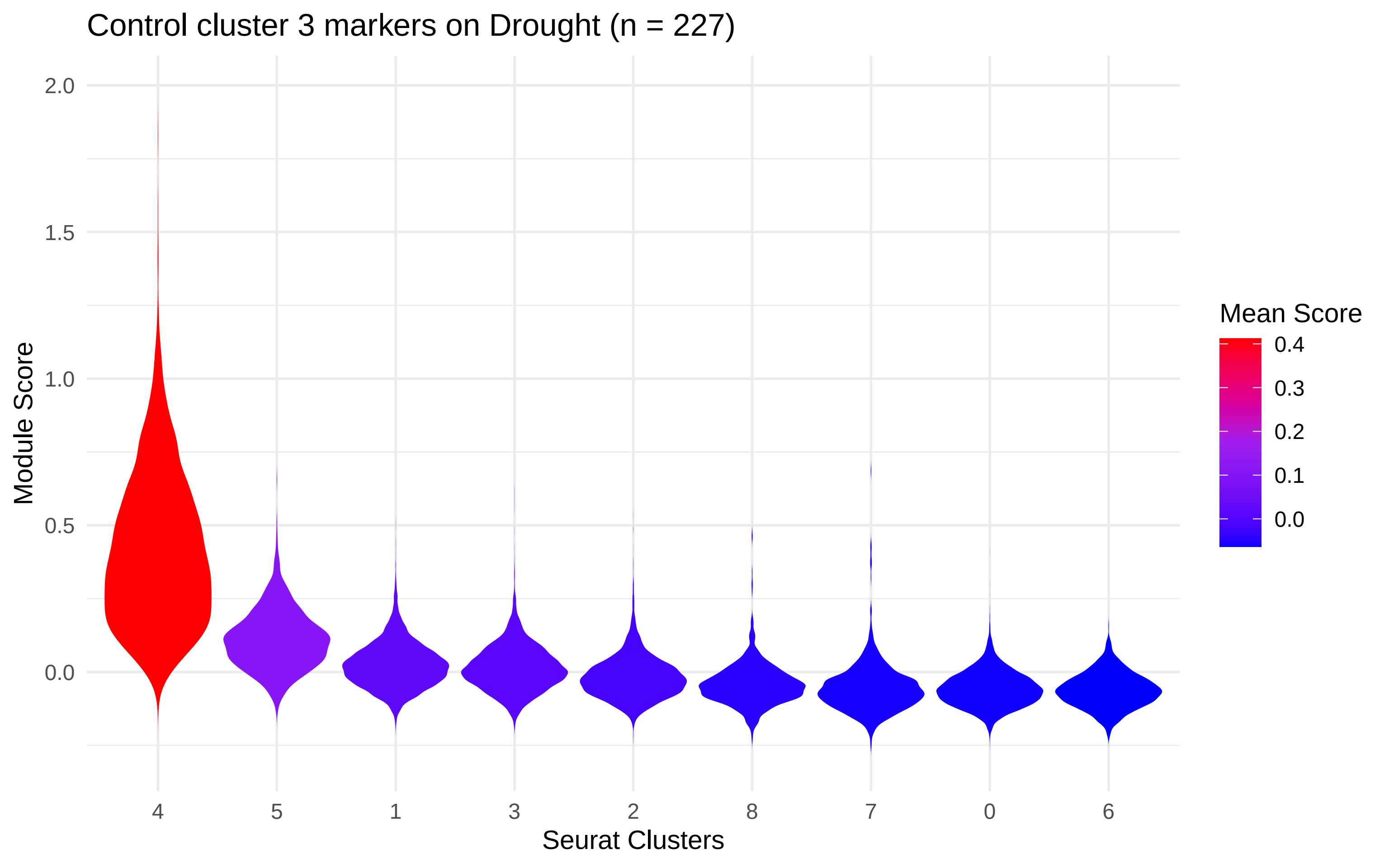

Control cluster 3 markers on Drought (n = 227)
Module Score
2.0
1.5
1.0
0.5
0.0
4
5
1
3
2
8
7
0
6
Seurat Clusters
Mean Score
0.4
0.3
0.2
0.1
0.0

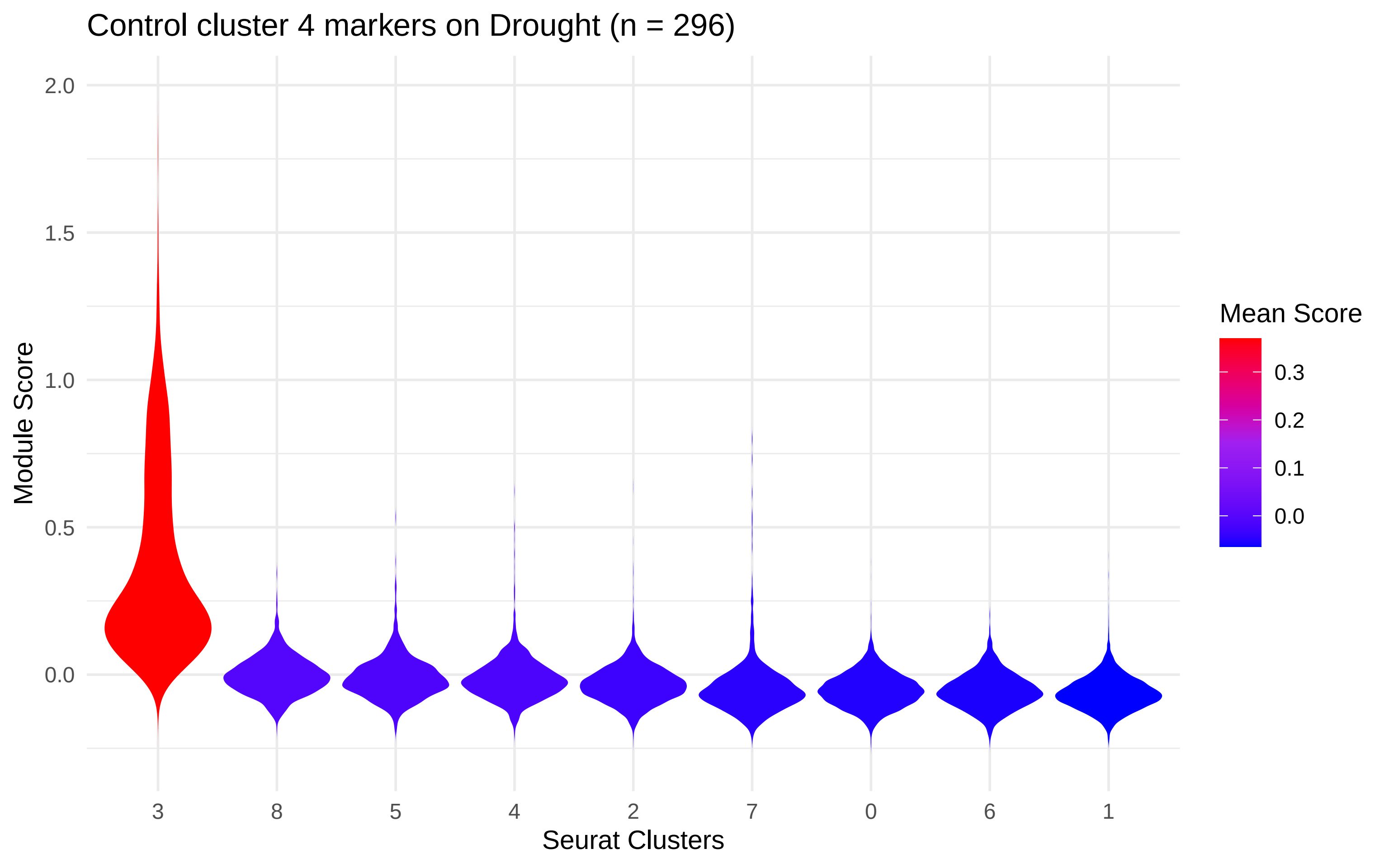

Control cluster 4 markers on Drought (n = 296)
Module Score
2.0
1.5
1.0
0.5
0.0
3
8
5
4
2
7
0
6
1
Seurat Clusters
Mean Score
0.3
0.2
0.1
0.0

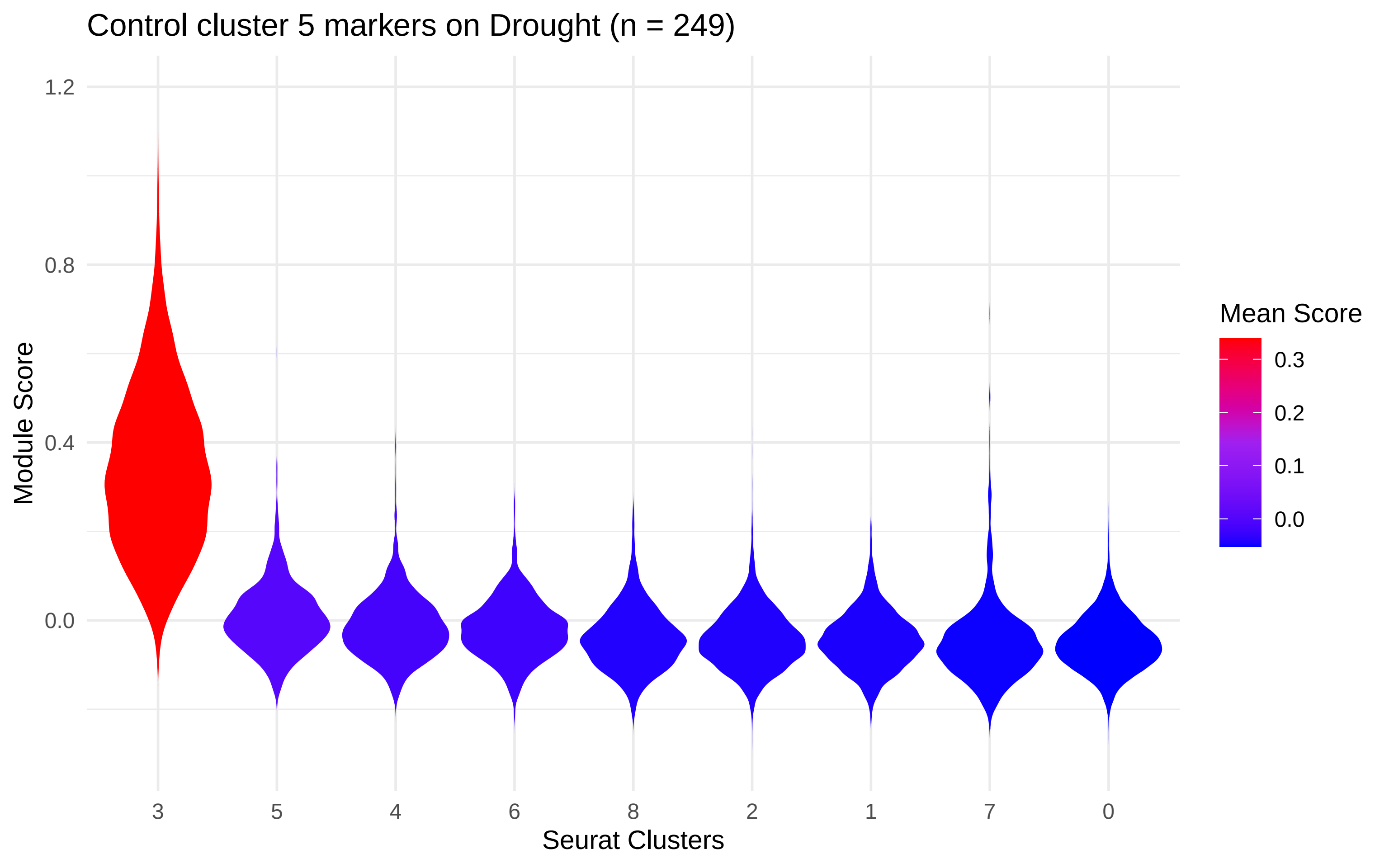

Control cluster 5 markers on Drought (n = 249)
Module Score
1.2
0.8
0.4
0.0
3
5
4
6
8
2
1
7
0
Seurat Clusters
Mean Score
0.3
0.2
0.1
0.0

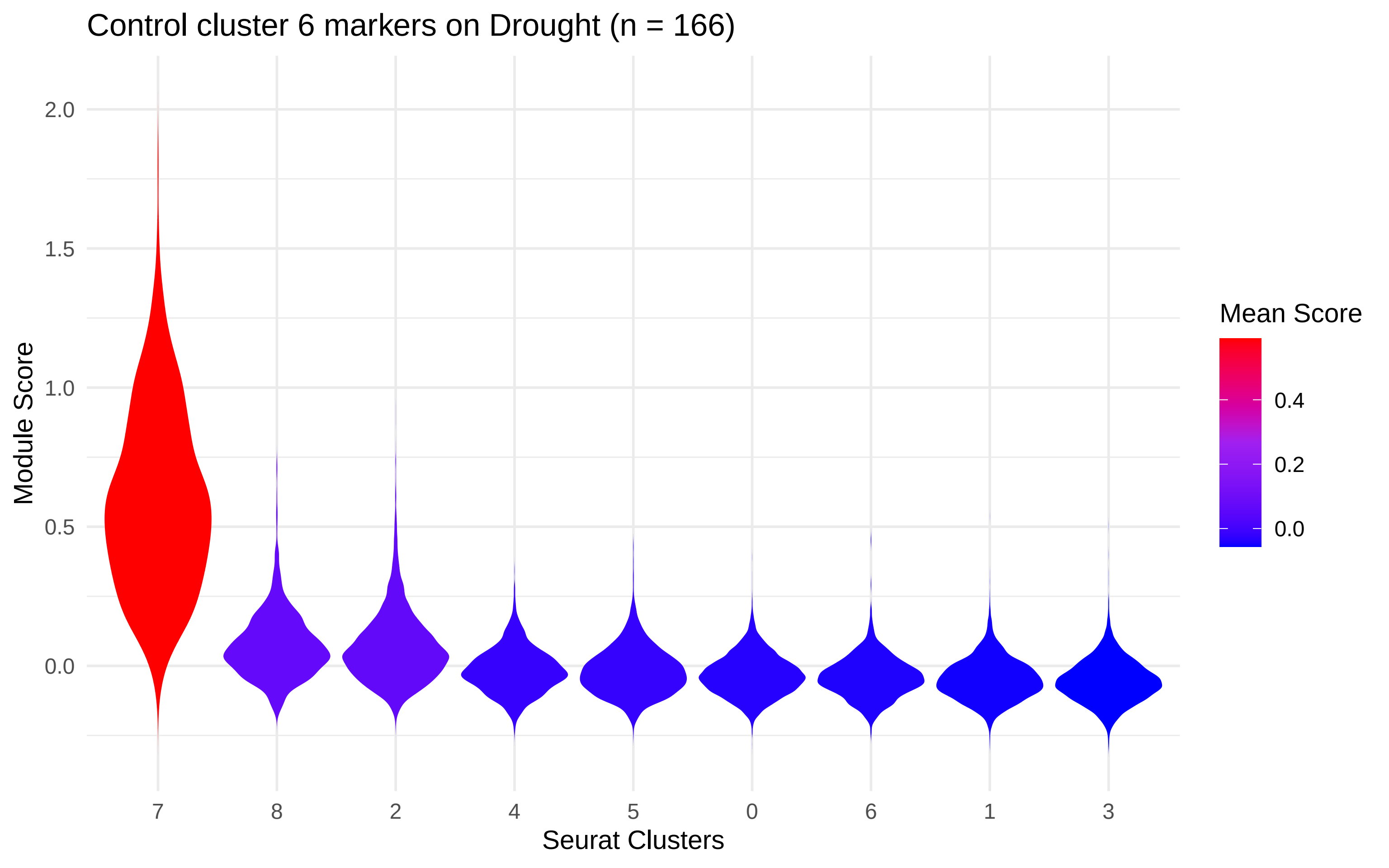
Control cluster 6 markers on Drought (n = 166)
Module Score
2.0
1.5
1.0
0.5
0.0
7
8
2
4
5
0
6
1
3
Seurat Clusters
Mean Score
0.4
0.2
0.0

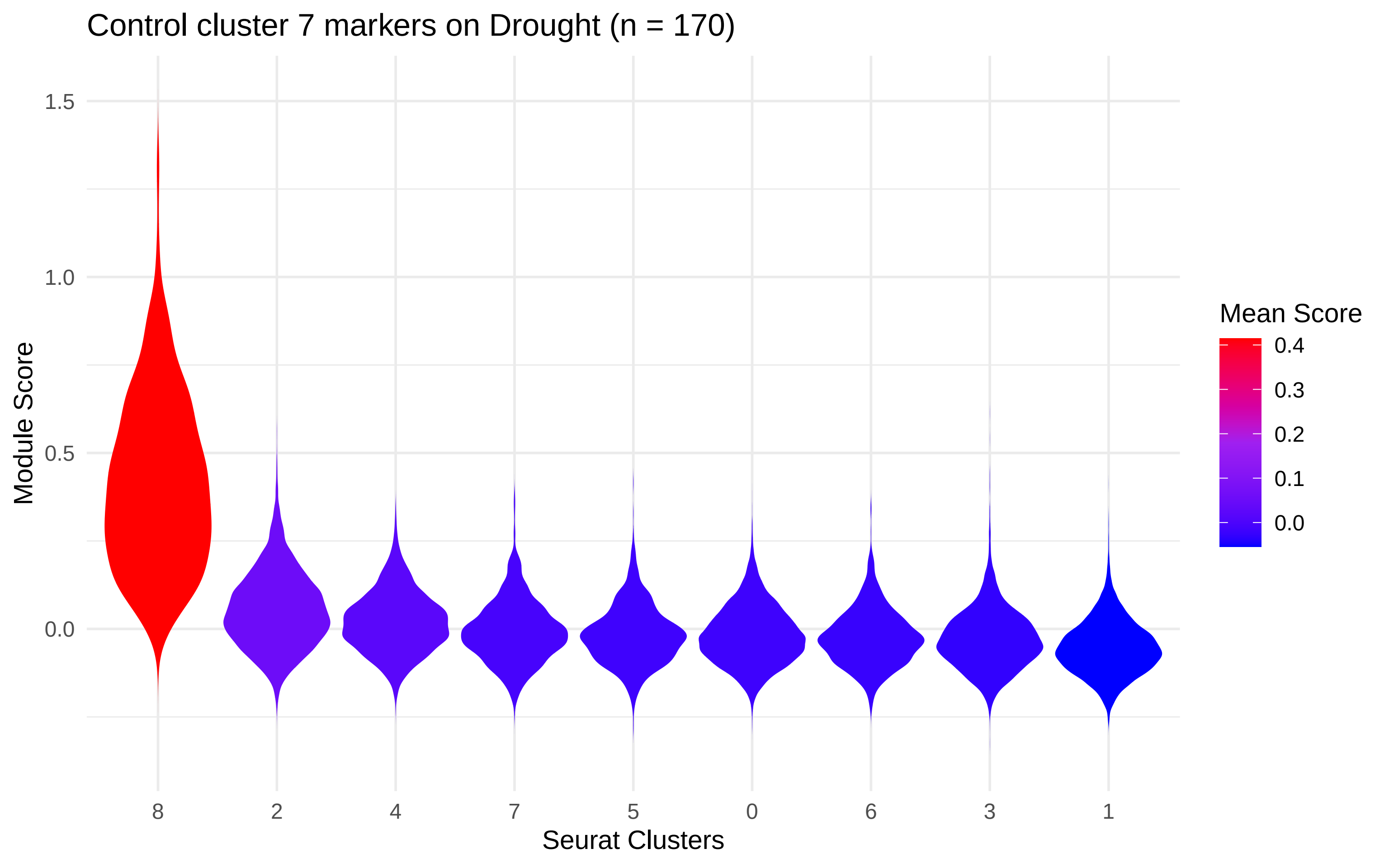

Control cluster 7 markers on Drought (n = 170)
Module Score
1.5
1.0
0.5
0.0
8
2
4
7
5
0
6
3
1
Seurat Clusters
Mean Score
0.4
0.3
0.2
0.1
0.0

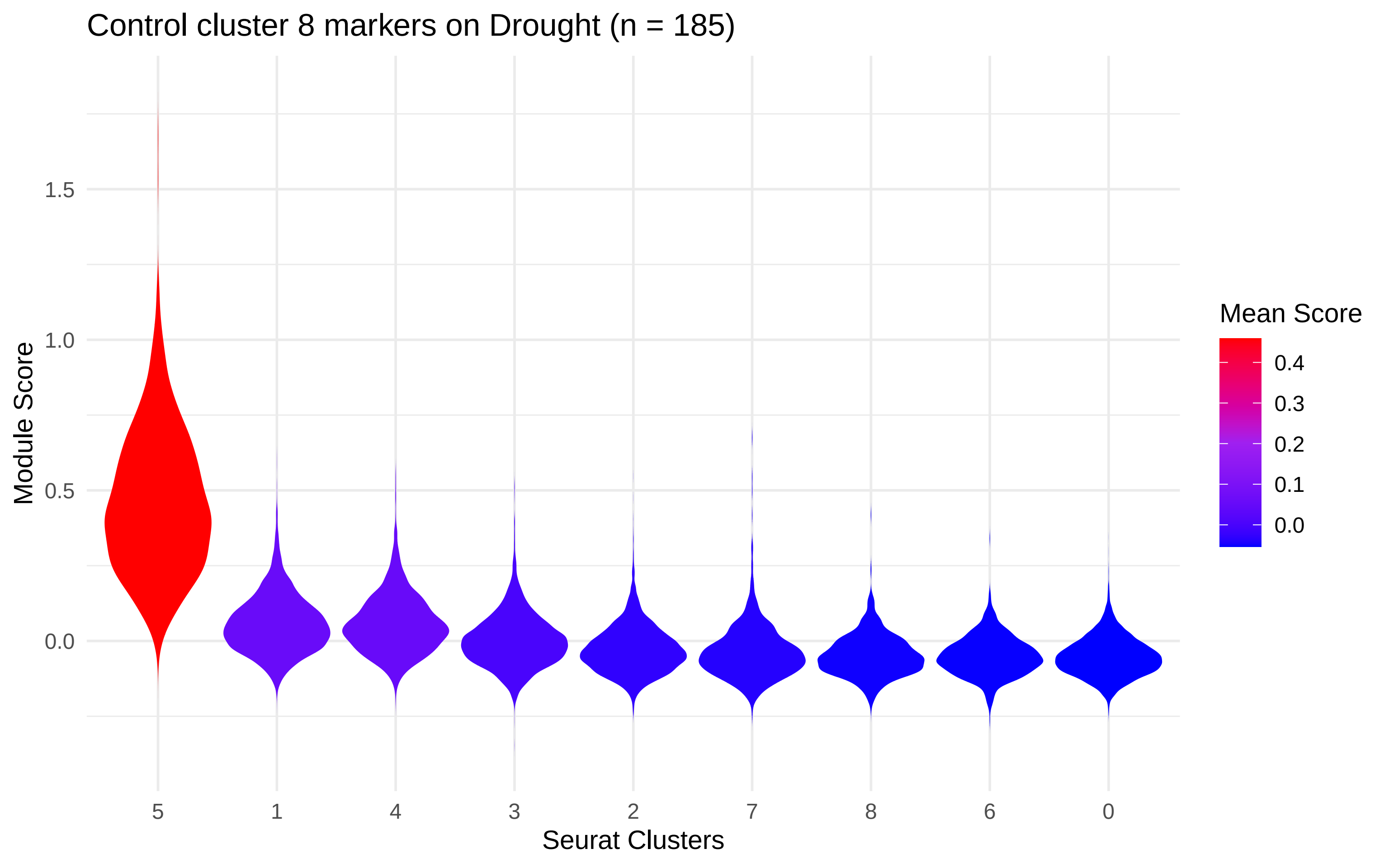
Control cluster 8 markers on Drought (n = 185)
Module Score
1.5
1.0
0.5
0.0
5
1
4
3
2
7
8
6
0
Seurat Clusters
Mean Score
0.4
0.3
0.2
0.1
0.0

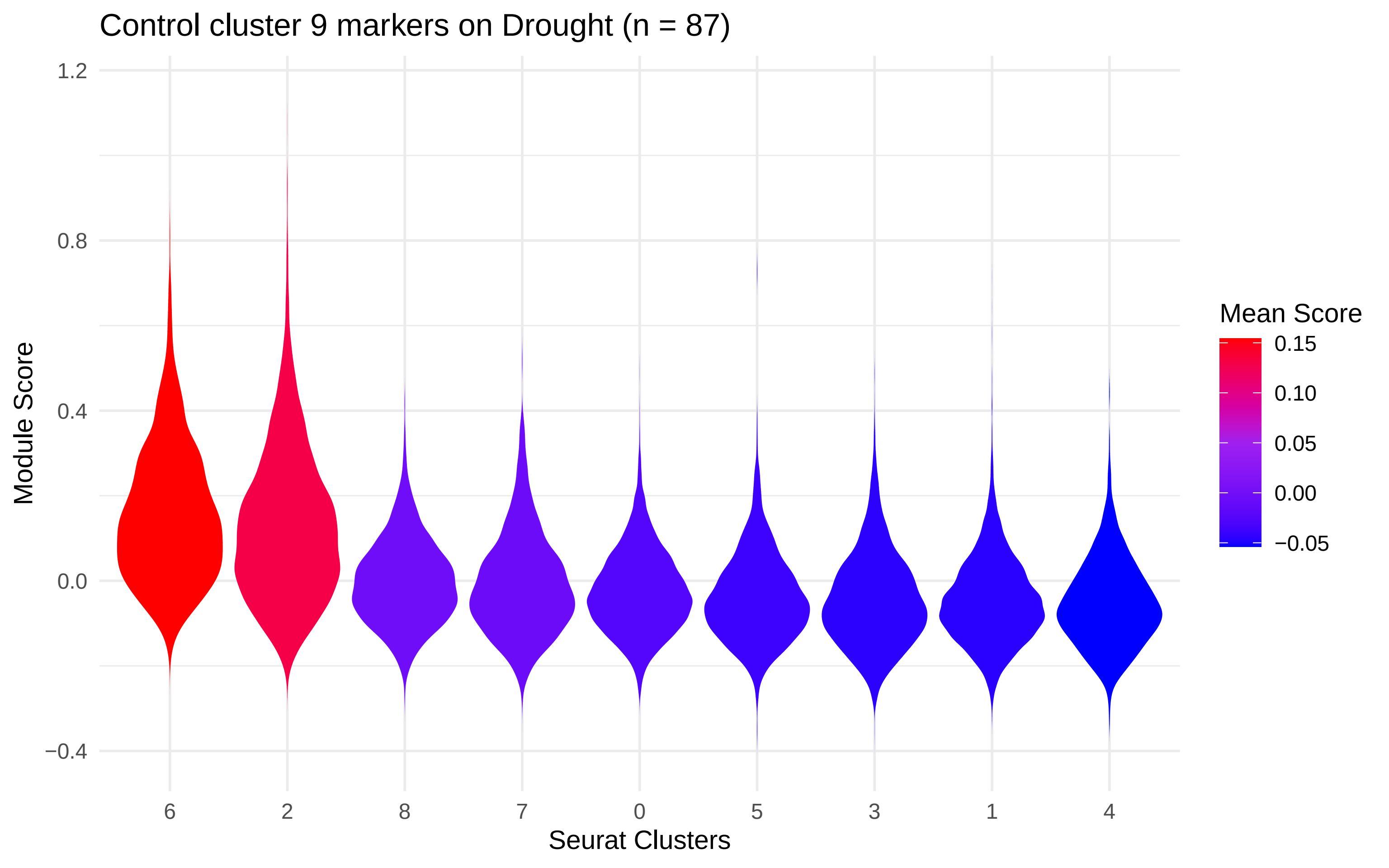
Control cluster 9 markers on Drought (n = 87)
Module Score
1.2
0.8
0.4
0.0
−0.4
6
2
8
7
0
5
3
1
4
Seurat Clusters
Mean Score
0.15
0.10
0.05
0.00
−0.05

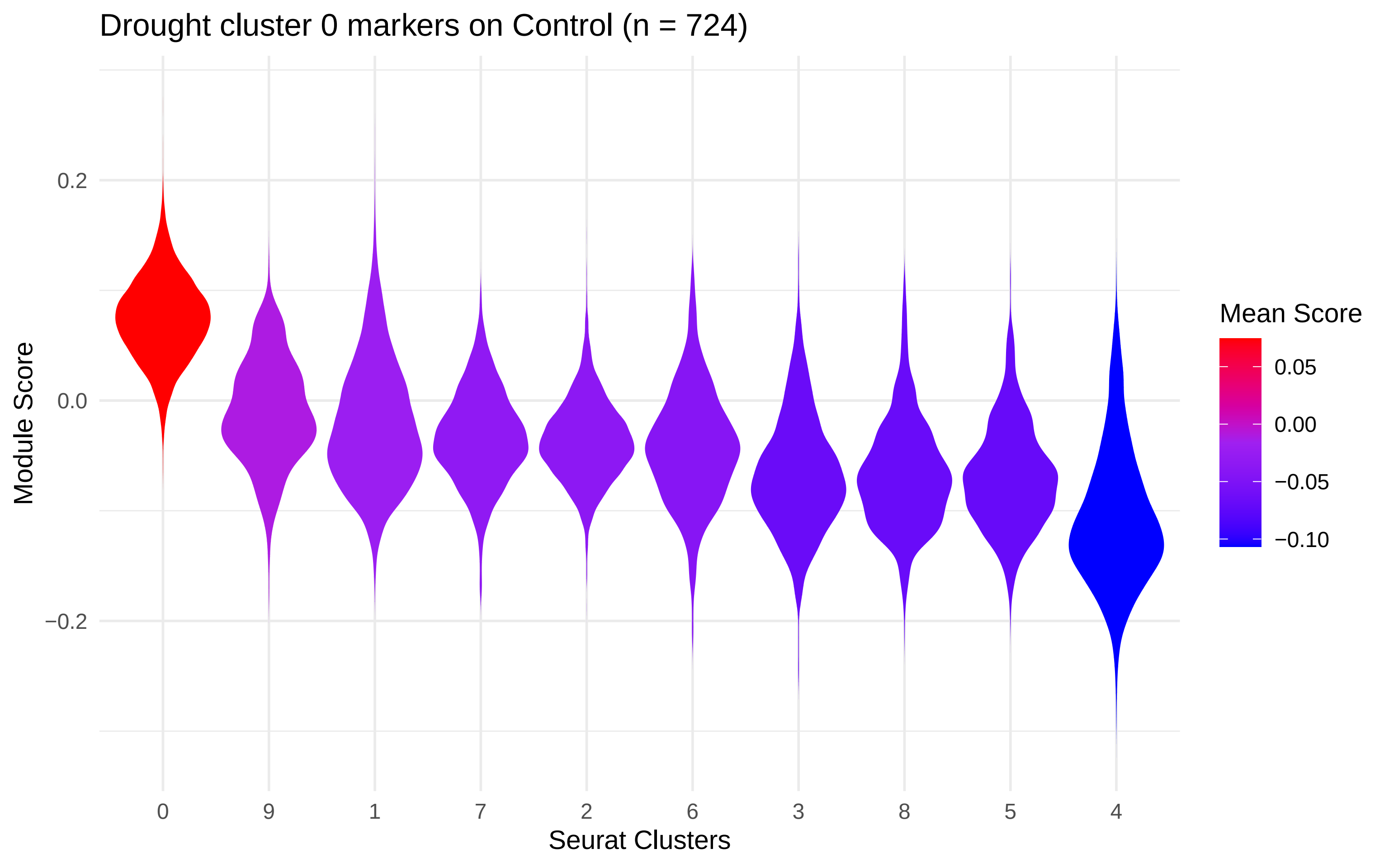

Drought cluster 0 markers on Control (n = 724)
Module Score
0.2
0.0
−0.2
0
9
1
7
2
6
3
8
5
4
Seurat Clusters
Mean Score
0.05
0.00
−0.05
−0.10

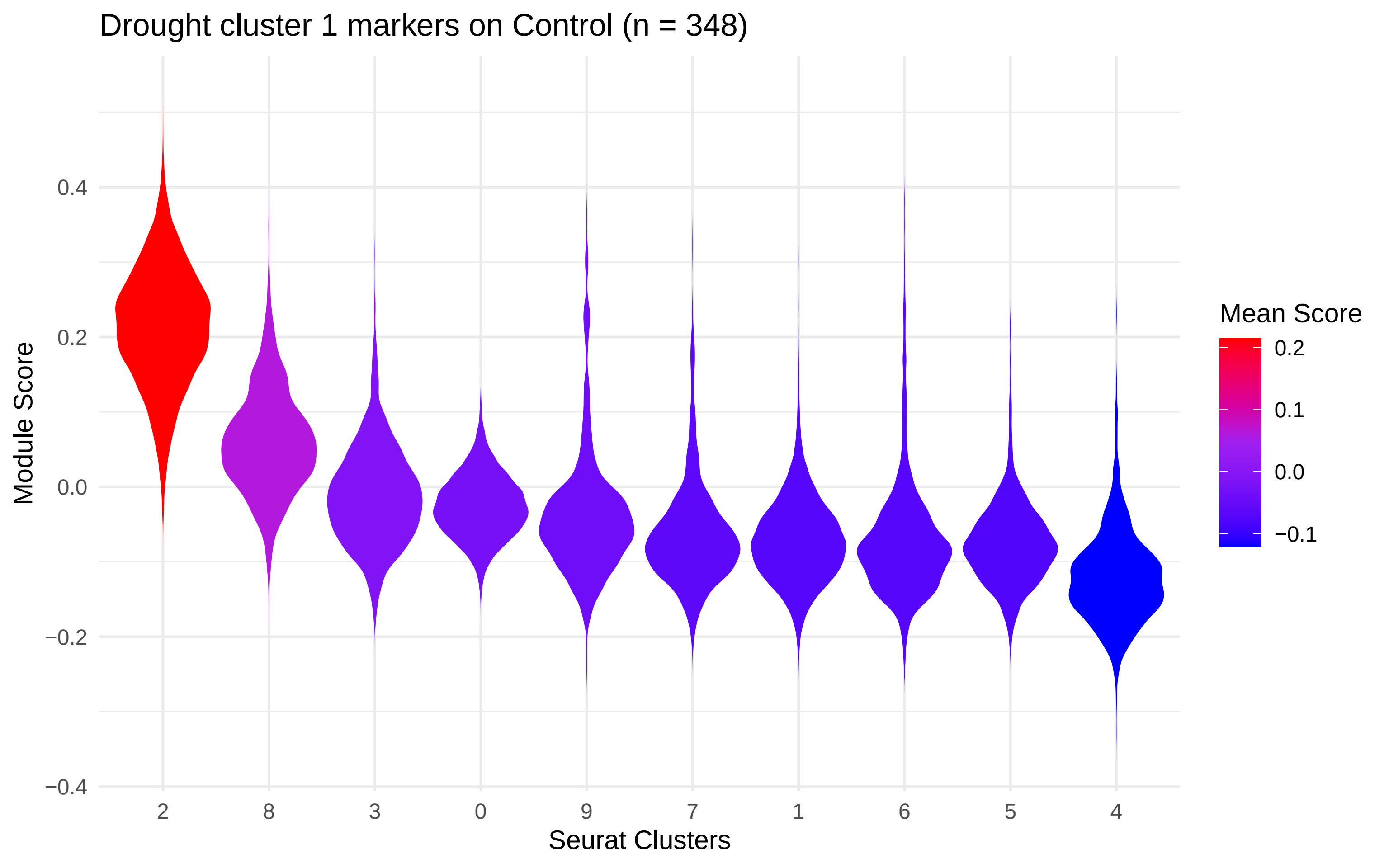
Drought cluster 1 markers on Control (n = 348)
Module Score
0.4
0.2
0.0
−0.2
−0.4
2
8
3
0
9
7
1
6
5
4
Seurat Clusters
Mean Score
0.2
0.1
0.0
−0.1

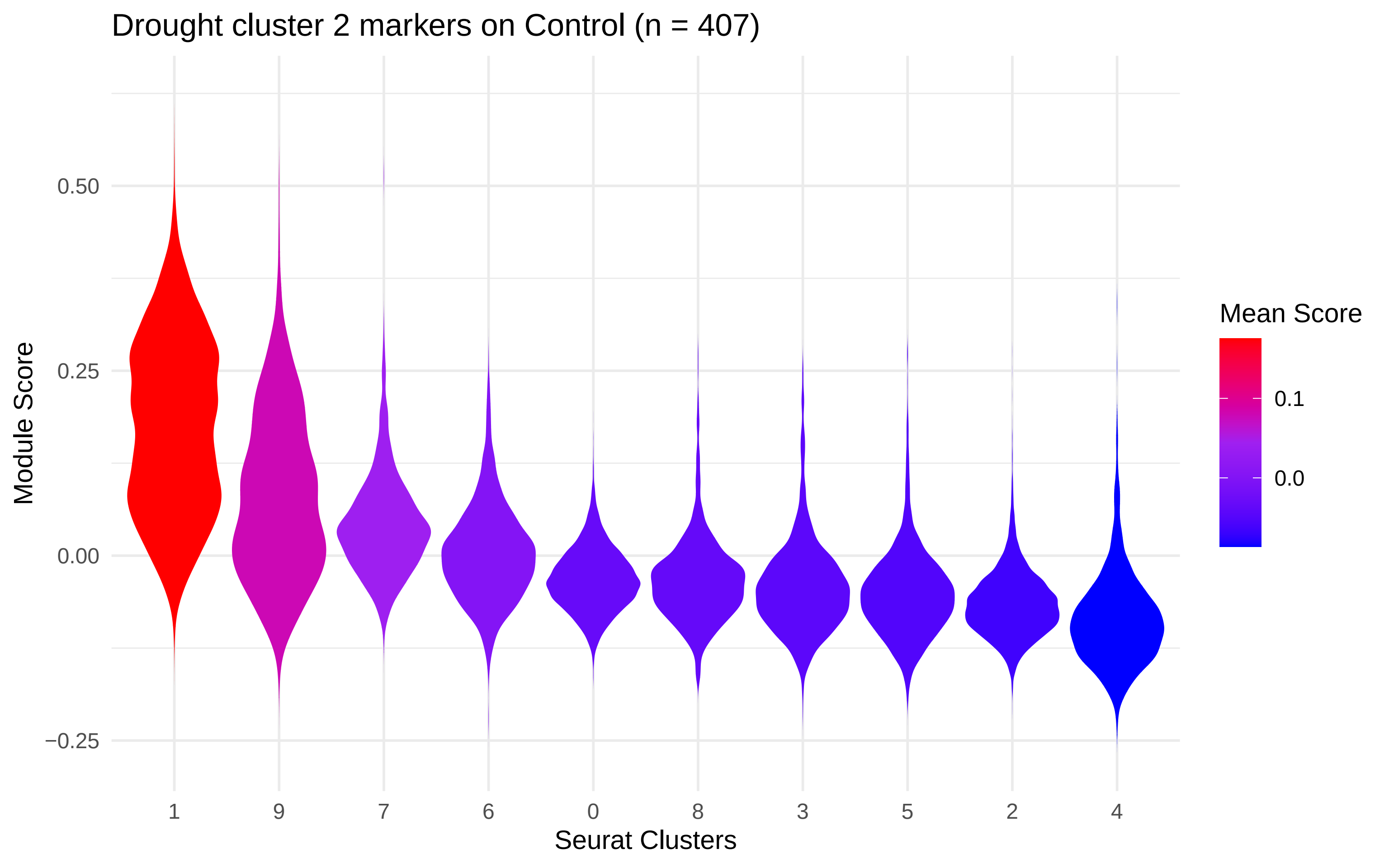

Drought cluster 2 markers on Control (n = 407)
Module Score
0.50
0.25
0.00
−0.25
1
9
7
6
0
8
3
5
2
4
Seurat Clusters
Mean Score
0.1
0.0

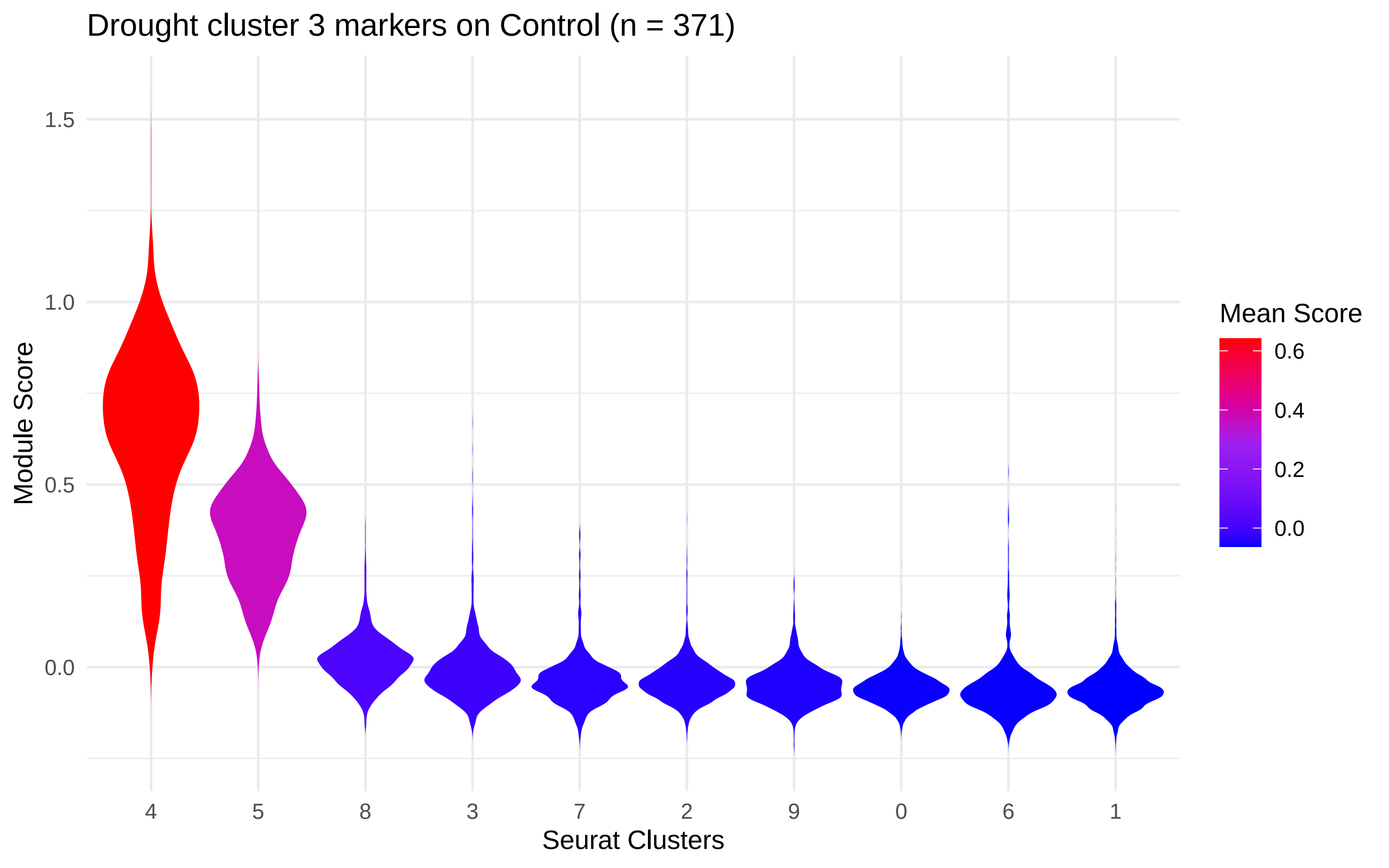
Drought cluster 3 markers on Control (n = 371)
Module Score
1.5
1.0
0.5
0.0
4
5
8
3
7
2
9
0
6
1
Seurat Clusters
Mean Score
0.6
0.4
0.2
0.0

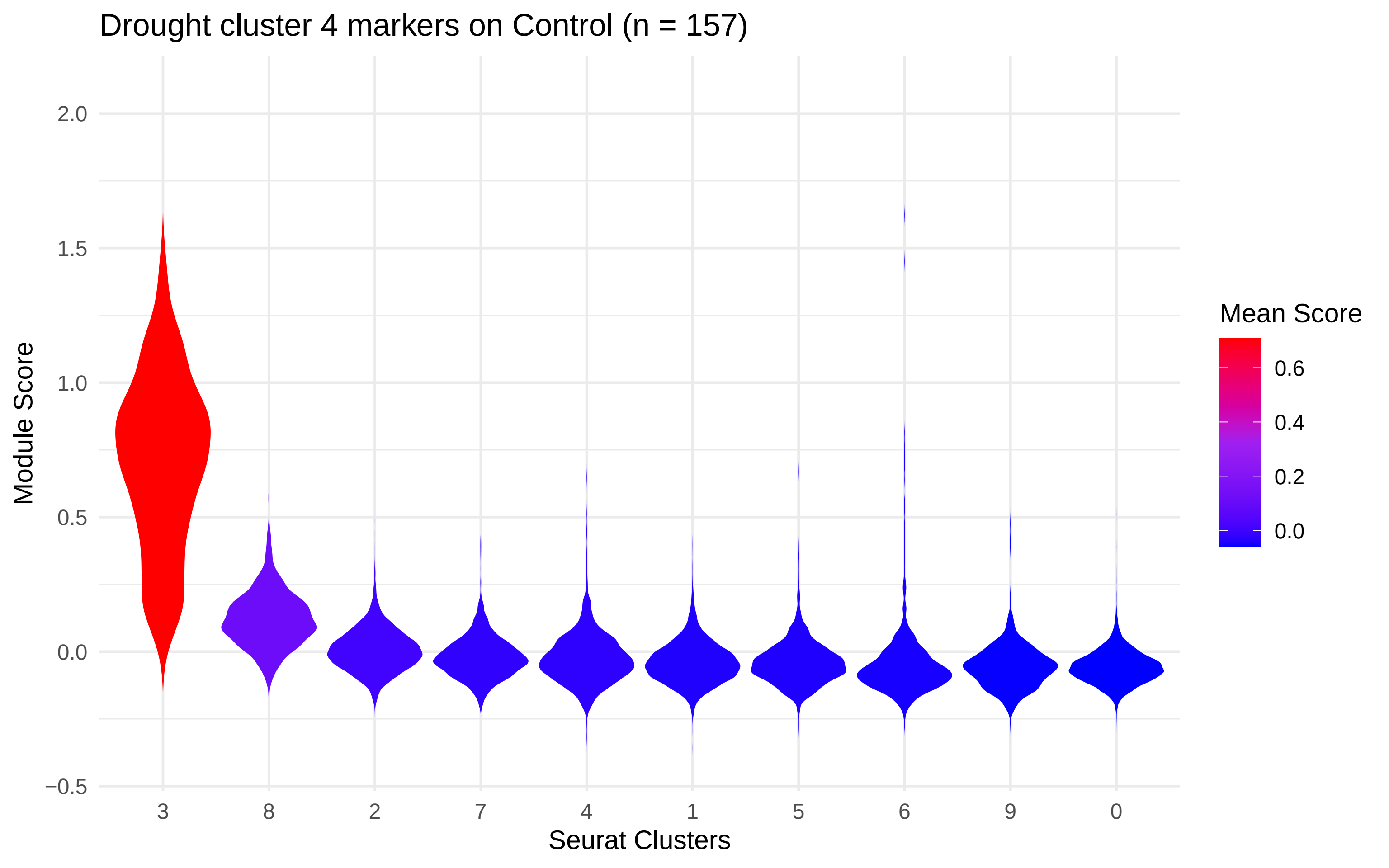
Drought cluster 4 markers on Control (n = 157)
Module Score
2.0
1.5
1.0
0.5
0.0
−0.5
3
8
2
7
4
1
5
6
9
0
Seurat Clusters
Mean Score
0.6
0.4
0.2
0.0

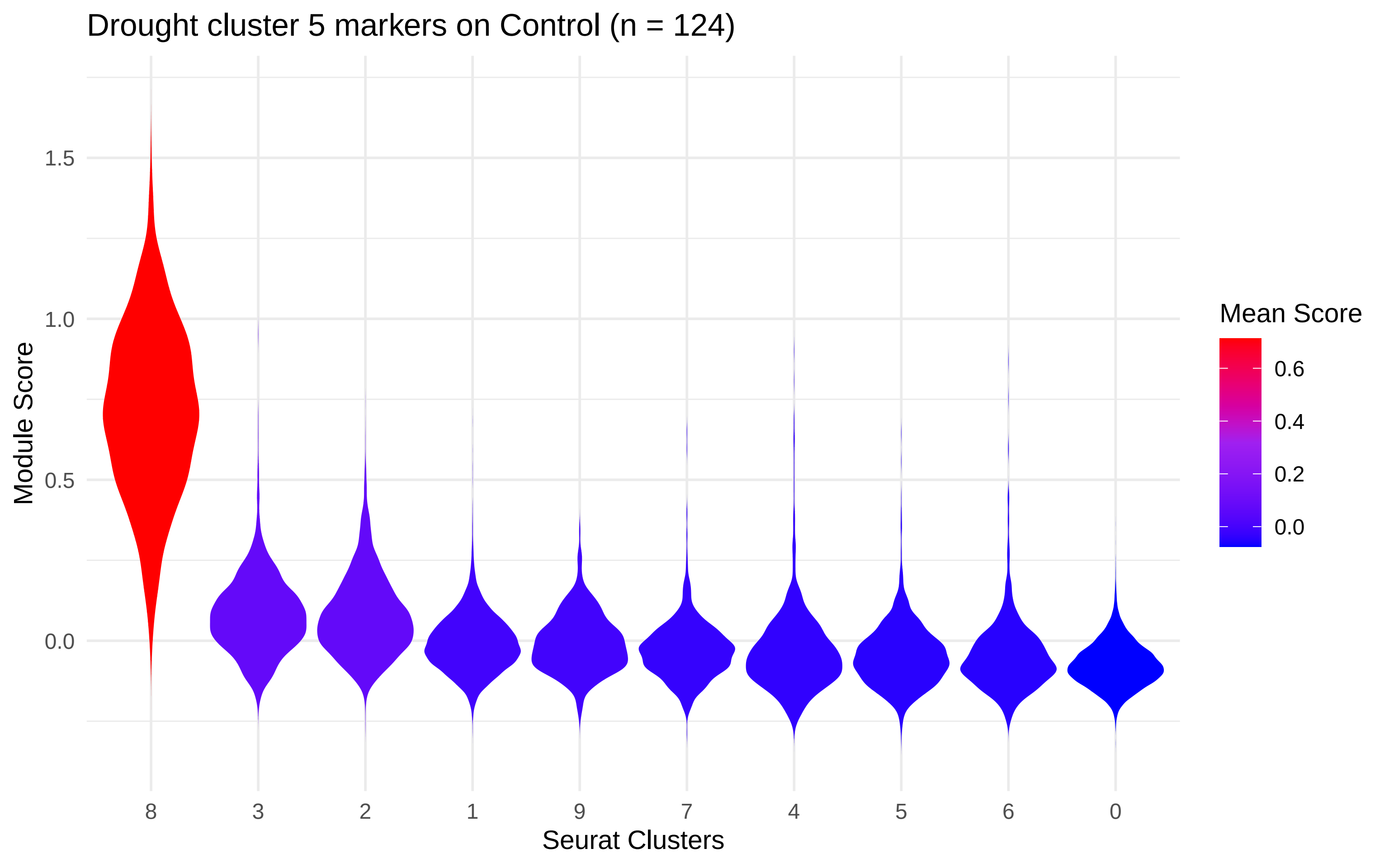

Drought cluster 5 markers on Control (n = 124)
Module Score
1.5
1.0
0.5
0.0
8
3
2
1
9
7
4
5
6
0
Seurat Clusters
Mean Score
0.6
0.4
0.2
0.0

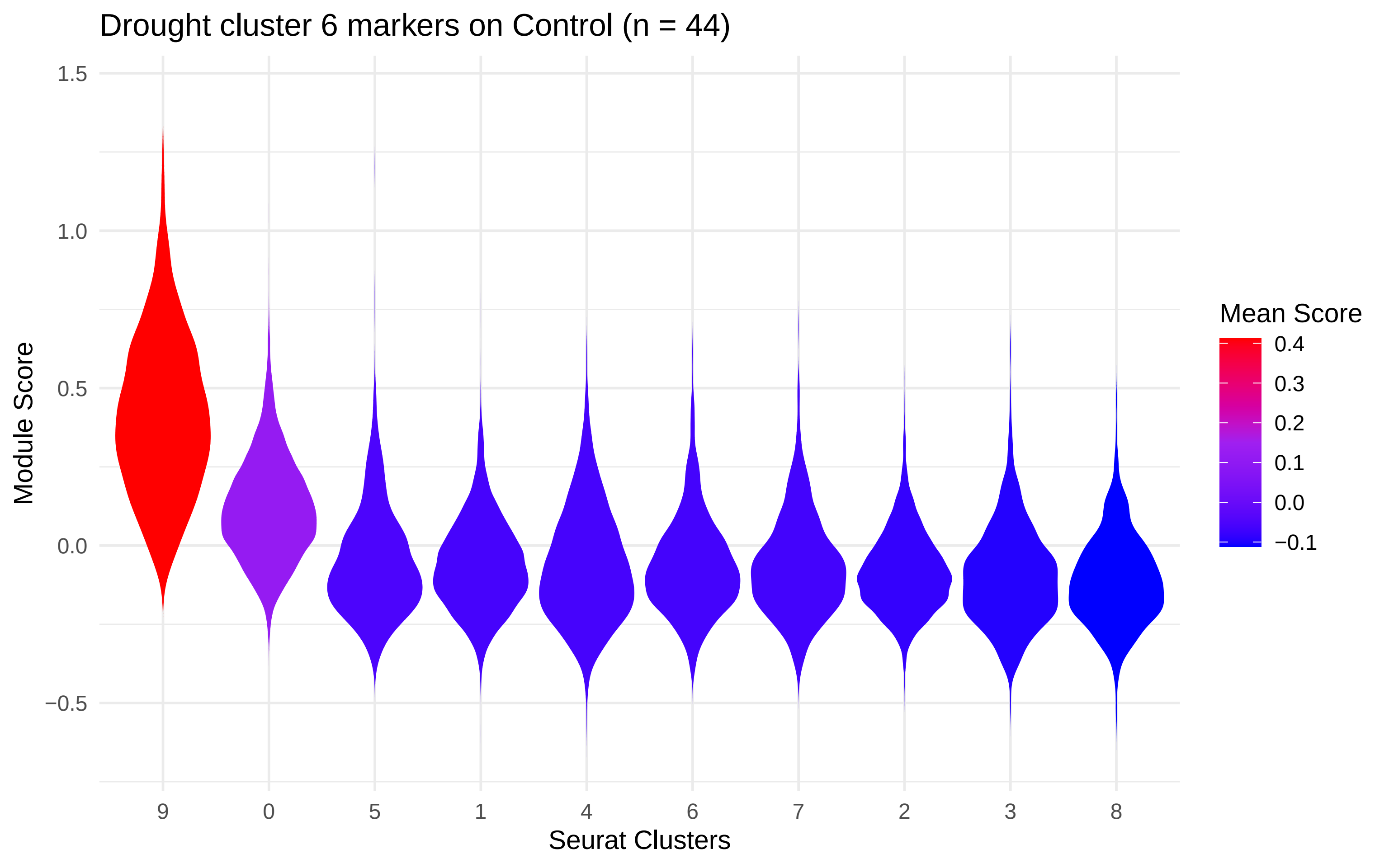

Drought cluster 6 markers on Control (n = 44)
Module Score
1.5
1.0
0.5
0.0
−0.5
9
0
5
1
4
6
7
2
3
8
Seurat Clusters
Mean Score
0.4
0.3
0.2
0.1
0.0
−0.1

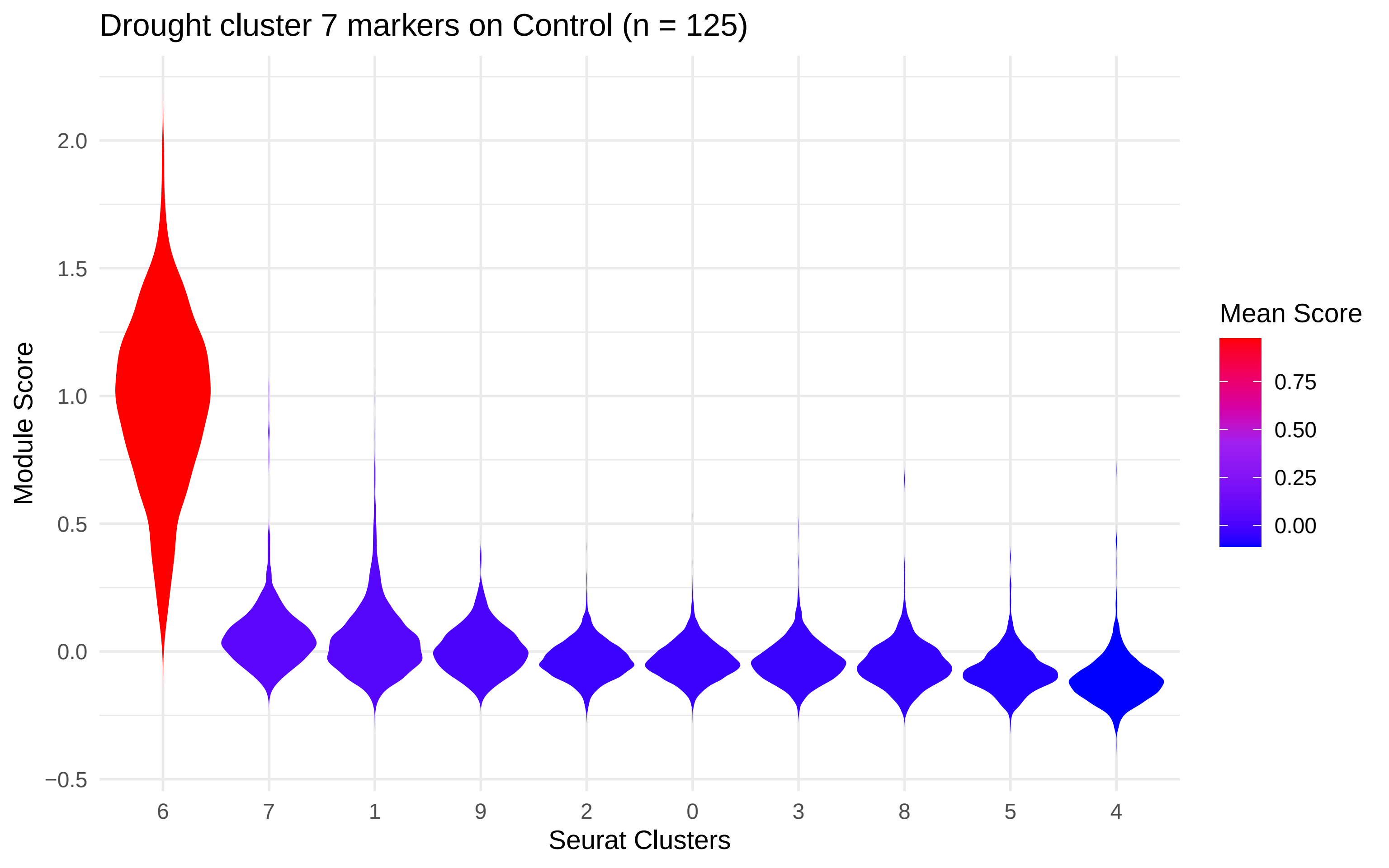
Drought cluster 7 markers on Control (n = 125)
Module Score
2.0
1.5
1.0
0.5
0.0
−0.5
6
7
1
9
2
0
3
8
5
4
Seurat Clusters
Mean Score
0.75
0.50
0.25
0.00

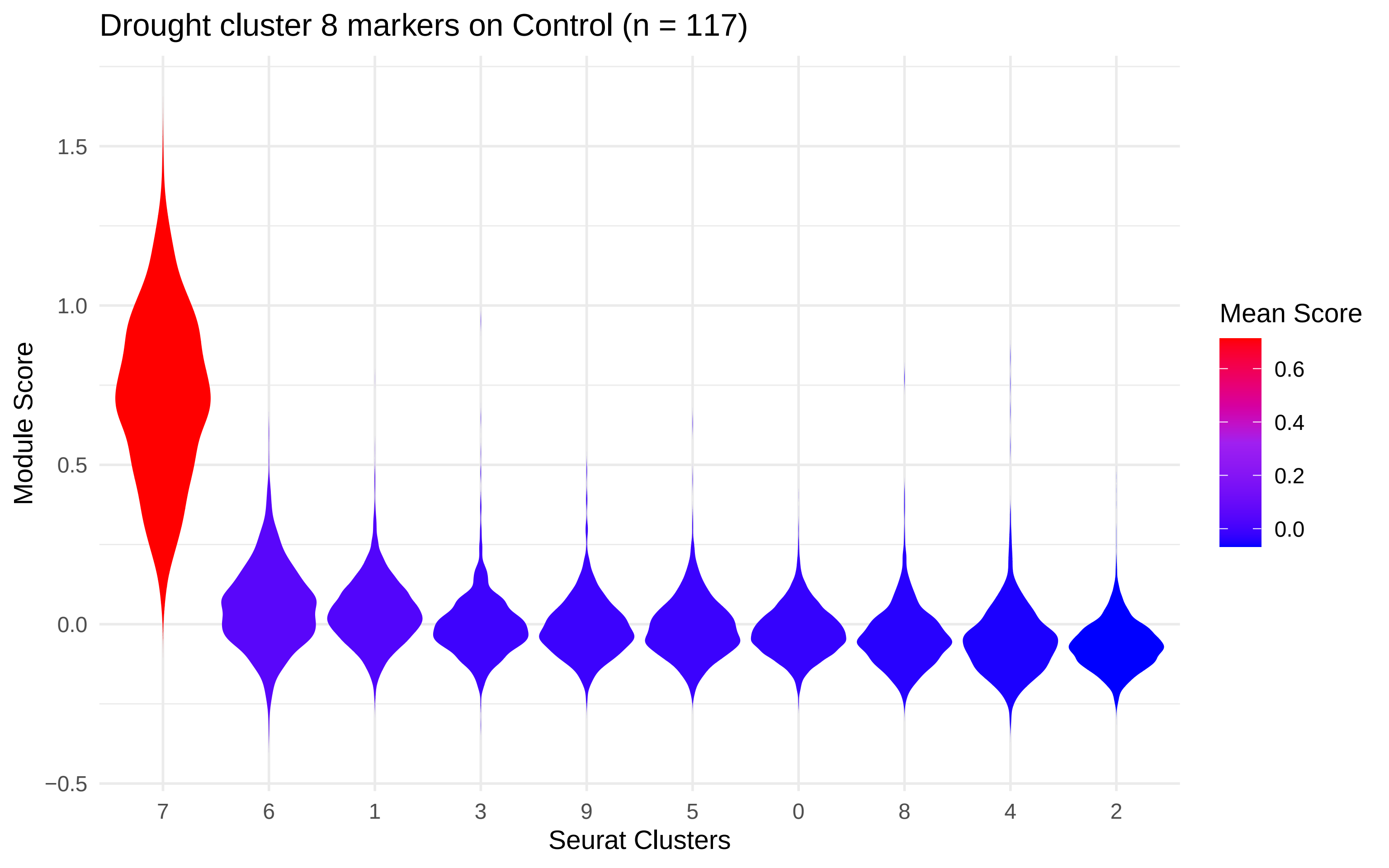

Drought cluster 8 markers on Control (n = 117)
Module Score
1.5
1.0
0.5
0.0
−0.5
7
6
1
3
9
5
0
8
4
2
Seurat Clusters
Mean Score
0.6
0.4
0.2
0.0
