## Supplementary material for "Single-cell-level response to drought in *Sorghum bicolor* reveals novel targets for improving water use efficiency": Cell type marker gene mapping, control

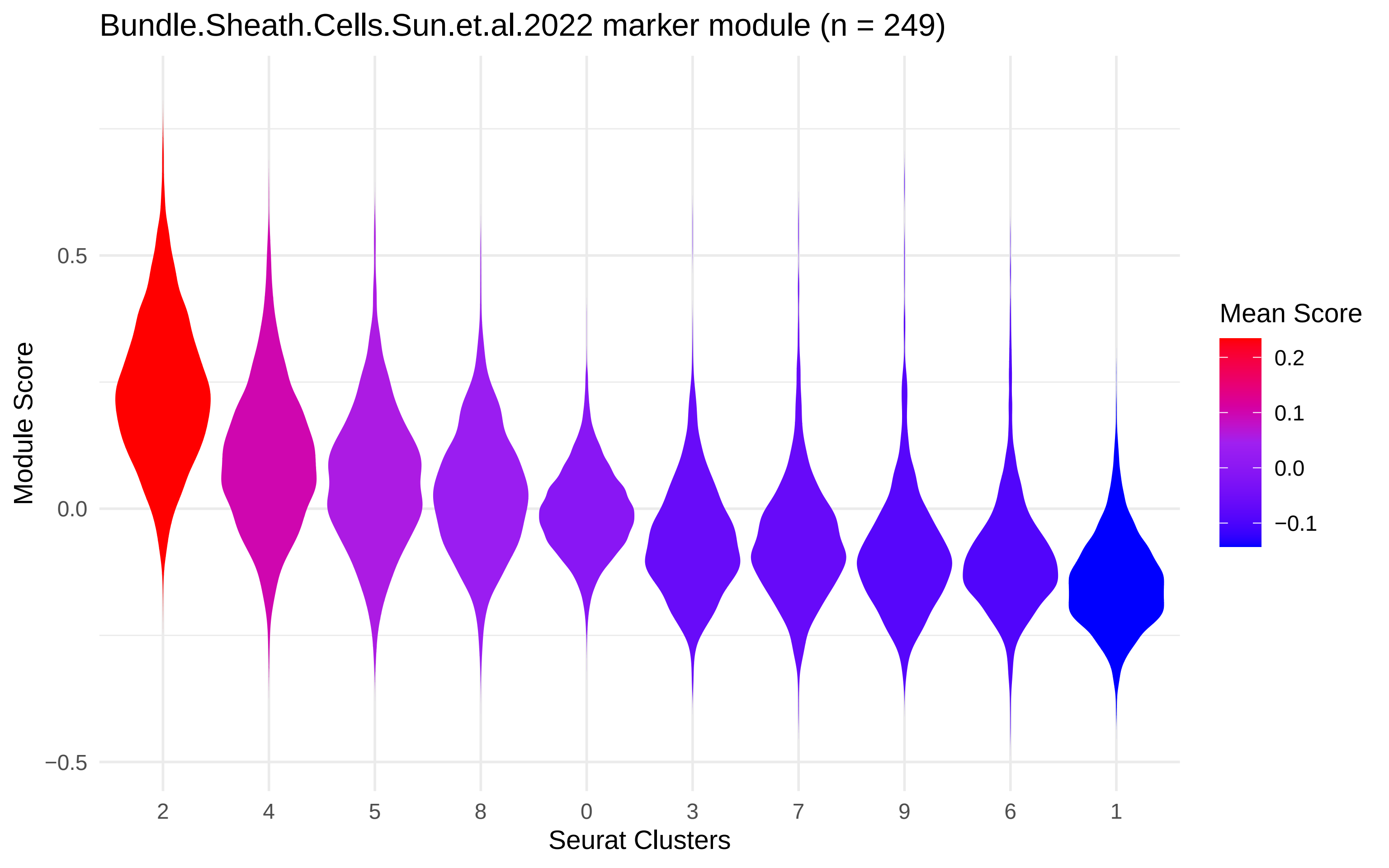

Bundle.Sheath.Cells.Sun.et.al.2022 marker module (n = 249)
Module Score
0.5
0.0
−0.5
2
4
5
8
0
3
7
9
6
1
Seurat Clusters
Mean Score
0.2
0.1
0.0
−0.1

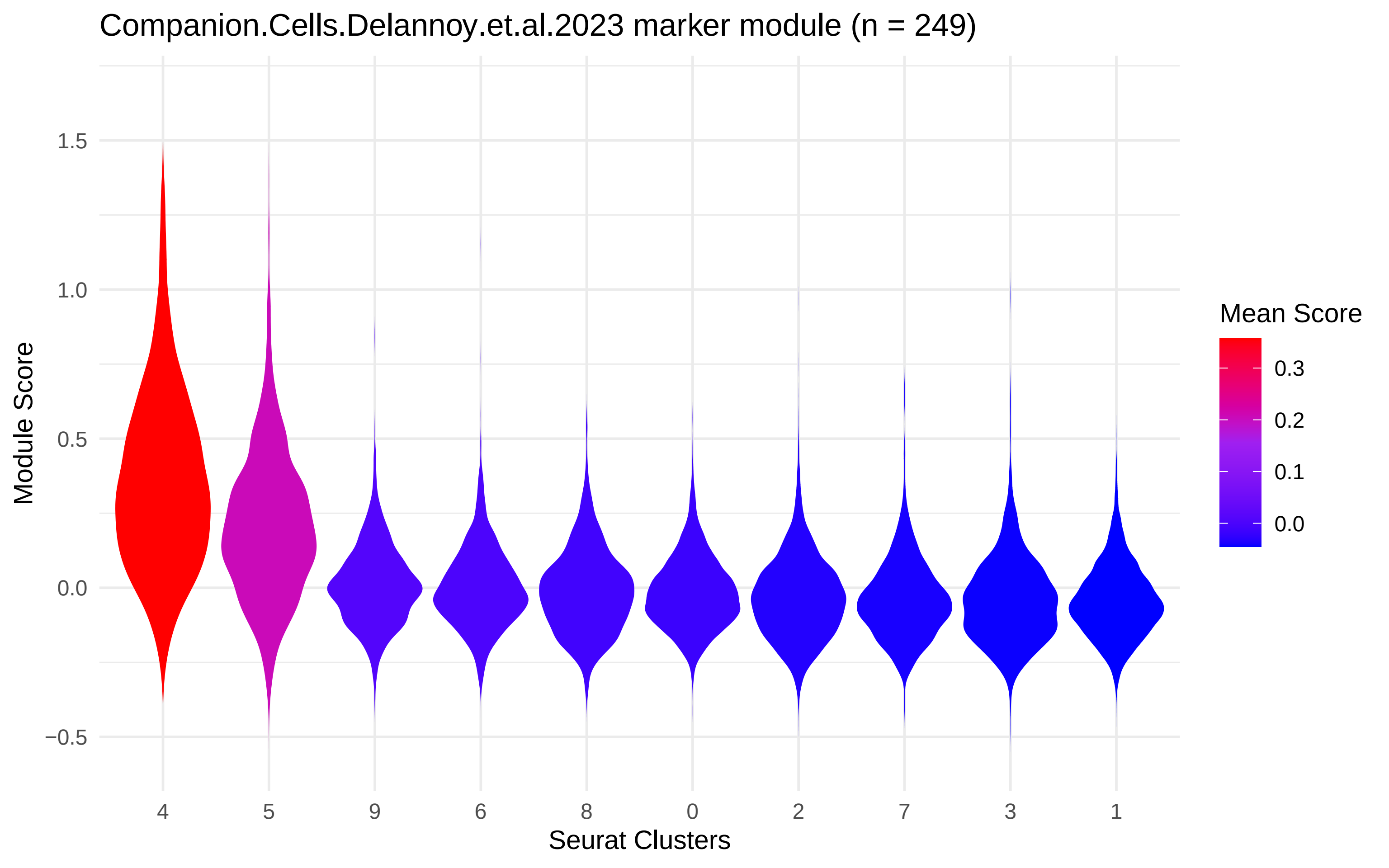

Companion.Cells.Delannoy.et.al.2023 marker module (n = 249)
Module Score
1.5
1.0
0.5
0.0
−0.5
4
5
9
6
8
0
2
7
3
1
Seurat Clusters
Mean Score
0.3
0.2
0.1
0.0

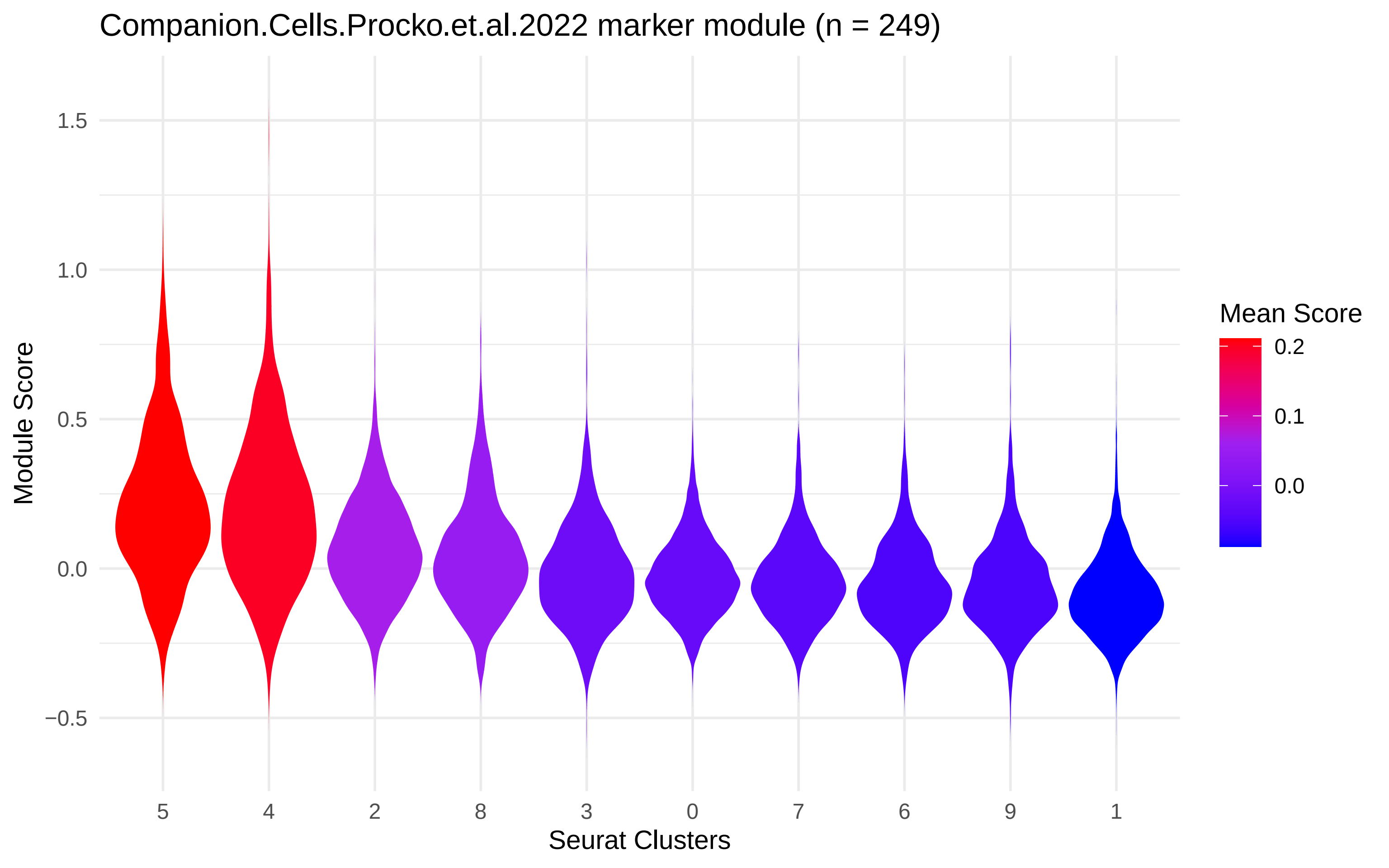

Companion.Cells.Procko.et.al.2022 marker module (n = 249)
Module Score
1.5
1.0
0.5
0.0
−0.5
5
4
2
8
3
0
7
6
9
1
Seurat Clusters
Mean Score
0.2
0.1
0.0

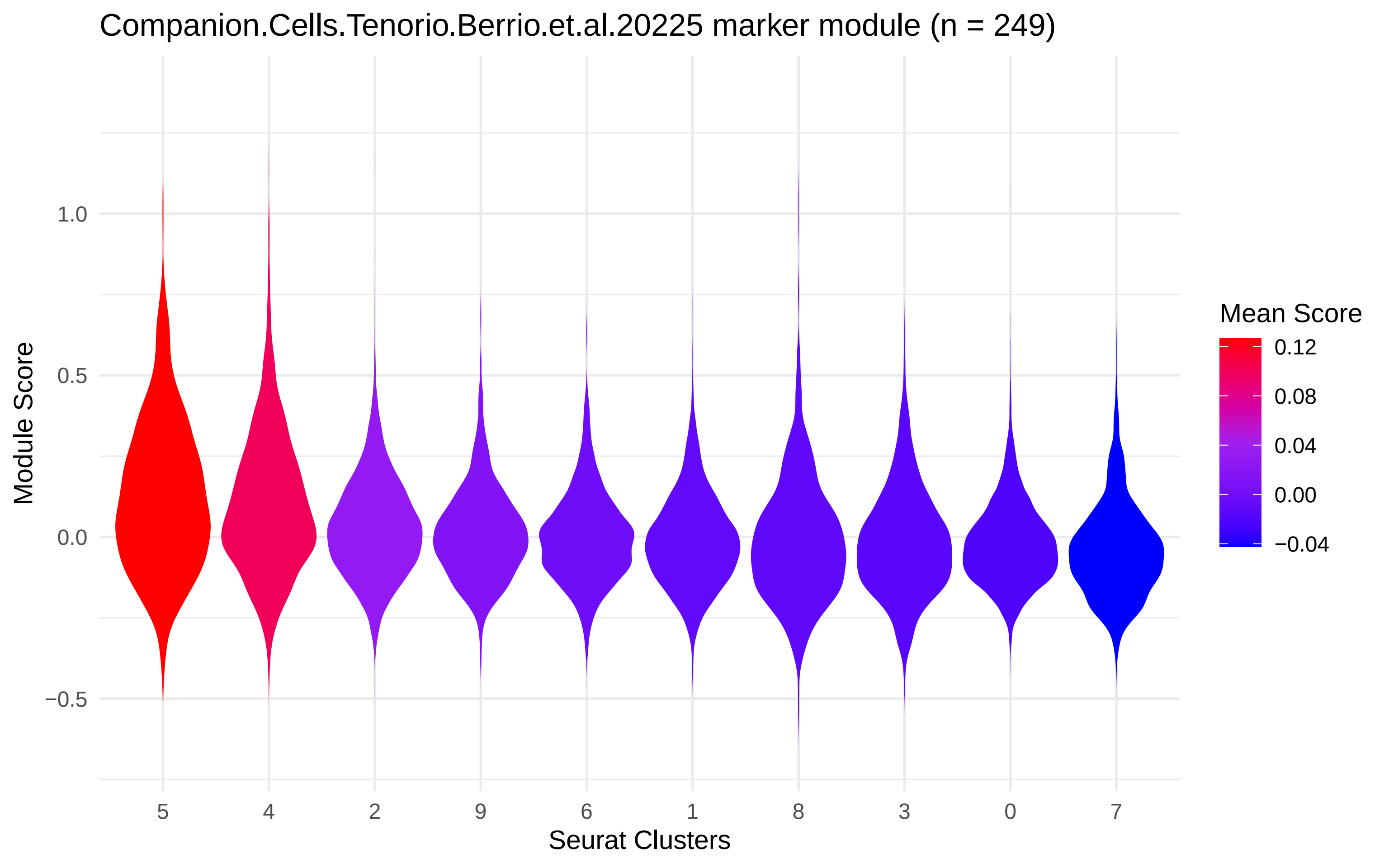
Companion.Cells.Tenorio.Berrio.et.al.20225 marker module (n = 249)
Module Score
1.0
0.5
0.0
−0.5
5
4
2
9
6
1
8
3
0
7
Seurat Clusters
Mean Score
0.12
0.08
0.04
0.00
−0.04

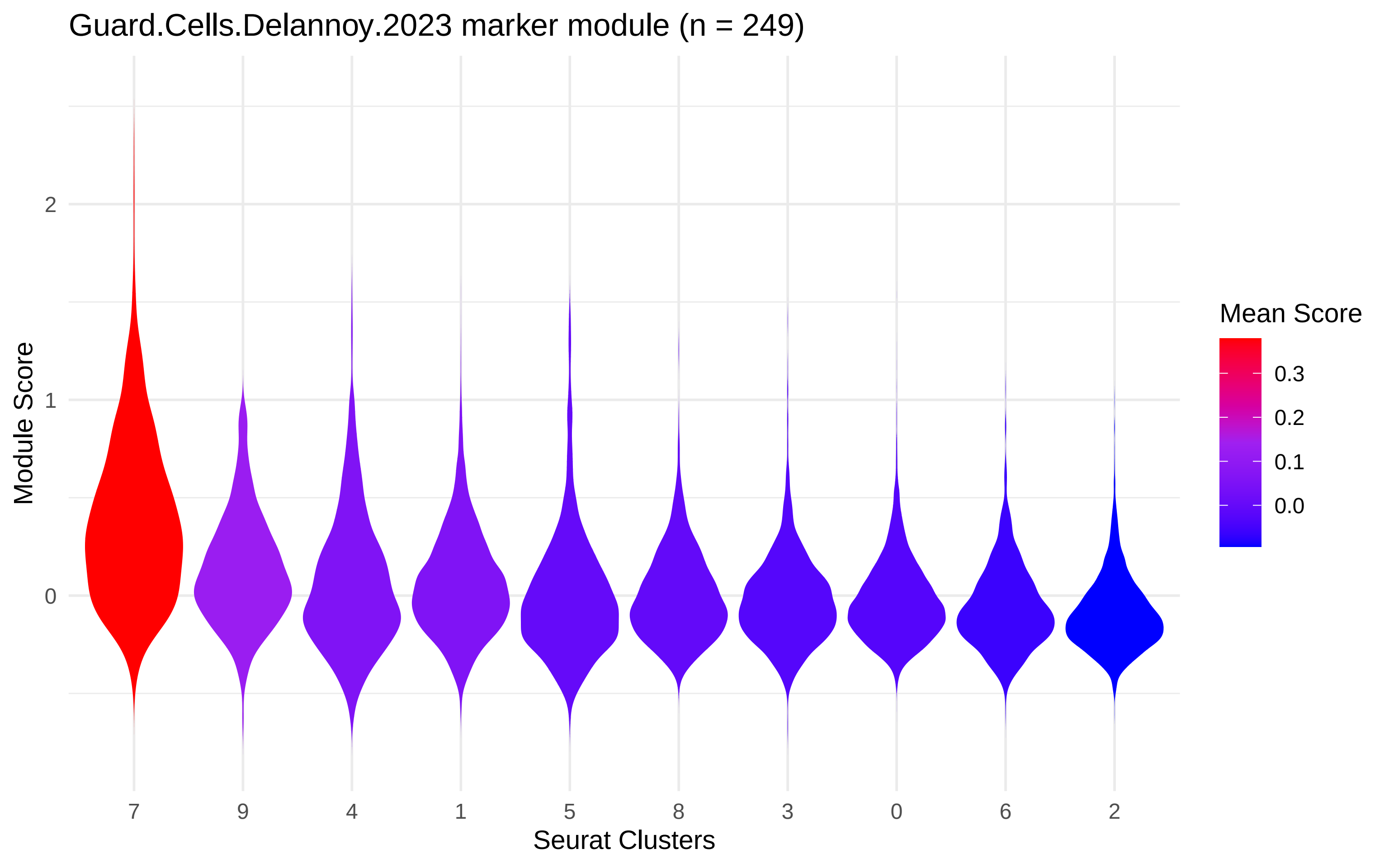

Guard.Cells.Delannoy.2023 marker module (n = 249)
Module Score
2
1
0
7
9
4
1
5
8
3
0
6
2
Seurat Clusters
Mean Score
0.3
0.2
0.1
0.0

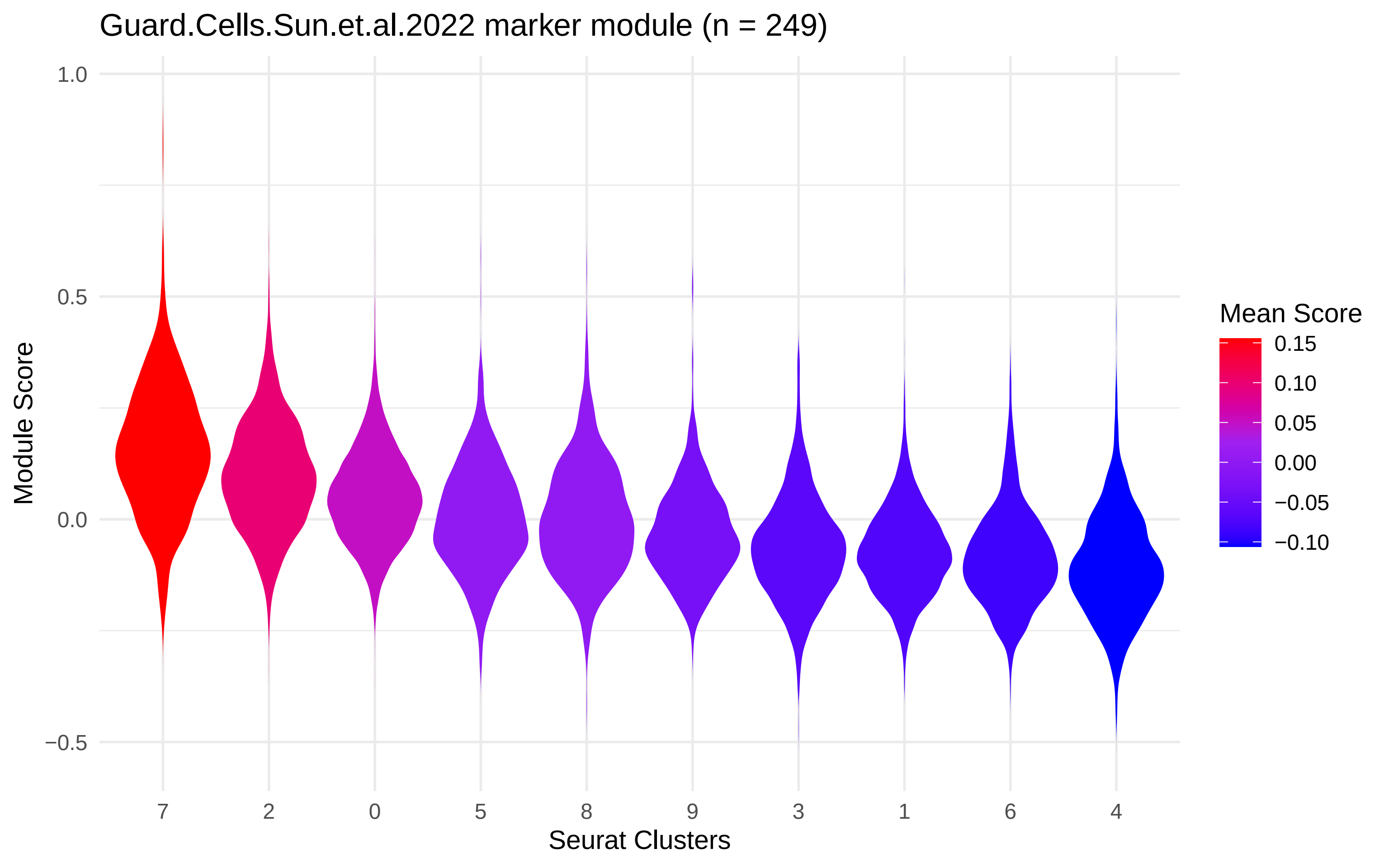

Guard.Cells.Sun.et.al.2022 marker module (n = 249)
Module Score
1.0
0.5
0.0
−0.5
7
2
0
5
8
9
3
1
6
4
Seurat Clusters
Mean Score
0.15
0.10
0.05
0.00
−0.05
−0.10

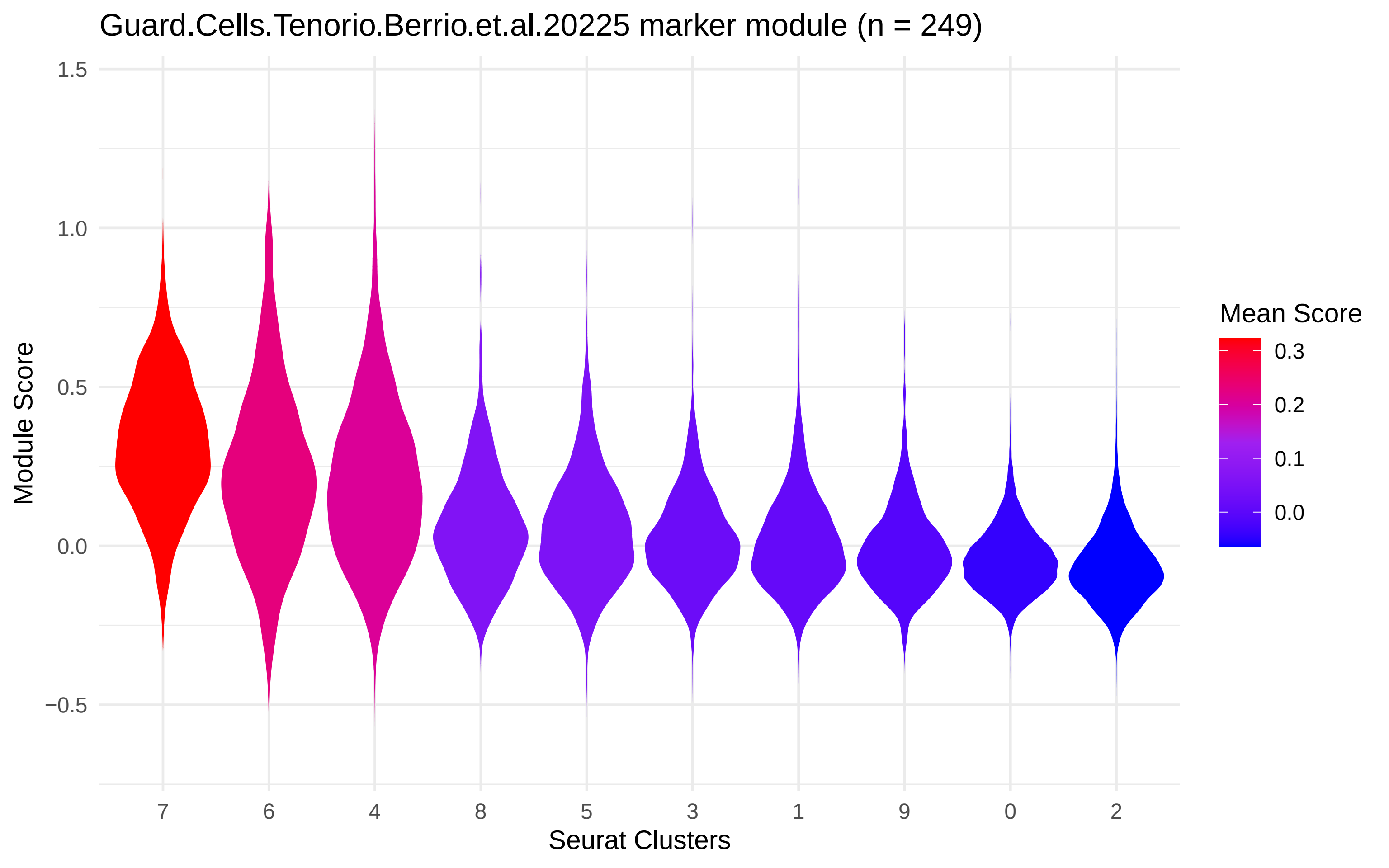

Guard.Cells.Tenorio.Berrio.et.al.20225 marker module (n = 249)
Module Score
1.5
1.0
0.5
0.0
−0.5
7
6
4
8
5
3
1
9
0
2
Seurat Clusters
Mean Score
0.3
0.2
0.1
0.0

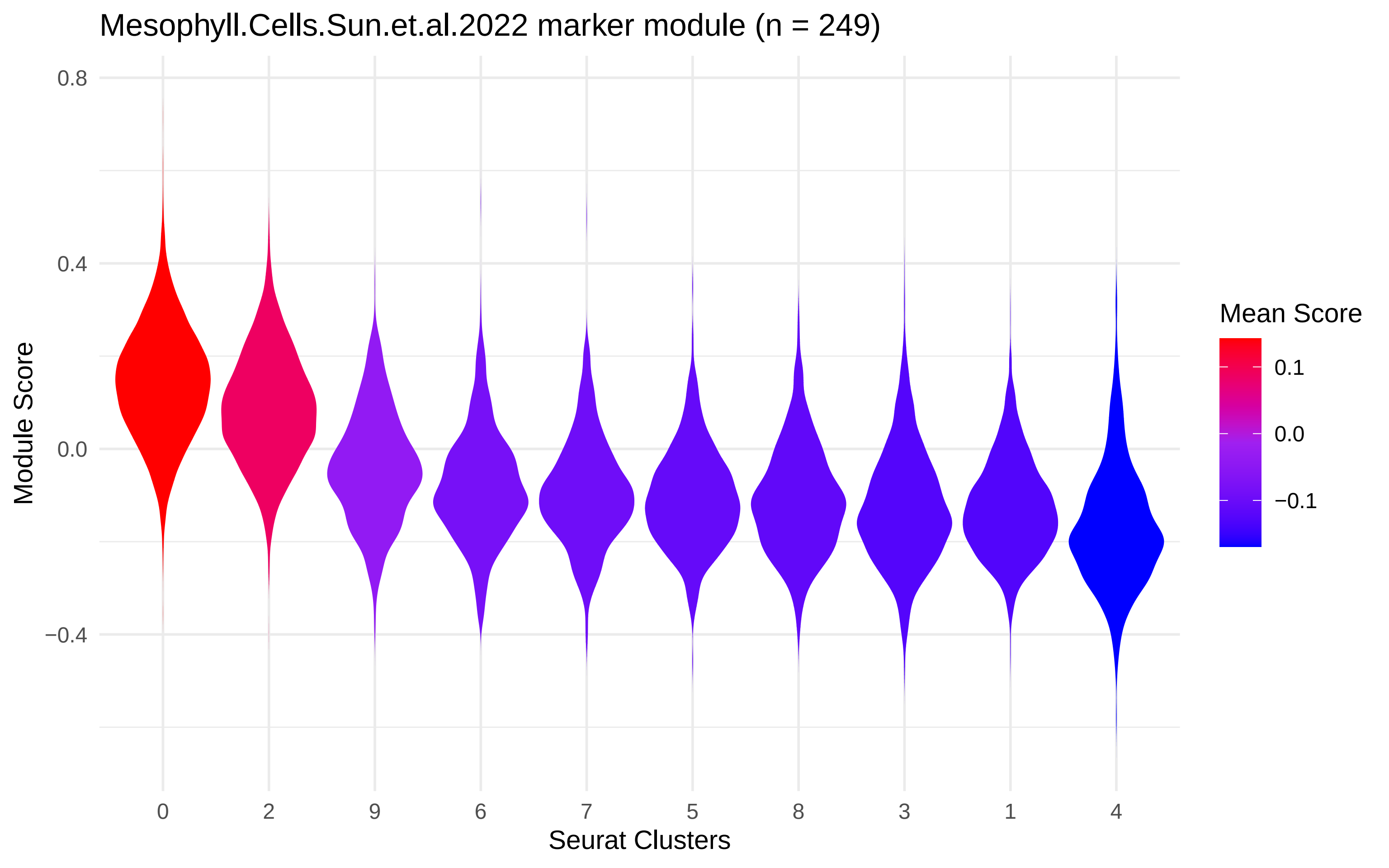

Mesophyll.Cells.Sun.et.al.2022 marker module (n = 249)
Module Score
0.8
0.4
0.0
−0.4
0
2
9
6
7
5
8
3
1
4
Seurat Clusters
Mean Score
0.1
0.0
−0.1

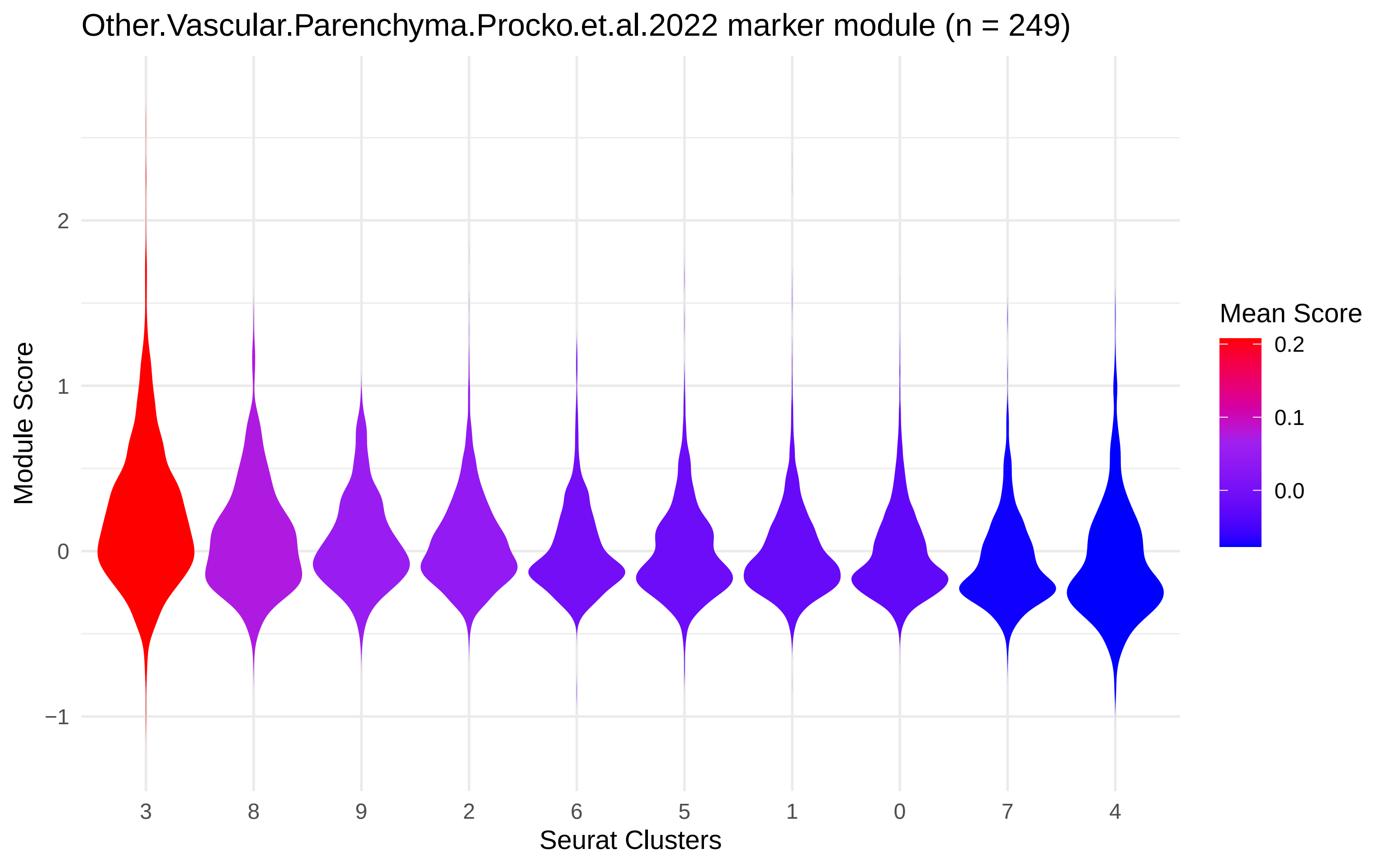

Other.Vascular.Parenchyma.Procko.et.al.2022 marker module (n = 249)
Module Score
2
1
0
−1
3
8
9
2
6
5
1
0
7
4
Seurat Clusters
Mean Score
0.2
0.1
0.0

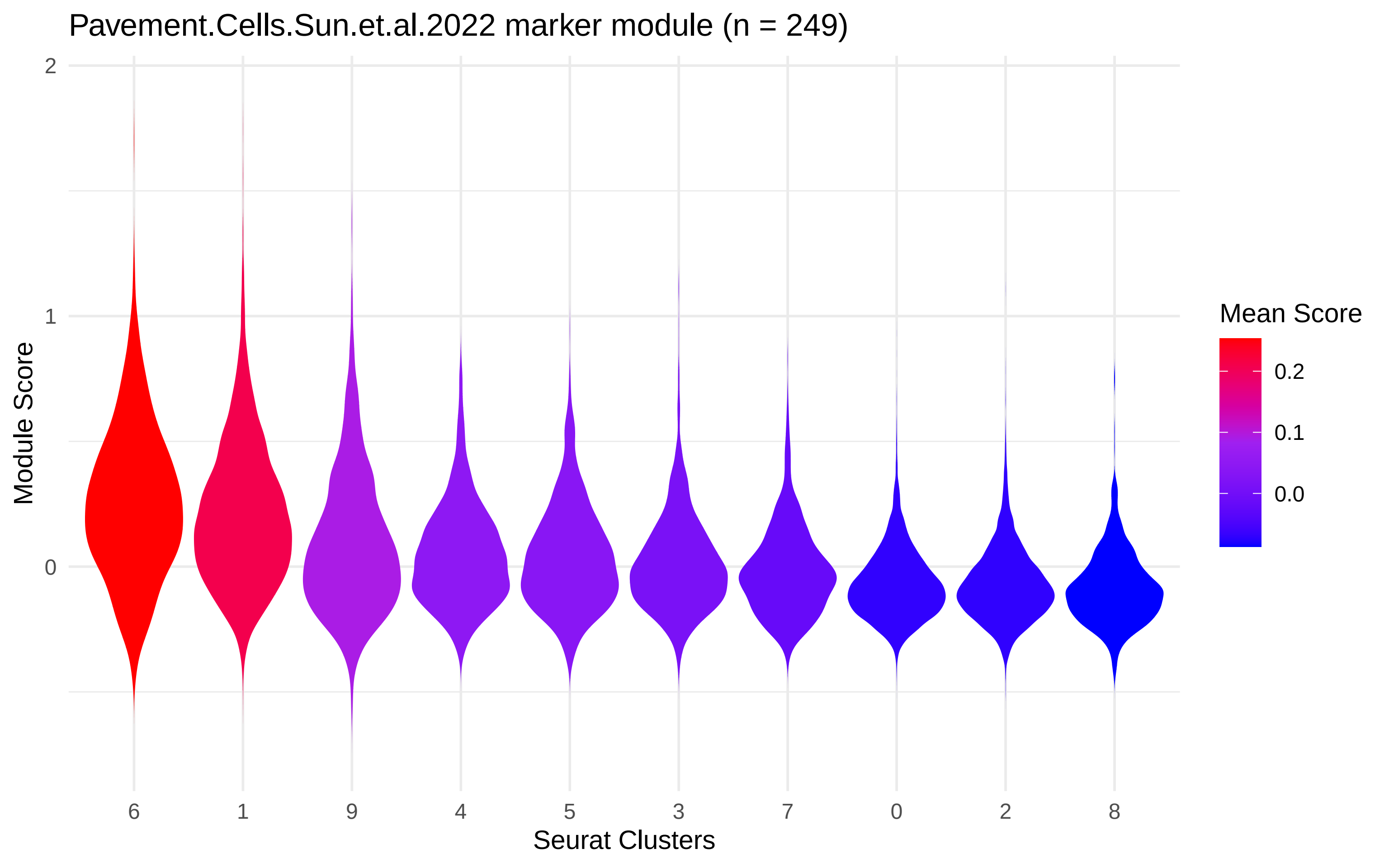

Pavement.Cells.Sun.et.al.2022 marker module (n = 249)
Module Score
2
1
0
Seurat Clusters
6
1
9
4
5
3
7
0
2
8
Mean Score
0.2
0.1
0.0

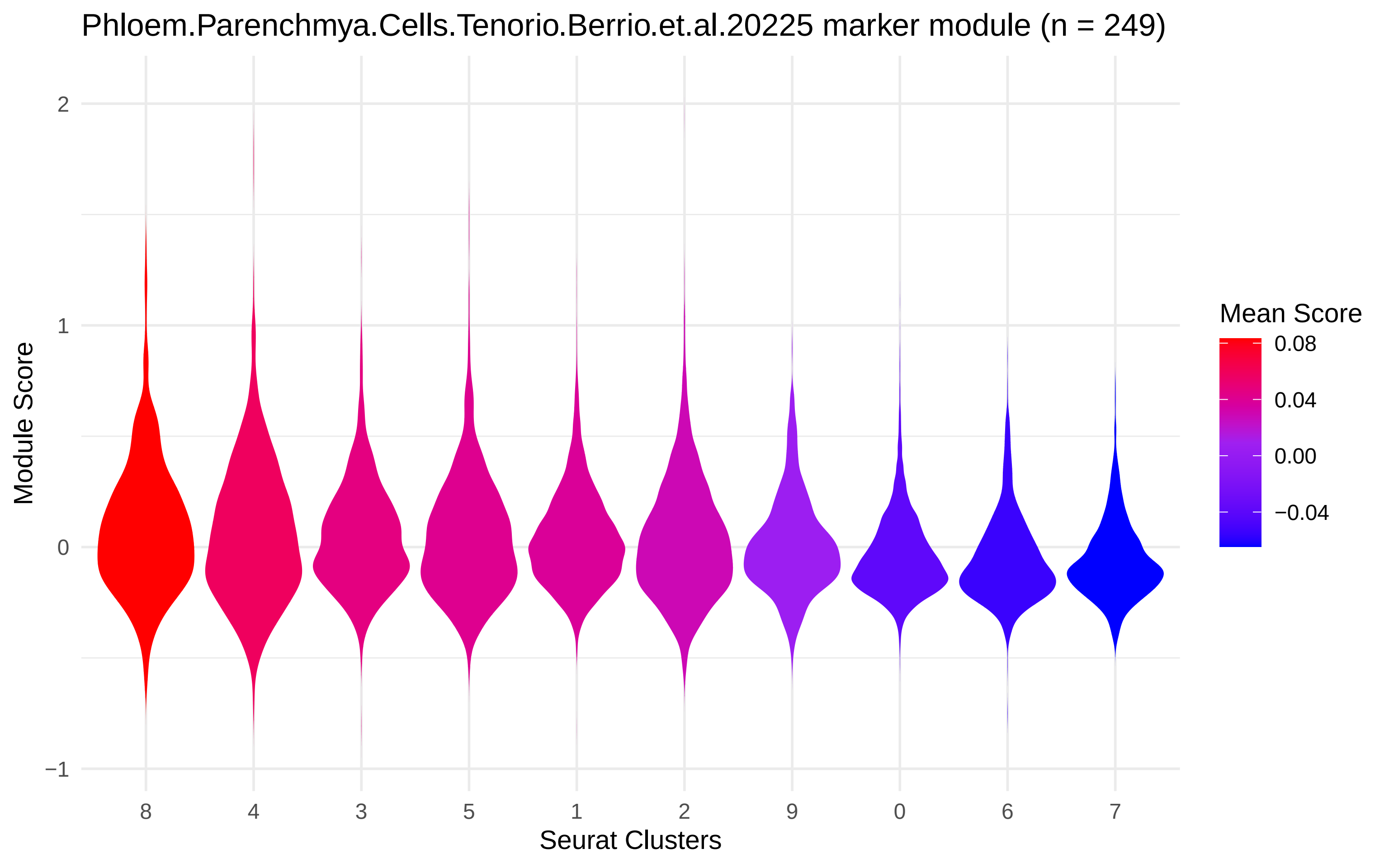

Phloem.Parenchmya.Cells.Tenorio.Berrio.et.al.20225 marker module (n = 249)
Module Score
2
1
0
−1
8
4
3
5
1
2
9
0
6
7
Seurat Clusters
Mean Score
0.08
0.04
0.00
−0.04

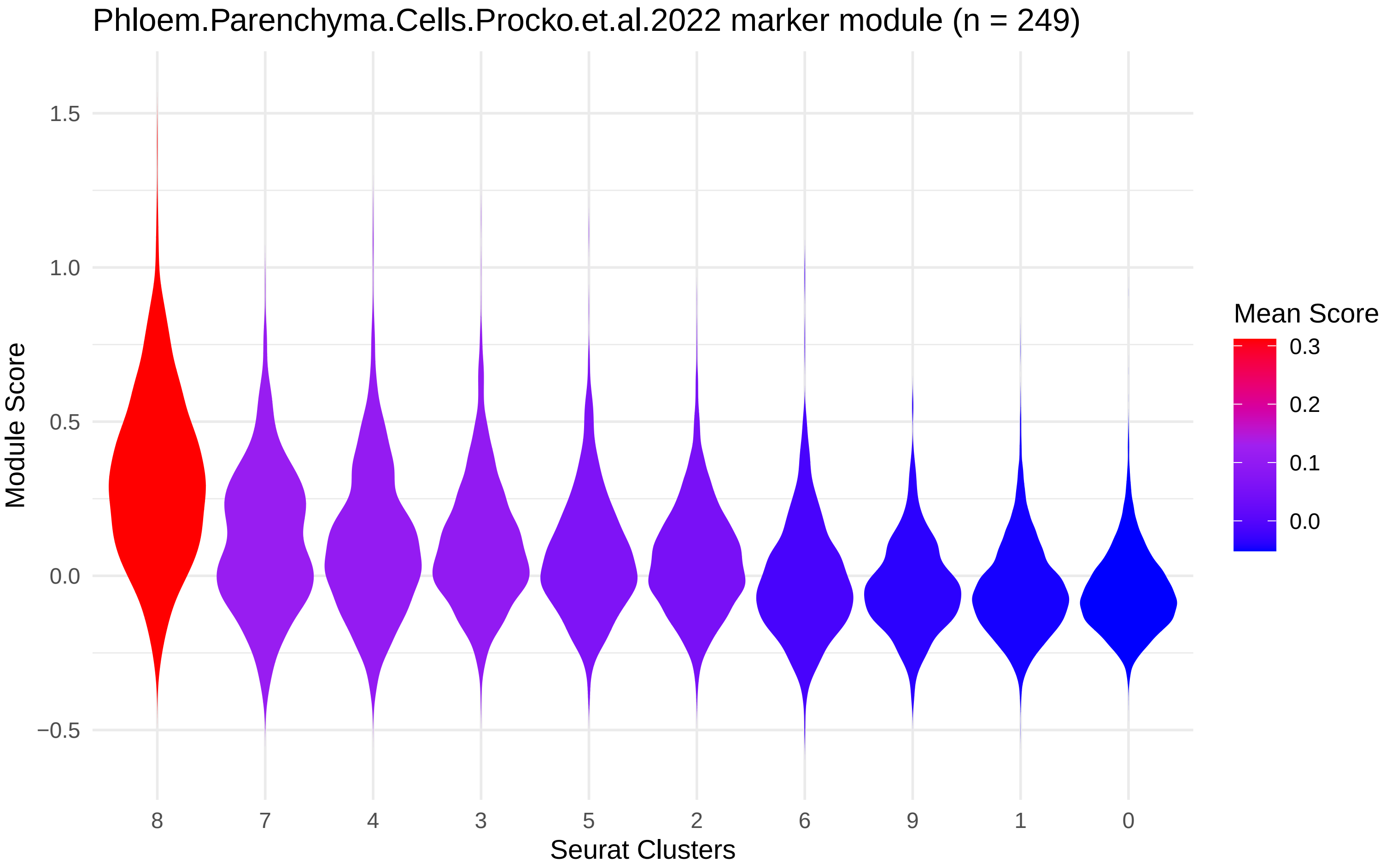
Phloem.Parenchyma.Cells.Procko.et.al.2022 marker module (n = 249)
Module Score
1.5
1.0
0.5
0.0
−0.5
8
7
4
3
5
2
6
9
1
0
Seurat Clusters
Mean Score
0.3
0.2
0.1
0.0

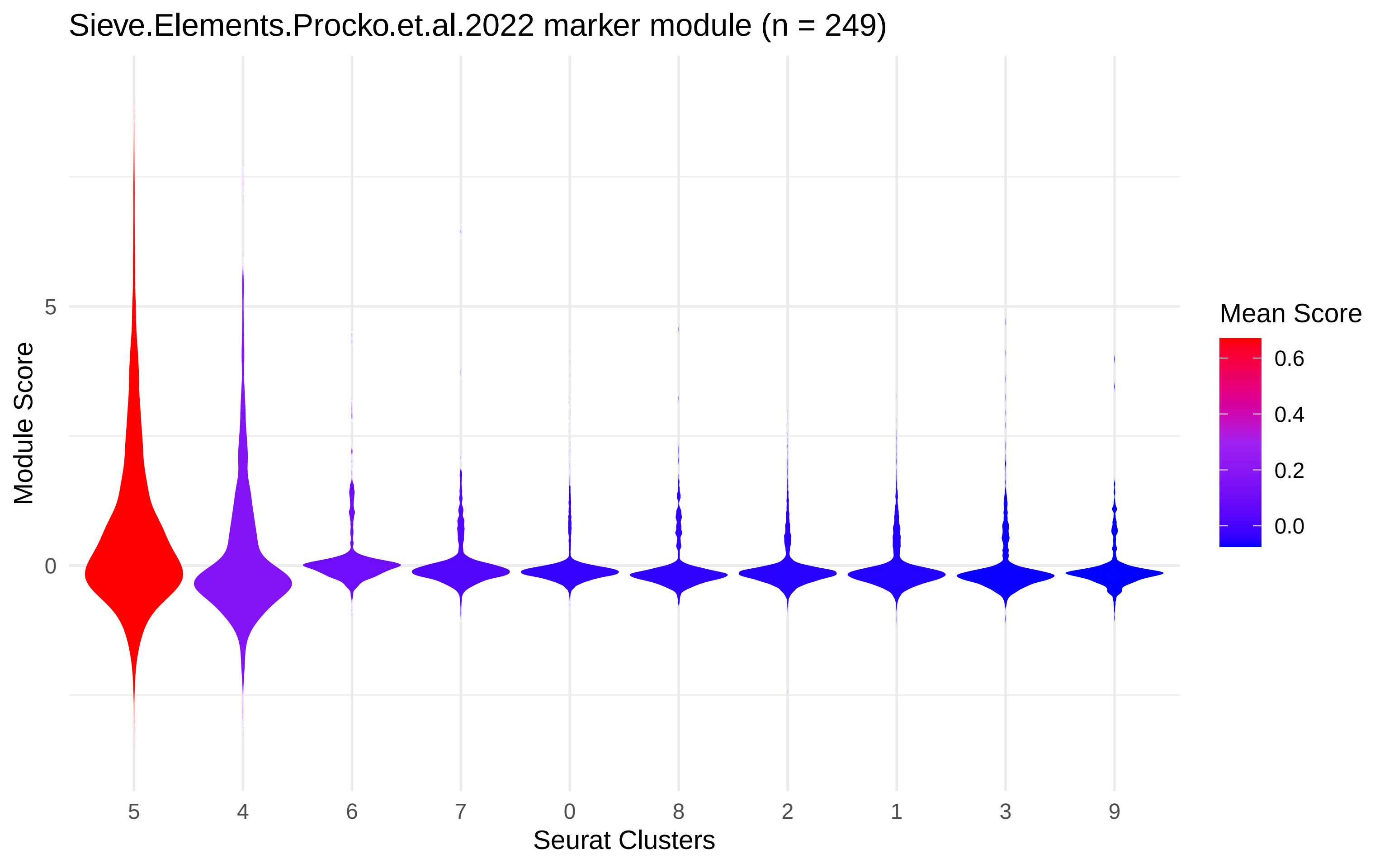
Sieve.Elements.Procko.et.al.2022 marker module (n = 249)
Module Score
5
0
5
4
6
7
0
8
2
1
3
9
Seurat Clusters
Mean Score
0.6
0.4
0.2
0.0

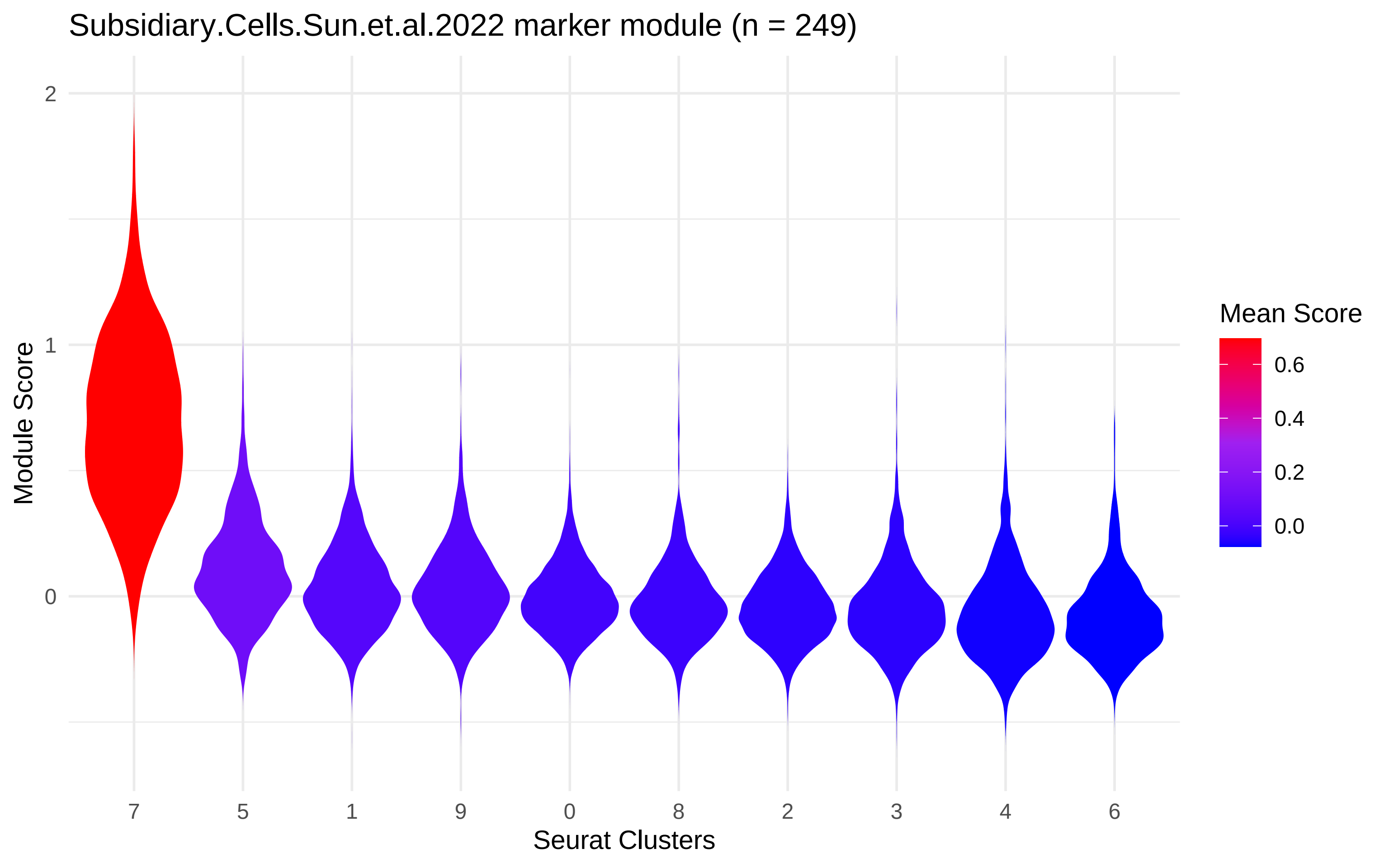

Subsidiary.Cells.Sun.et.al.2022 marker module (n = 249)
Module Score
2
1
0
7
5
1
9
0
8
2
3
4
6
Seurat Clusters
Mean Score
0.6
0.4
0.2
0.0

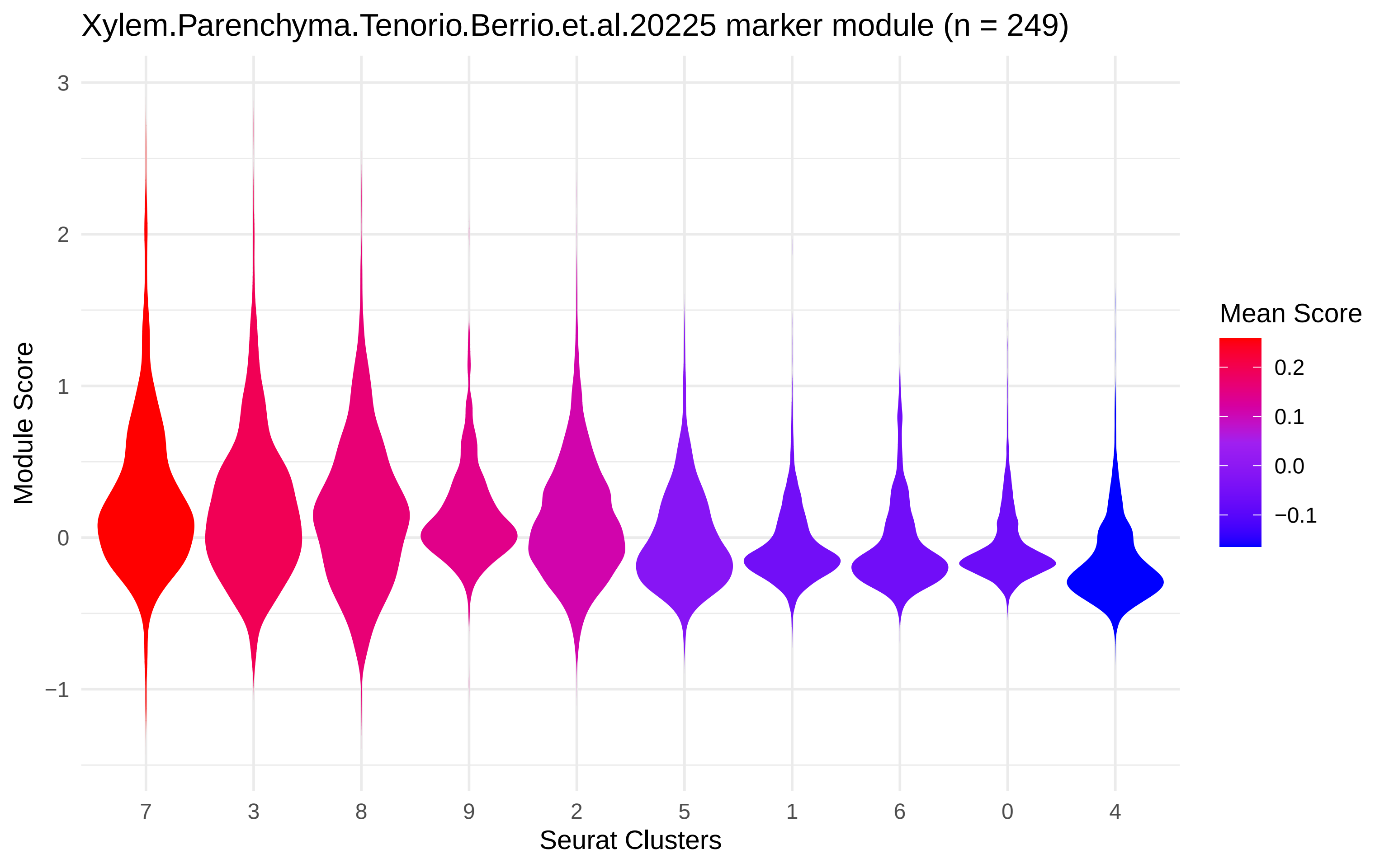

Xylem.Parenchyma.Tenorio.Berrio.et.al.20225 marker module (n = 249)
Module Score
3
2
1
0
−1
7
3
8
9
2
5
1
6
0
4
Seurat Clusters
Mean Score
0.2
0.1
0.0
−0.1
