## Supplementary material for "Single-cell-level response to drought in *Sorghum bicolor* reveals novel targets for improving water use efficiency": Cell type marker gene mapping, drought

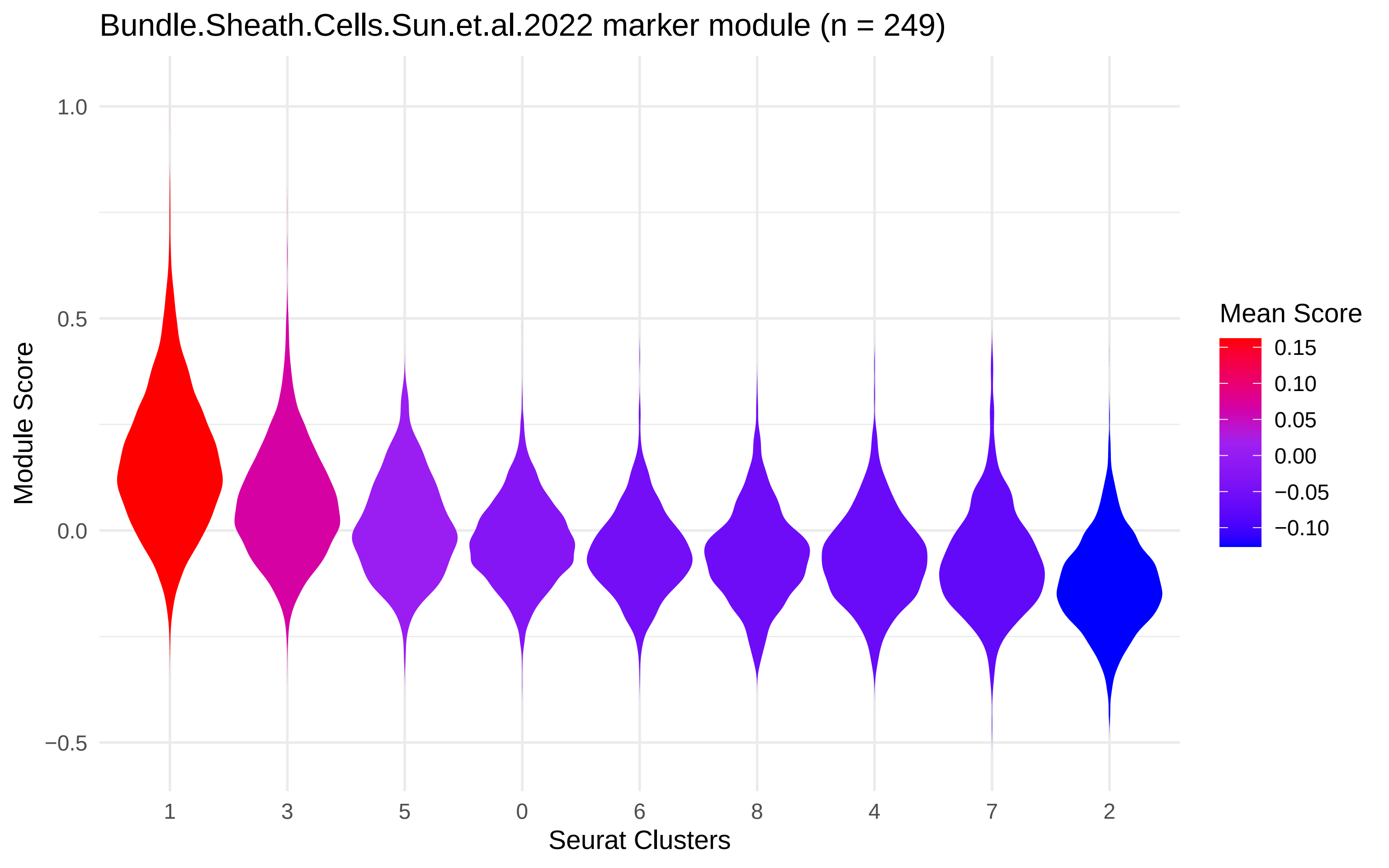

Bundle.Sheath.Cells.Sun.et.al.2022 marker module (n = 249)
Module Score
1.0
0.5
0.0
−0.5
1
3
5
0
6
8
4
7
2
Seurat Clusters
Mean Score
0.15
0.10
0.05
0.00
−0.05
−0.10

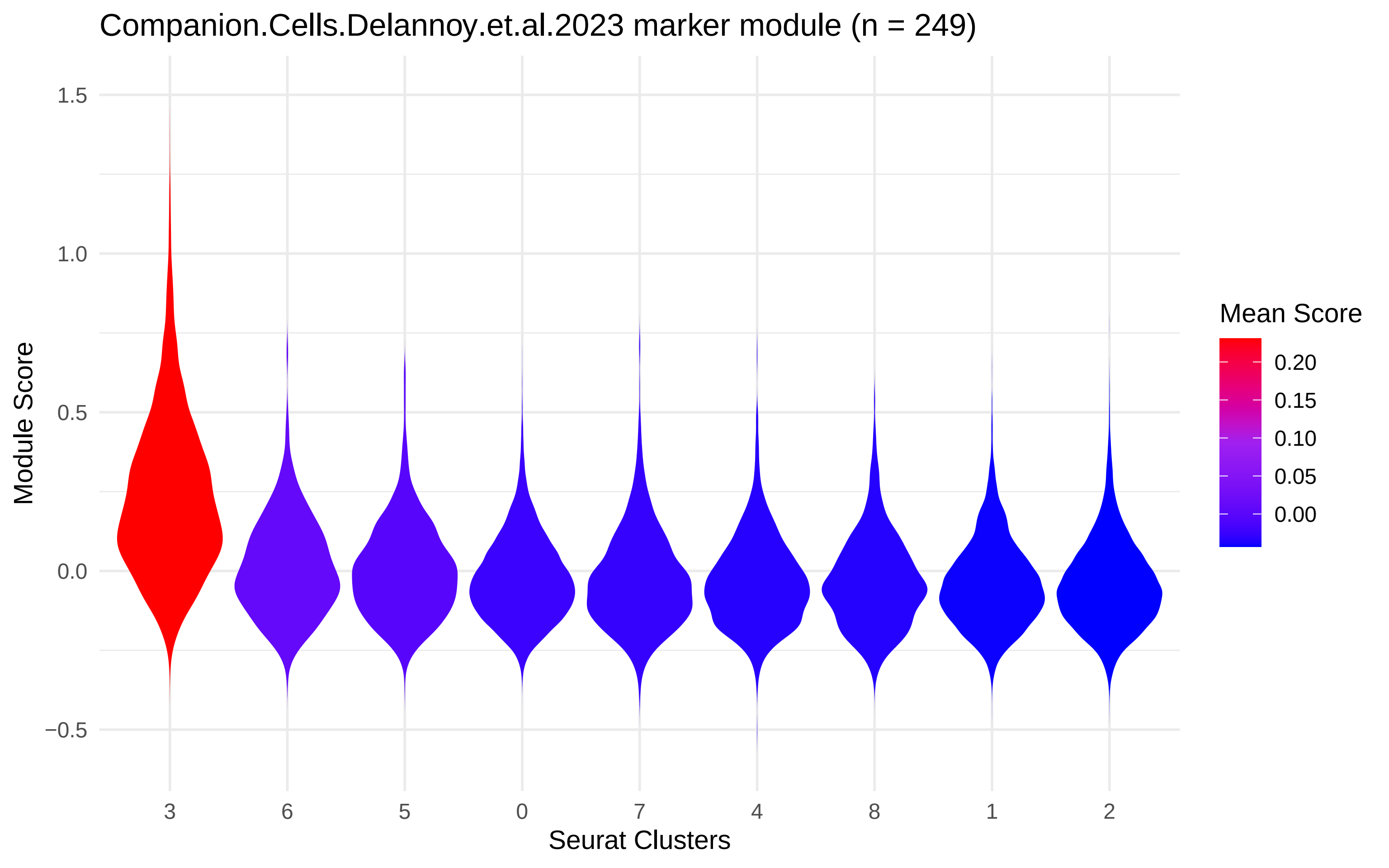

Companion.Cells.Delannoy.et.al.2023 marker module (n = 249)
Module Score
1.5
1.0
0.5
0.0
−0.5
3
6
5
0
7
4
8
1
2
Seurat Clusters
Mean Score
0.20
0.15
0.10
0.05
0.00

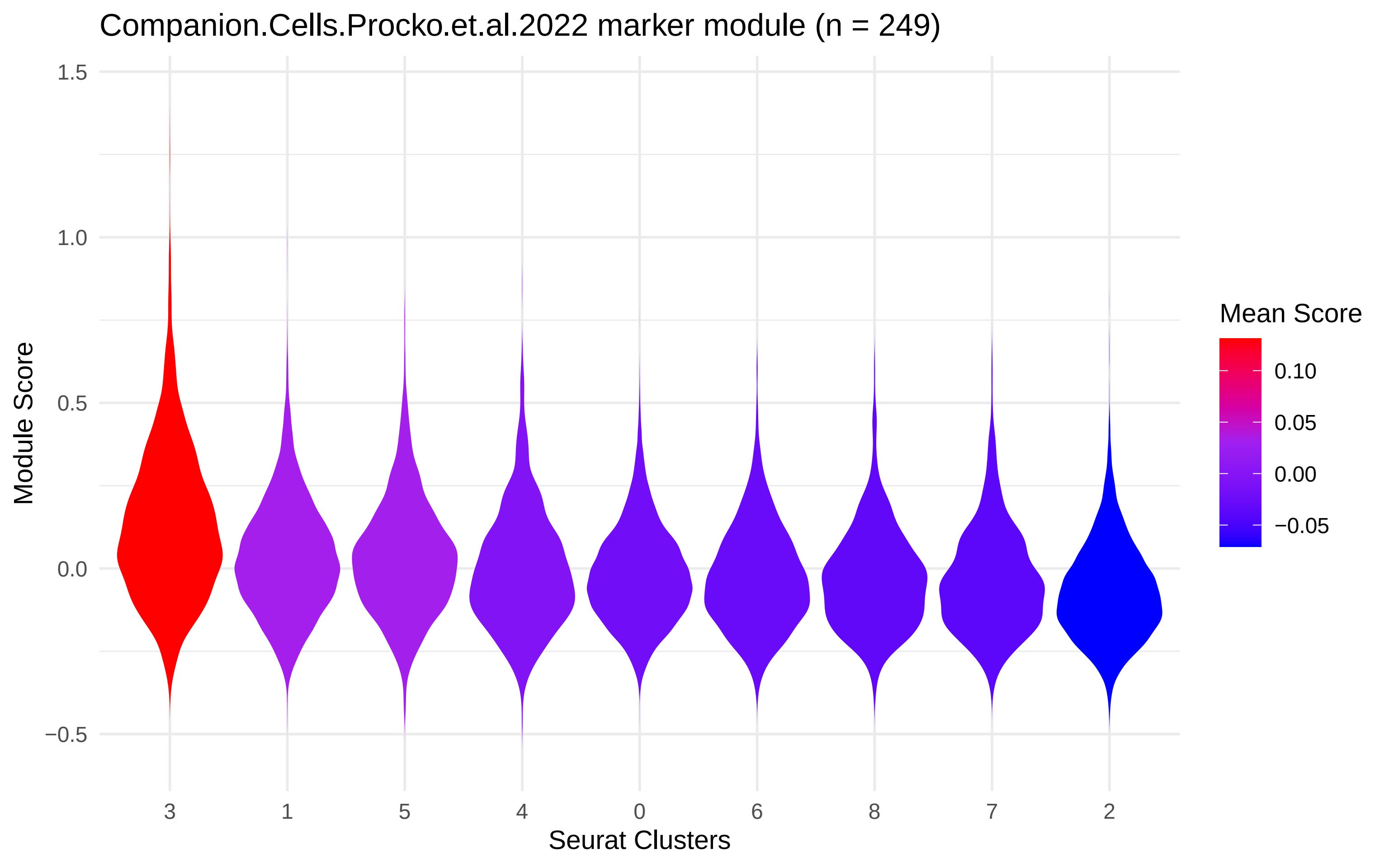
Companion.Cells.Procko.et.al.2022 marker module (n = 249)
Module Score
1.5
1.0
0.5
0.0
−0.5
3
1
5
4
0
6
8
7
2
Seurat Clusters
Mean Score
0.10
0.05
0.00
−0.05

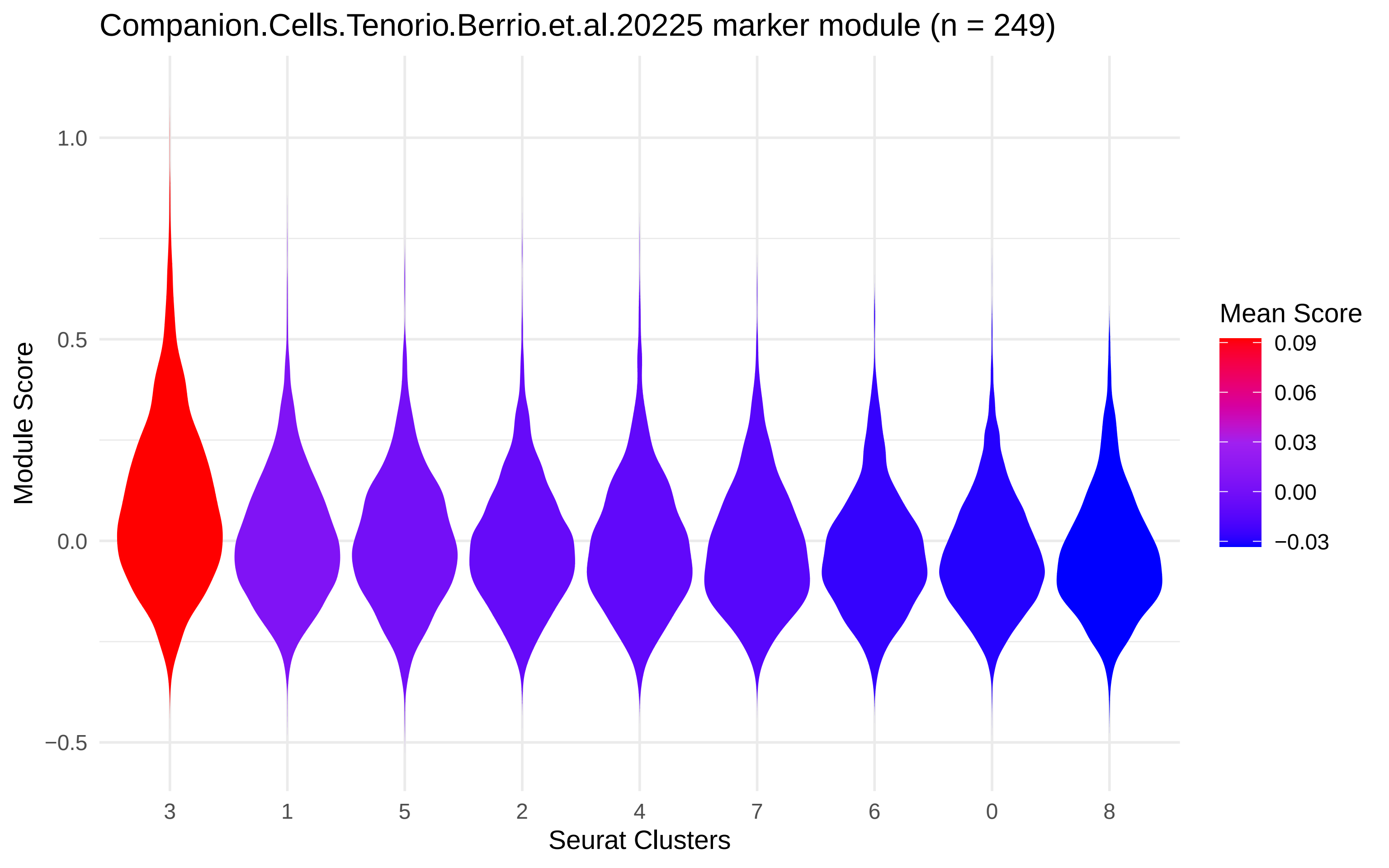
Companion.Cells.Tenorio.Berrio.et.al.20225 marker module (n = 249)
Module Score
1.0
0.5
0.0
−0.5
3
1
5
2
4
7
6
0
8
Seurat Clusters
Mean Score
0.09
0.06
0.03
0.00
−0.03

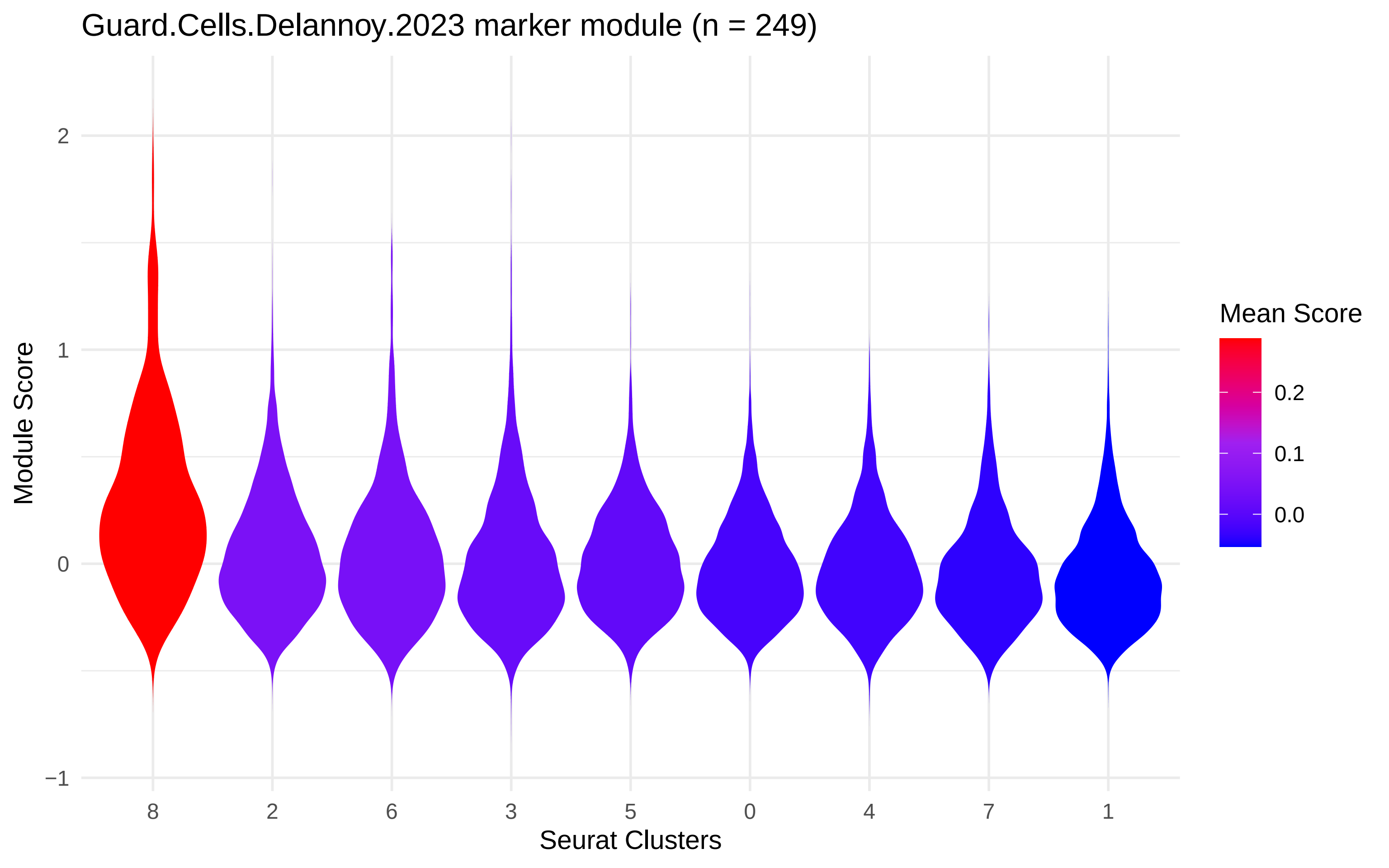
Guard.Cells.Delannoy.2023 marker module (n = 249)
Module Score
2
1
0
−1
8
2
6
3
5
0
4
7
1
Seurat Clusters
Mean Score
0.2
0.1
0.0

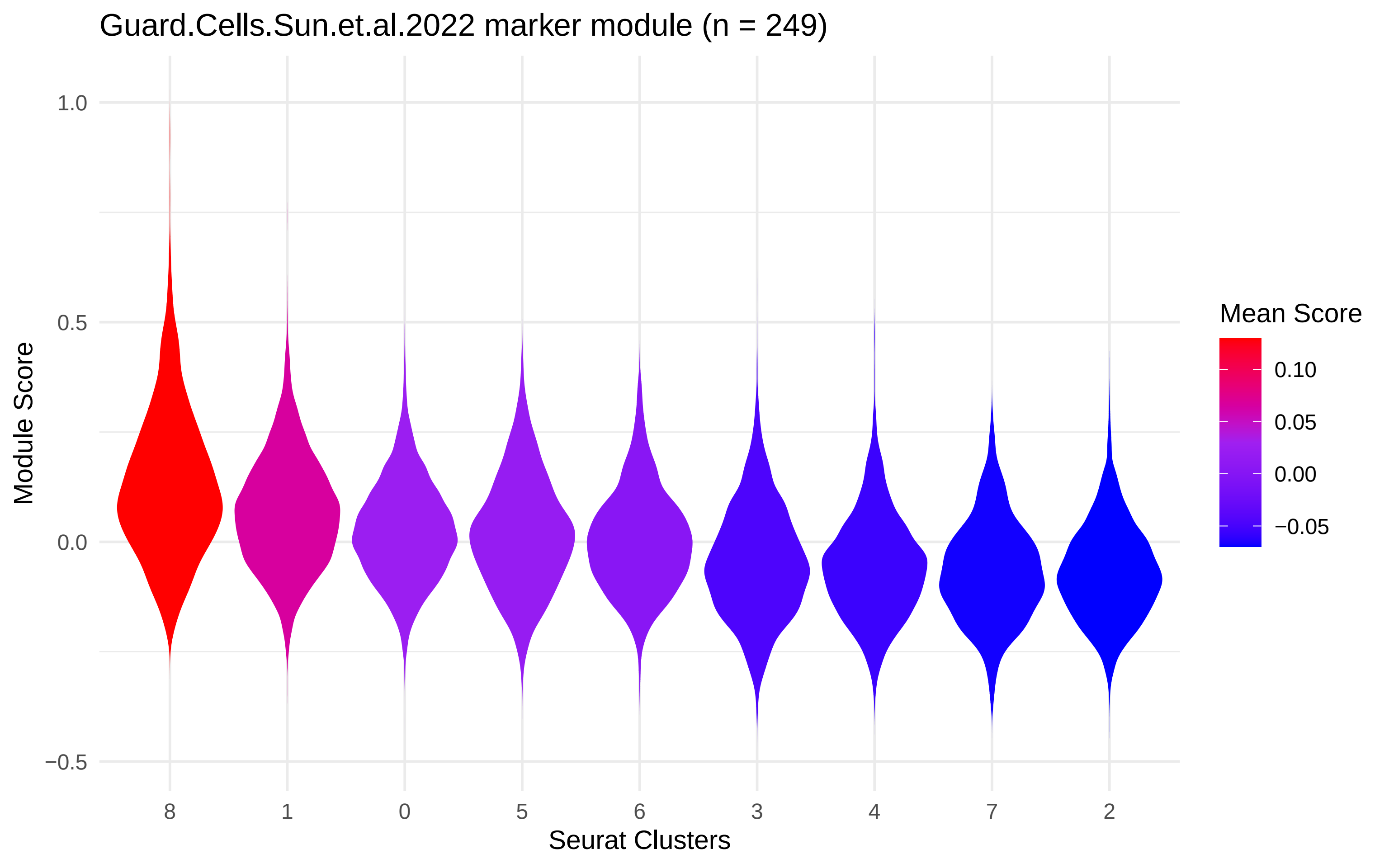
Guard.Cells.Sun.et.al.2022 marker module (n = 249)
Module Score
1.0
0.5
0.0
−0.5
8
1
0
5
6
3
4
7
2
Seurat Clusters
Mean Score
0.10
0.05
0.00
−0.05

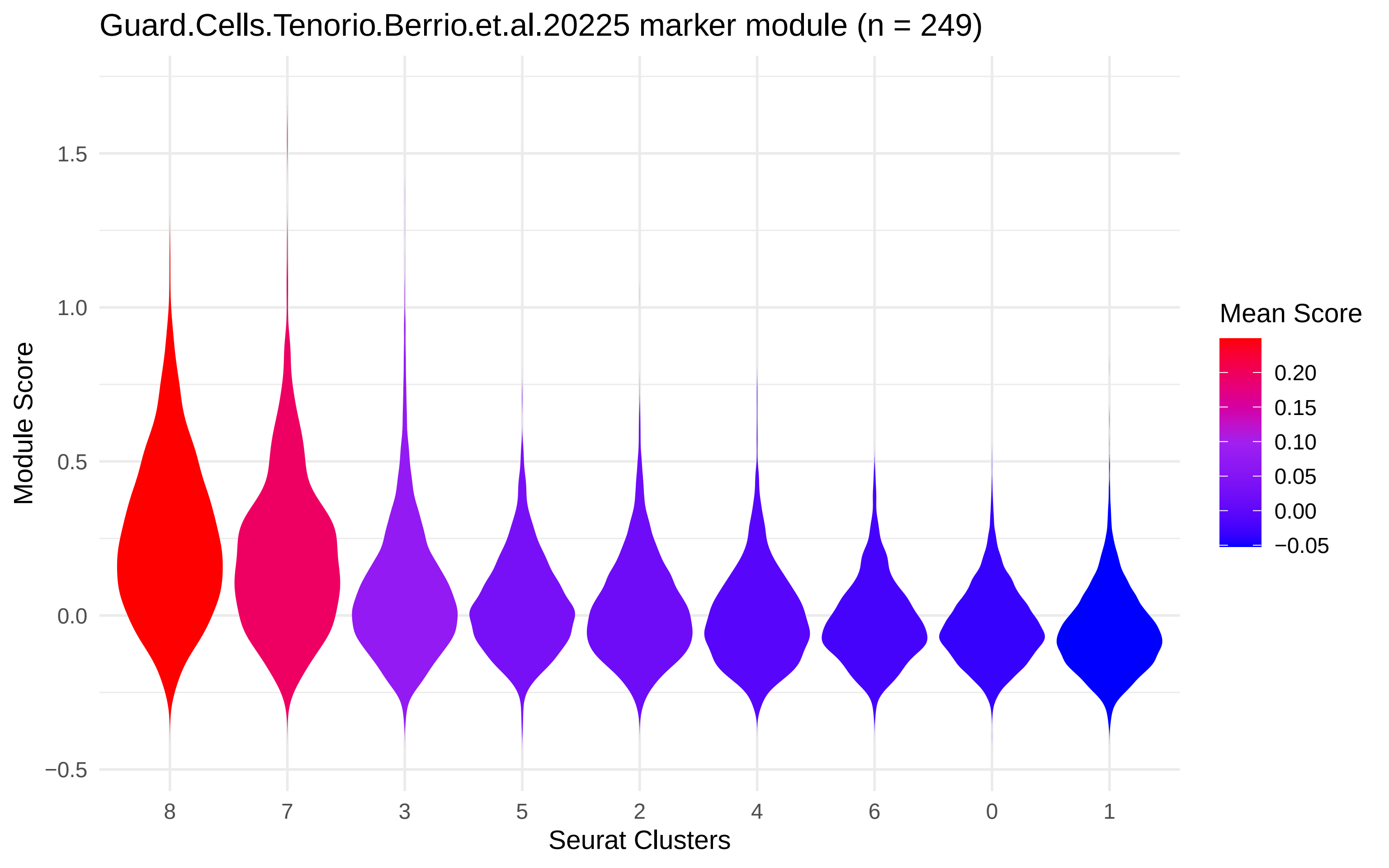

Guard.Cells.Tenorio.Berrio.et.al.20225 marker module (n = 249)
Module Score
1.5
1.0
0.5
0.0
−0.5
8
7
3
5
2
4
6
0
1
Seurat Clusters
Mean Score
0.20
0.15
0.10
0.05
0.00
−0.05

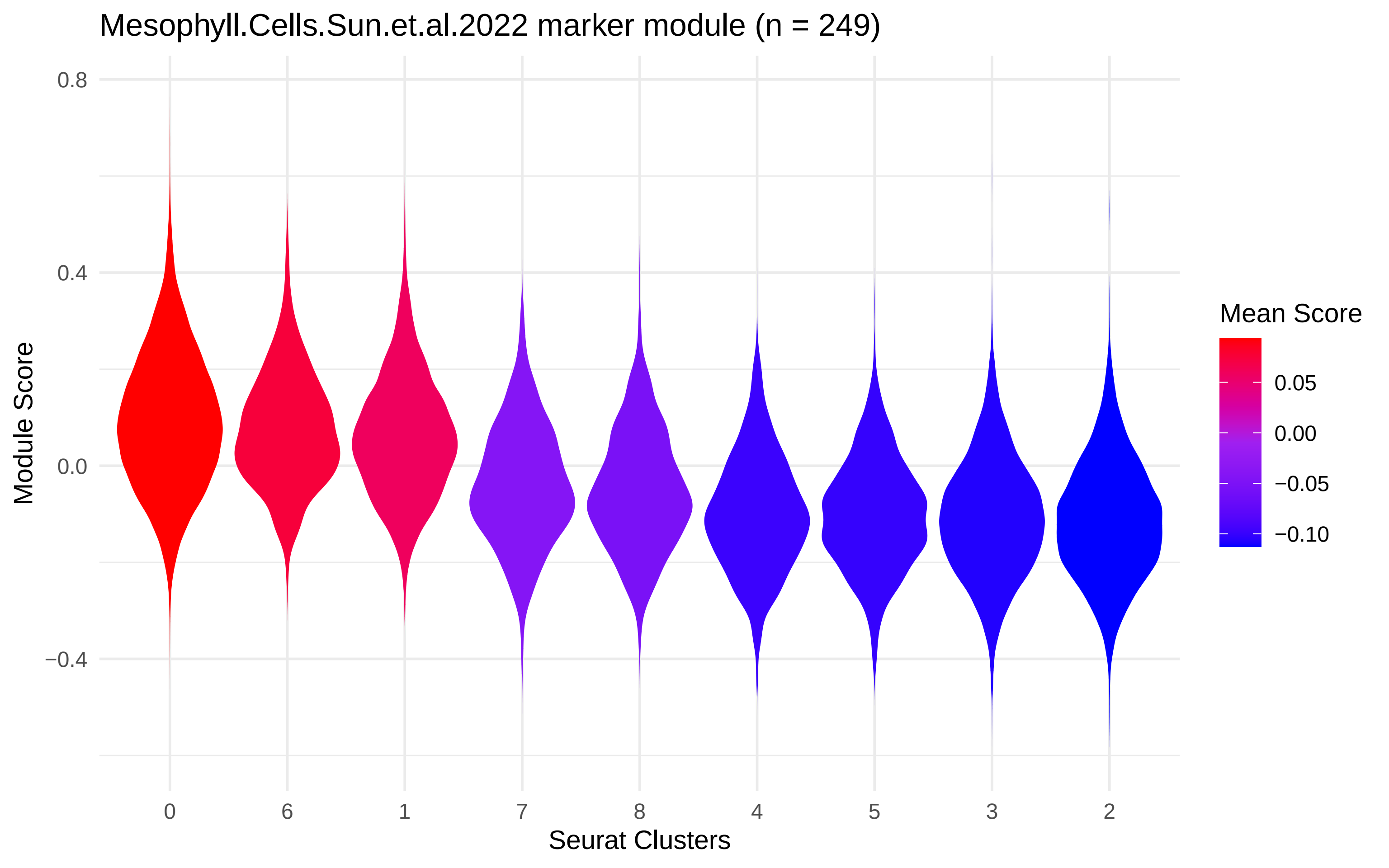

Mesophyll.Cells.Sun.et.al.2022 marker module (n = 249)
Module Score
0.8
0.4
0.0
−0.4
0
6
1
7
8
4
5
3
2
Seurat Clusters
Mean Score
0.05
0.00
−0.05
−0.10

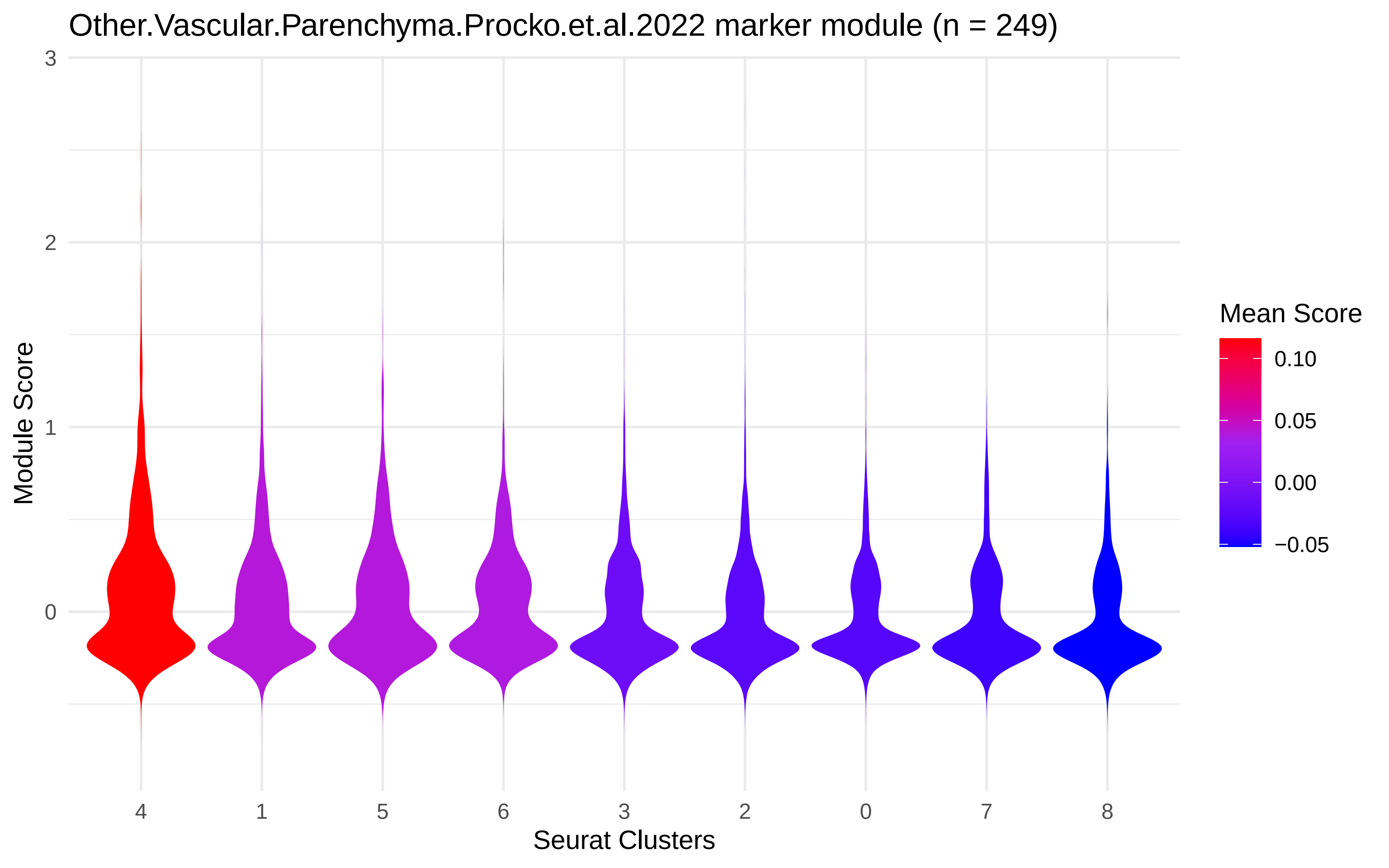
Other.Vascular.Parenchyma.Procko.et.al.2022 marker module (n = 249)
Module Score
3
2
1
0
Seurat Clusters
4
1
5
6
3
2
0
7
8
Mean Score
0.10
0.05
0.00
−0.05

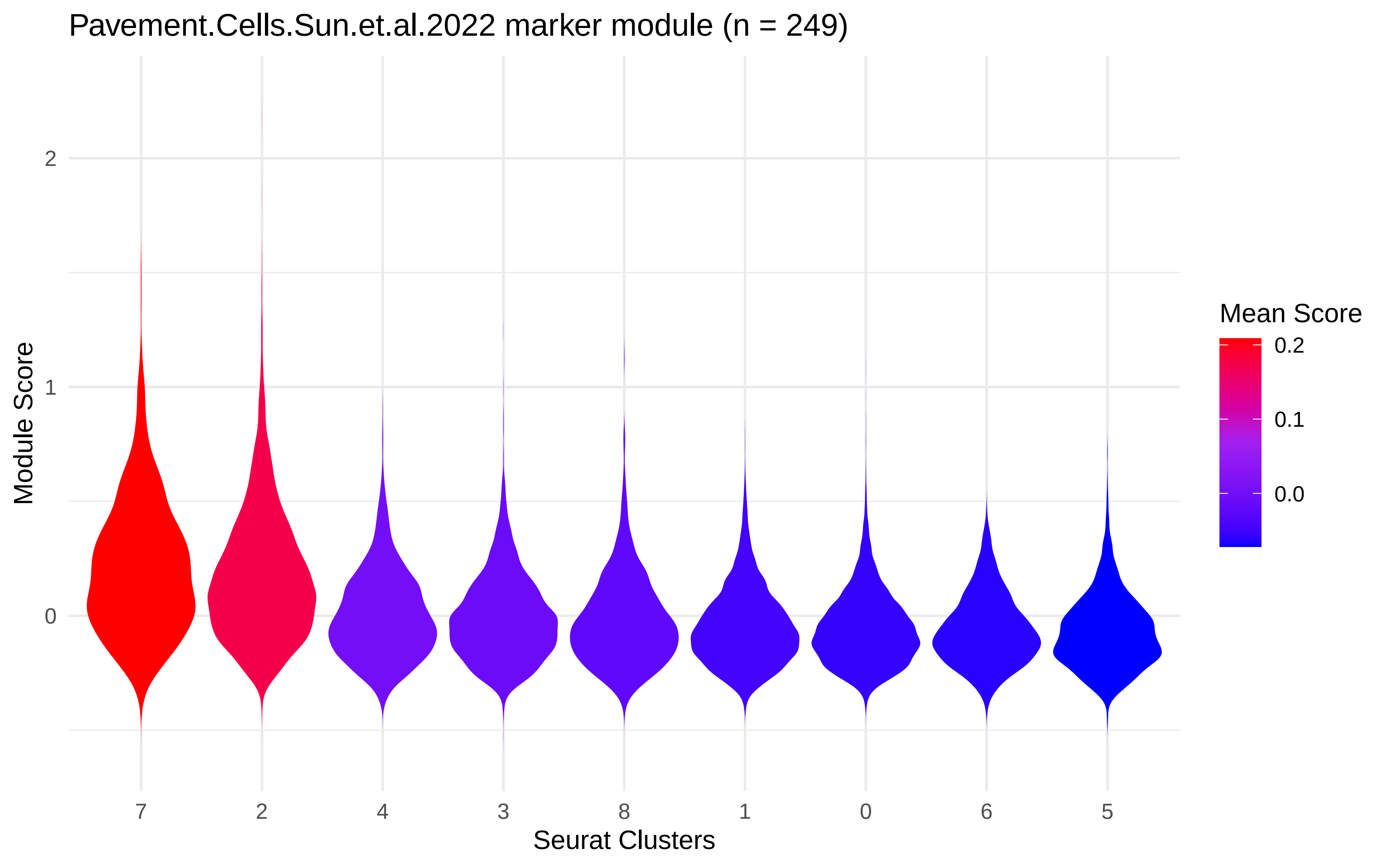
Pavement.Cells.Sun.et.al.2022 marker module (n = 249)
Module Score
2
1
0
7
2
4
3
8
1
0
6
5
Seurat Clusters
Mean Score
0.2
0.1
0.0

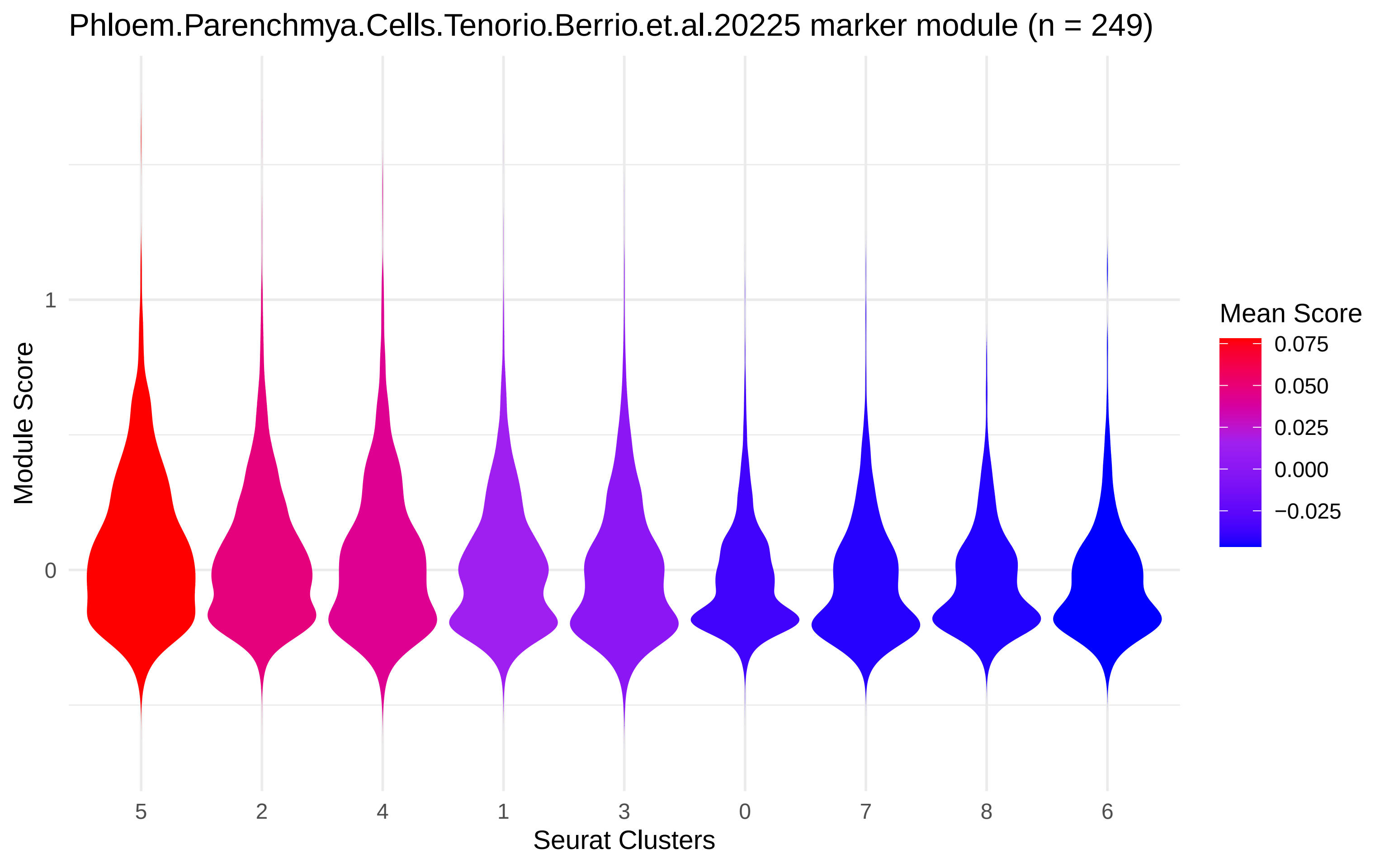
Phloem.Parenchmya.Cells.Tenorio.Berrio.et.al.20225 marker module (n = 249)
Module Score
1
0
5
2
4
1
3
0
7
8
6
Seurat Clusters
Mean Score
0.075
0.050
0.025
0.000
−0.025

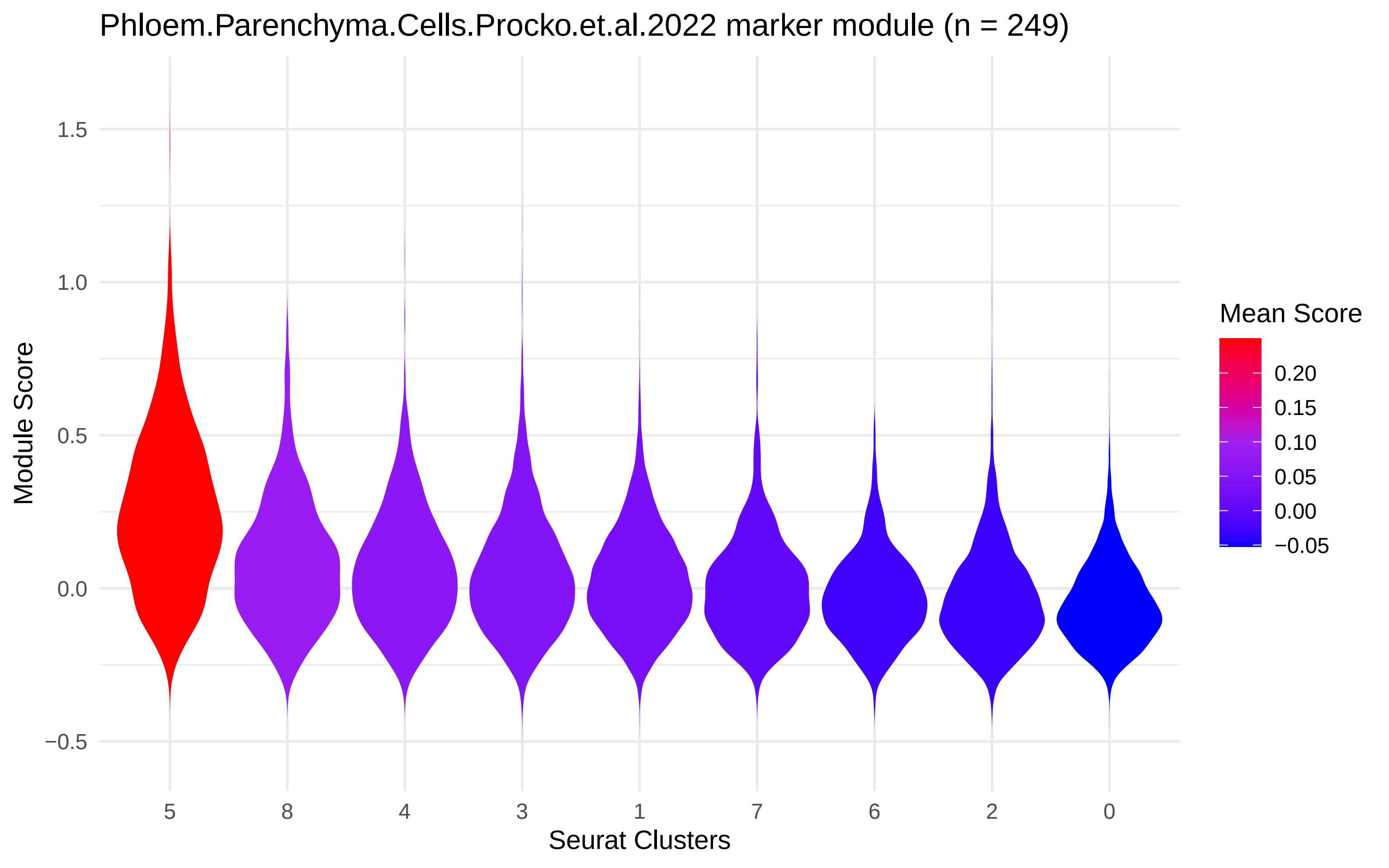

Phloem.Parenchyma.Cells.Procko.et.al.2022 marker module (n = 249)
Module Score
1.5
1.0
0.5
0.0
−0.5
5
8
4
3
1
7
6
2
0
Seurat Clusters
Mean Score
0.20
0.15
0.10
0.05
0.00
−0.05

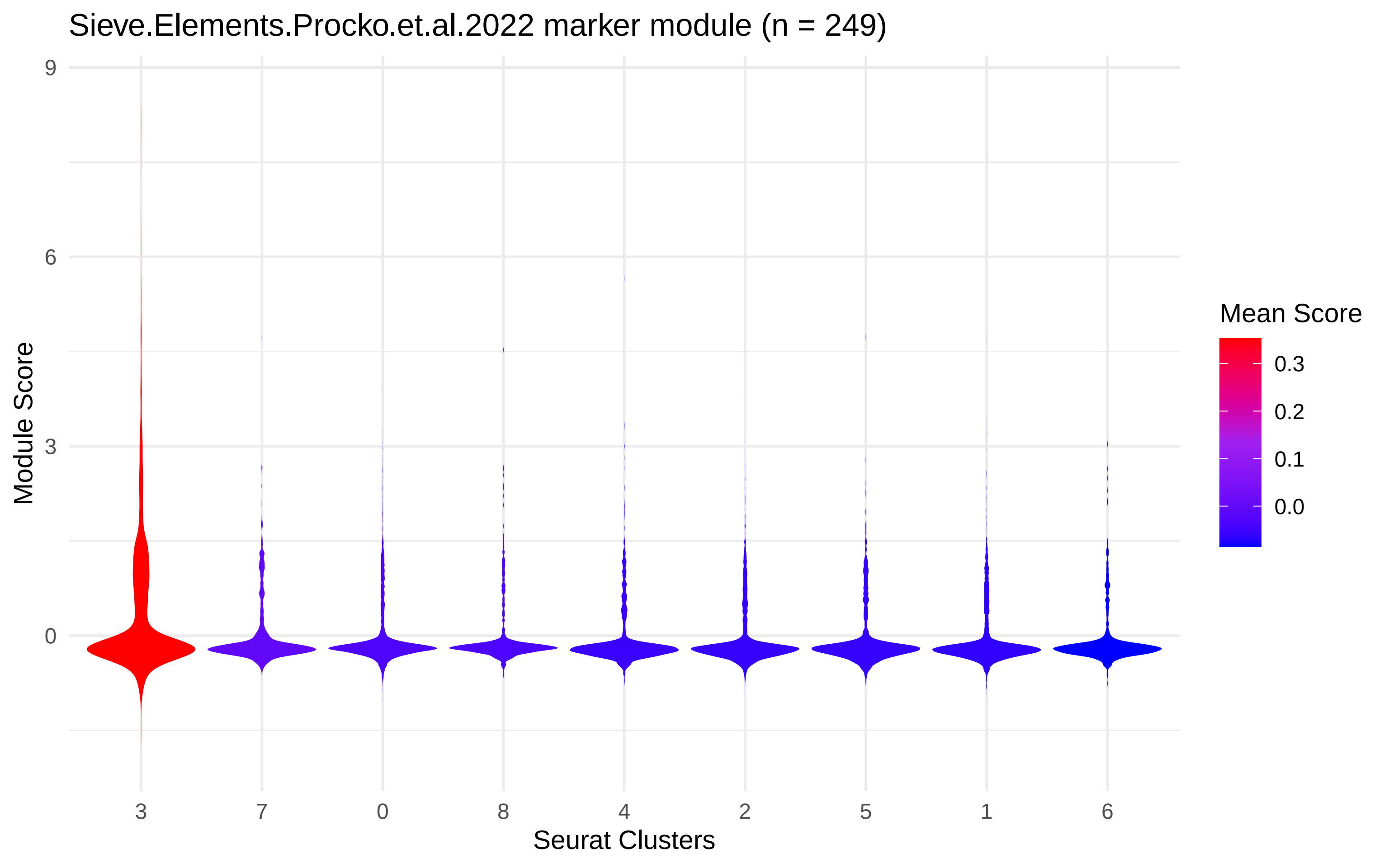
Sieve.Elements.Procko.et.al.2022 marker module (n = 249)
Module Score
9
6
3
0
3
7
0
8
4
2
5
1
6
Seurat Clusters
Mean Score
0.3
0.2
0.1
0.0

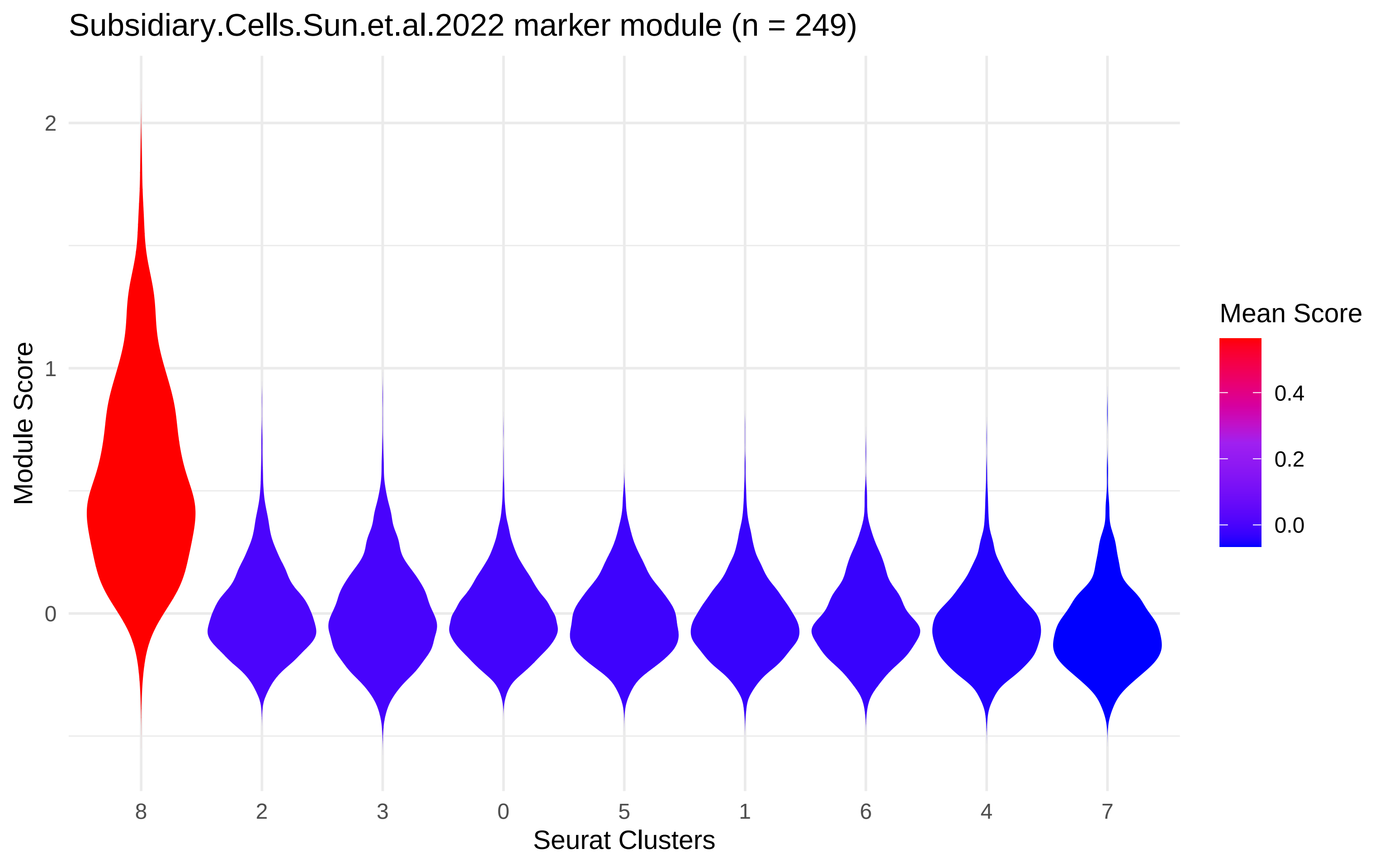

Subsidiary.Cells.Sun.et.al.2022 marker module (n = 249)
Module Score
2
1
0
8
2
3
0
5
1
6
4
7
Seurat Clusters
Mean Score
0.4
0.2
0.0

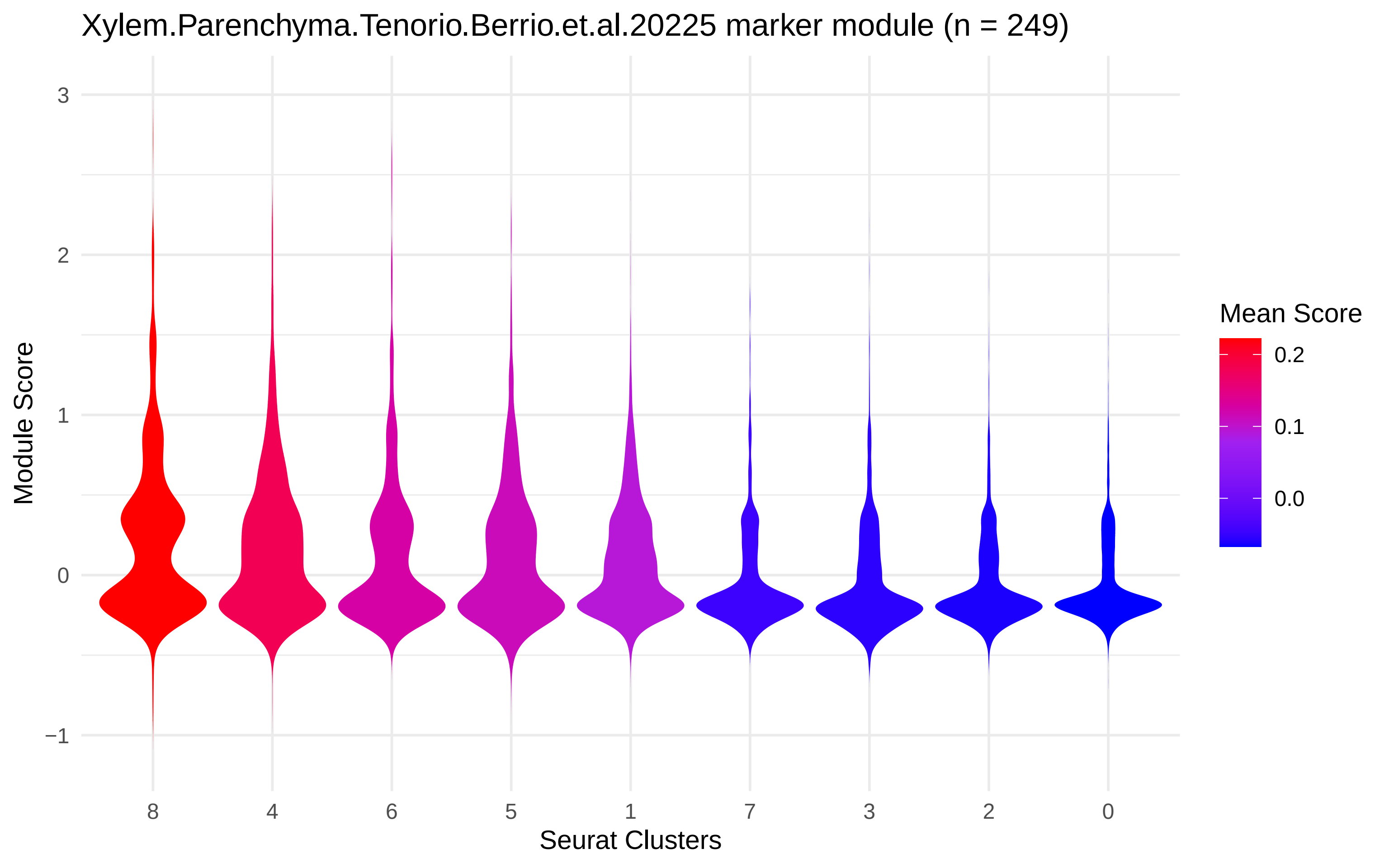

Xylem.Parenchyma.Tenorio.Berrio.et.al.20225 marker module (n = 249)
Module Score
3
2
1
0
−1
8
4
6
5
1
7
3
2
0
Seurat Clusters
Mean Score
0.2
0.1
0.0
